# Thalamo-insular basis for updating physiological needs

**DOI:** 10.64898/2026.10.20.760372

**Authors:** Yael Prilutski, Inbar Perets, Stav Shtiglitz, Katayun Cohen-Kashi Malina, Itay Talpir, Yoav Livneh

## Abstract

Physiological needs, such as thirst and hunger, produce powerful motivation for behaviors that are essential for survival. Yet, how the brain estimates these needs remains unknown. It is unclear whether the brain continuously monitors bodily signals to create real-time representations, or if existing internal representations are only updated when meaningful changes occur. To address this question, we manipulated thalamic internal sensory inputs to the interoceptive insular cortex. We found that inhibiting internal sensory inputs prevents the updating of insular cortex representations of thirst and hunger, thereby keeping its activity closer to its initial physiological need state. Consequently, cued feeding and drinking, as well as free consumption of food and water, persist beyond the current physiological need. We further show that insular cortex activity normally represents both predictions and sensory information. However, violating learned expectations evokes widespread need-specific prediction error signals. Our results thus reveal a thalamo-cortical basis for updating physiological needs in the brain.

---

Interoception, the sense of the physiological condition of the body, involves sensing and integrating various signals related to heart rate, blood pressure, blood sugar, gastric stretch, blood CO_2_ levels, temperature, inflammation, taste, and more^1,2^. Physiological needs, such as hunger and thirst, generate motivations for specific consummatory behaviors. Yet, it remains unclear how physiological need levels are estimated in the brain using interoceptive signals. There are two dominant views of how interoceptive signals form an internal model of the body^3–6^. Classic models propose that the brain represents bodily signals in real-time to maintain a dynamic representation of internal needs. However, more recent models arising from fields of active inference and predictive processing, propose that existing internally generated representations are updated when meaningful changes occur. Here, we set out to test these ideas.

Insular cortex (InsCtx) is the main cortical site that receives interoceptive sensory signals from brainstem and thalamic interoceptive sensory pathways^7–10^, and is thus considered the “primary interoceptive cortex”. InsCtx is thought to mediate the interoceptive aspects of numerous behaviors, from feeding and drinking, to emotions and social behaviors^11–19^. Nevertheless, the mechanistic role of InsCtx in interoception still remains unclear. Predictive processing models suggest that InsCtx combines interoceptive information with external sensory information to predict upcoming physiological changes, which guide need-driven behavior. These models suggest that when interoceptive predictions are violated, InsCtx computes interoceptive prediction error signals^3,4,8,20,21^. Consistent with these models, InsCtx responds to predictive cues across different sensory modalities^12–15,17,22–27^. Furthermore, InsCtx activity on longer timescales represents physiological need states such as hunger and thirst^13,28–30^. Nevertheless, there has been little direct mechanistic evidence for interoceptive prediction and prediction error signals in InsCtx. A deeper understanding of the role of InsCtx in interoception is essential for understanding its involvement in diverse pathological conditions, including eating disorders, obesity, anxiety, depression and drug addiction^3,6,31–34^.

Here, we set out to test the two models of interoception in the context of physiological needs. We inhibited incoming interoceptive inputs to InsCtx and examined how this would affect InsCtx activity and need-driven behavior. To do so, we combined a novel behavioral apparatus, longitudinal microprism-based two-photon imaging of InsCtx activity, and pathway-specific manipulations.

## Interoceptive thalamic inputs to InsCtx are essential for updating need-driven behavior

To directly examine how the brain tracks and updates physiological needs, we designed an experimental approach that allows simultaneous assessment of two internal need states, hunger and thirst, and their corresponding need-fulfilling behaviors - eating and drinking. We developed a novel behavioral apparatus that allowed head-fixed mice (for compatibility with two-photon imaging, see below) to consume food pellets and/or water drops during the same session (**Fig. 1a**). Our behavioral apparatus utilized a combination of infrared beam-breaks, capacitance touch sensors, and deep learning-based video analysis (DeepLabCut^35^) to precisely determine the timing of each lick, pellet delivery, and pellet consumption event (**Fig. 1a; Extended Data Video 1; Supp. Materials**).

**Figure 1:**
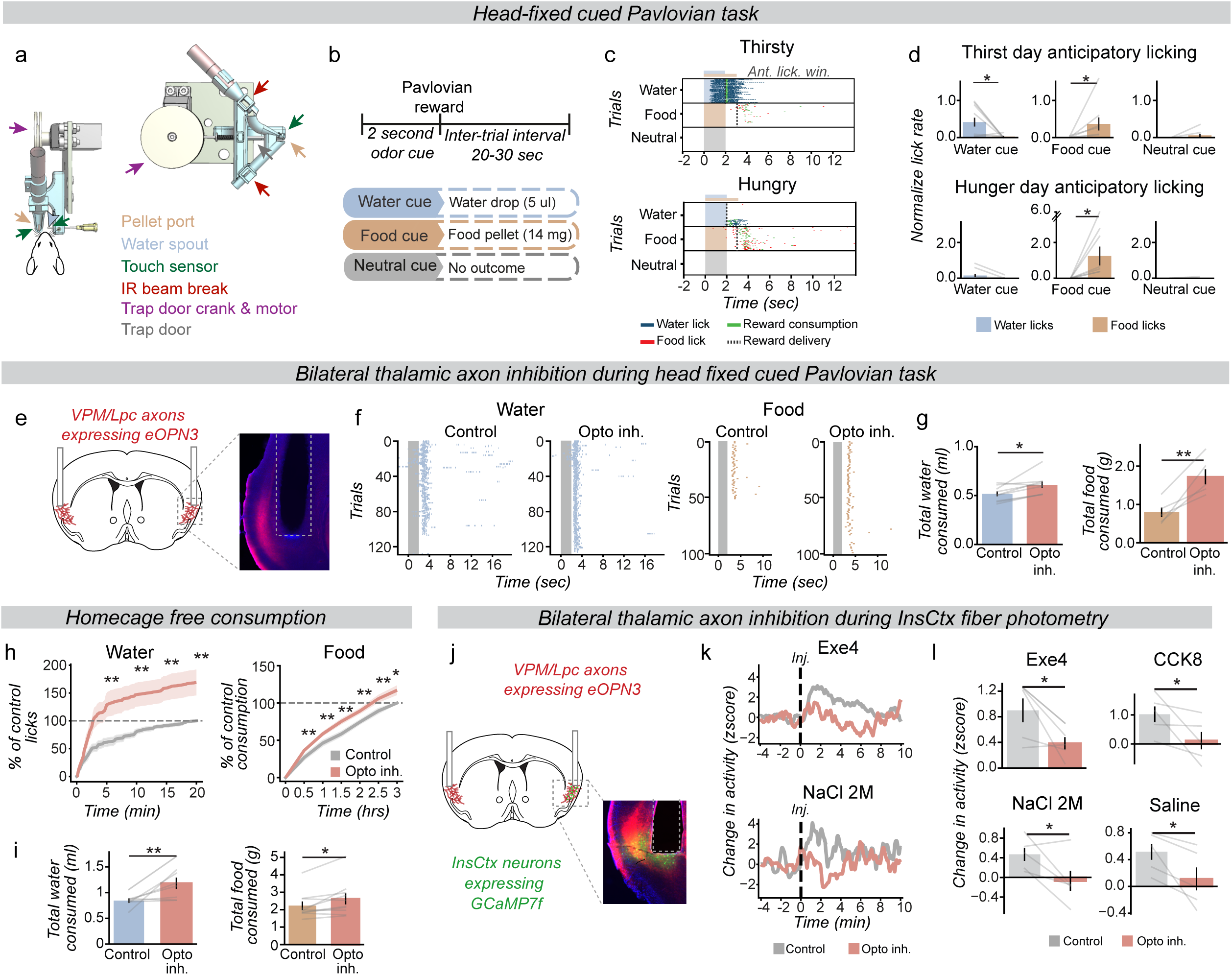
Interoceptive thalamic inputs to InsCtx are essential for updating need-driven behavior. a. Behavioral apparatus for cue-guided Pavlovian consumption of solid food pellets (14 mg) and drinking water (5 µl) in head-fixed mice (for compatibility with two-photon imaging). Pellet arrival from a pellet dispenser (not shown) is detected with an infrared (IR) beam break. If the pellet is not consumed, a “trap door” opens, allowing the pellet to drop and this is detected by another IR beam break. Licks are detected with capacitance sensors. Water is delivered via a blunt needle. See further details in Methods and Ext. Data Video 1. b. Behavior task trial structure: three different odor cues, followed by a Pavlovian food or water reward, or no outcome. Note that inter-trial interval durations are randomized and differ between different trial types. c. Example licking behavior from an example mouse for water (blue ticks) and food (red ticks) across trials during thirst (*top*) and hunger (*bottom*) state. Horizontal lines above the top raster marks the anticipatory licking window (blue-water trials, 2 sec; brown-food trials, 3 sec). Green ticks: reward consumption. Shaded rectangles indicate cue period. Dashed line: reward delivery (note different timing between water and food trials). d. Anticipatory licking to the water spout (blue) or to the food pellet port (brown) in food, water, and neutral cues (last second of the cue for water, last second of the cue and until reward delivery for food- ∼1 sec after cue offset). Values are the mean of the first 30 food/water trials. Responses were normalized to food or water licks rates during food or water restriction, respectively, when only one reward was available. (*Thirst:* water-trials *p=0.02, food-trials *p=0.048, neutral-trials p=0.11, *Hunger*: water-trials p=0.067, food-trials *p=0.027, neutral-trials p=0.17). n = 10 mice; mean±SEM). e. *Left*: Schematic of bilateral optogenetic inhibition of VPM/Lpc→InsCtx axons in InsCtx. *Right*: coronal brain section showing of VPM/Lpc→InsCtx axons in InsCtx expressing eOPN3-mScarlet with the optic fibers located above the InsCtx. f. Effects of bilateral optogenetic inhibition of VPM/Lpc→InsCtx axons on behavior during water restriction (*left*) and food restriction (*right*). Lick raster plots from an example mouse across a control day and an optogenetic inhibition day. g. Quantification of the effects of optogenetic inhibition on cue-guided feeding and drinking behavior. (*left*-water, *p=0.033, n=9; mean±SEM) and food (*right*, **p=0.004, n=5; mean±SEM). h. Effects of bilateral optogenetic inhibition of VPM/Lpc→InsCtx axons on home-cage free consumption. Values on control and optogenetic inhibition days were normalized to the total consumption on the control day. *Left*: water lick count in water-restricted mice (p = 0.004, 0.003, 0.004, 0.005, n = 8). *Right*: food consumption in mice food-restricted overnight, with water freely available (p = 0.001, 0.002, 0.002, 0.005, 0.007, 0.025, n = 10; mean±SEM). i. Quantification of total consumption of ‘h’. *Left*-water consumption, **p=0.009, n = 8. *Right*-food consumption, *p=0.012, n = 10; mean±SEM) j. *Left*: Schematic of bilateral optogenetic inhibition of VPM/Lpc→InsCtx axons (red) with GCaMP fiber photometry (green) in the InsCtx. *Right*: coronal brain section showing of VPM/Lpc→InsCtx axons in InsCtx expressing eOPN3-mScarlet and InsCtx neurons expressing GCaMP7f with the optic fiber track (dashed rectangle). k. Example GCaMP7f fiber photometry recording from an example mouse during Exendin-4 injection (*top*) and NaCl injection (*bottom*), under control conditions (gray) and bilateral optogenetic inhibition of VPM/Lpc→InsCtx axons (red). Dashed line indicates injection timing. l. Effects of bilateral optogenetic inhibition of VPM/Lpc→InsCtx axons on InsCtx activity during manipulation of interoceptive signals. *Top*: hunger-related signals in overnight food-restricted mice (Exendin-4: *p = 0.035; CCK8: *p = 0.039). *Bottom*: thirst-related signals in overnight water-restricted mice (saline: *p = 0.022) and non-restricted mice (NaCl: *p = 0.032). n = 5; mean±SEM.

We trained mice to associate different olfactory cues with Pavlovian delivery of a water drop (5 µL), delivery of a food pellet (14 mg, 3.6 kcal/gram), or no outcome (neutral cue; **Fig. 1b**). Mice readily learned the task, as evidenced by cue-specific, need-specific, anticipatory licking (from cue onset to reward delivery: 2 sec for water and 3 sec for food; **Fig. 1c-d**; **Extended Data Fig. 1a**). We designed this simple behavioral task as it allowed mice to freely choose whether or not to consume food or water, predicted by learned cues. Importantly, while most studies in food-restricted head-fixed mice use liquid rewards (e.g., Ensure milkshake, condensed milk, sugar water), we chose to use dry food pellets so that each reward will satisfy only one need, but not the other (i.e., liquid food like Ensure would satisfy both the need for fluids and energy). Mice readily performed this task either food-restricted or water-restricted. We note that we use the terms thirst/hunger and water/food-restriction interchangeably for brevity and clarity.

We aimed to inhibit incoming interoceptive inputs to InsCtx and examine the resulting behavioral effects. To do so, we focused on the main source of interoceptive input to InsCtx, the sensory visceral and gustatory thalamic nuclei (parvicellular portion of the ventroposteromedial and ventroposterolateral thalamus, VPMpc and VPLpc). These two adjacent nuclei are the primary source of visceral and gustatory sensory information to InsCtx^36–44^. Because VPMpc and VPLpc have not been functionally delineated in vivo (e.g., VPMpc, initially thought to be gustatory, also appears to have other functions^45,46^), and because they represent an anatomical medio-lateral continuum, we refer to them jointly here as VPM/Lpc. We specifically inhibited VPM/Lpc→InsCtx inputs using the inhibitory opsin, eOPN3^47^, which has been shown to efficiently inhibit thalamo-cortical axon terminals (**Fig. 1e, Extended Data Fig. 1b**; 620 nm, 15 Hz, 30% duty cycle, 9 mW). This way, we could specifically inhibit VPM/Lpc→InsCtx inputs without affecting VPM/Lpc inputs to other brain regions such as central amygdala.

We examined how bilateral VPM/Lpc→InsCtx optogenetic inhibition affected need-driven behavior either in food-restricted or water-restricted mice performing the behavioral task described above. We allowed each animal to consume food or water until it voluntarily stopped consumption. We compared behavior in control days and days with inhibition of VPM/Lpc→InsCtx inputs. During the optogenetic inhibition sessions, the illumination started after a short baseline period in order to establish the baseline behavior for that day. Strikingly, inhibition of VPM/Lpc→InsCtx inputs increased the amount of water consumed in water-restricted mice, and of food consumed in food-restricted mice (**Fig. 1f-g, Extended Data Fig. 1c**). We further tested whether the bilateral VPM/Lpc→InsCtx optogenetic inhibition would affect other behavioral parameters indicative of higher consummatory motivation. We examined its effects on anticipatory licking, lick latency, and consummatory licking. Interestingly, we only observed a consistent significant effect across mice for consummatory lick rate (**Extended Data Fig. 1d**). Importantly, we did not observe any behavioral effects in mice with unilateral illumination (see **Fig. 2d** and **Extended Data Fig. 3o-u** below).

**Figure 2:**
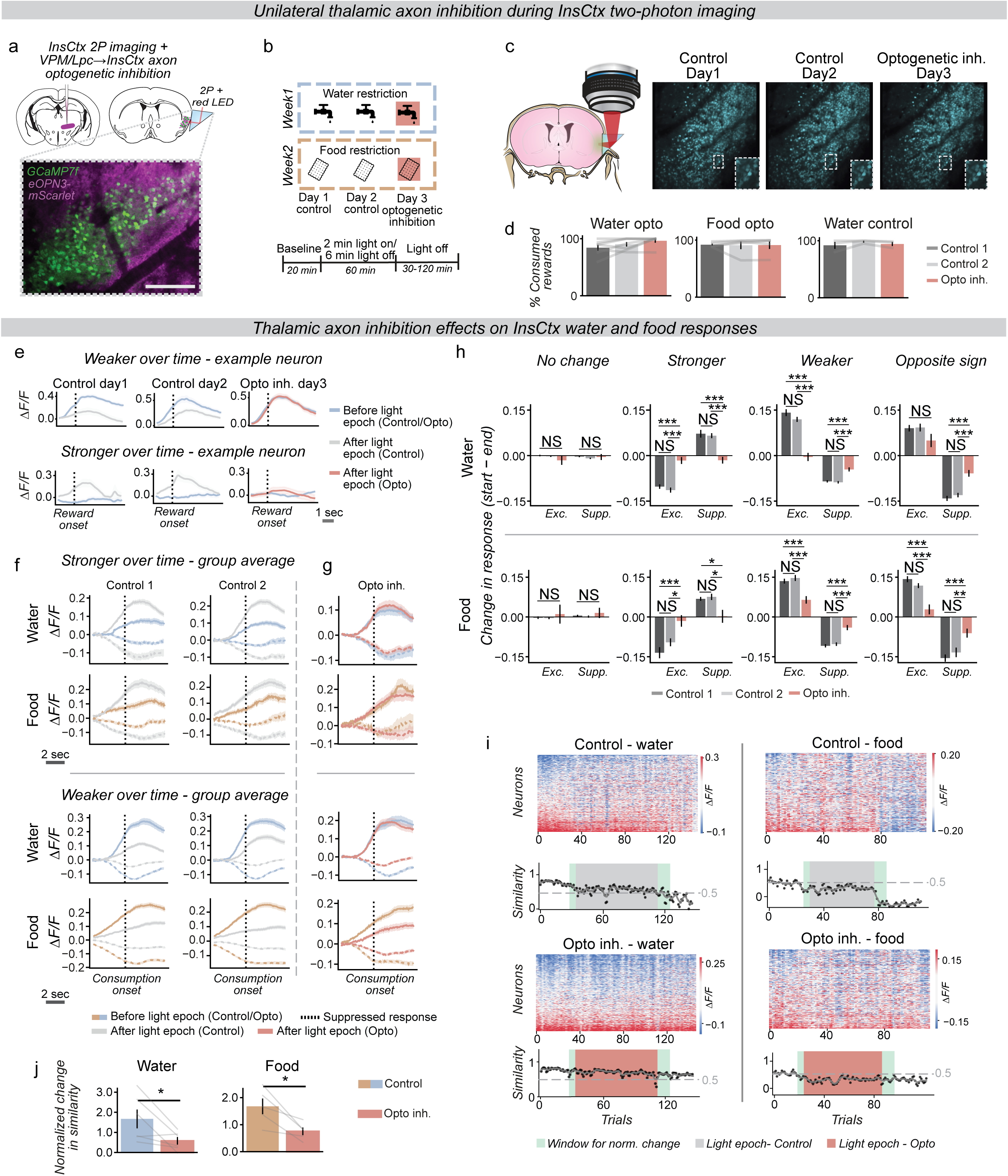
Interoceptive thalamic inputs are essential for updating representations of food and water in InsCtx. a. Approach for microprism-based two-photon (2P) imaging InsCtx activity while inhibiting VPM/Lpc→InsCtx axons. *Top*: schematic coronal brain section and approach. *Bottom*: Two-photon image of InsCtx neurons expressing GCaMP7f (green), and eOPN3-mScarlet VPM/Lpc axons (magenta). Scale bar: 250 µm. b. Experimental timeline of optogenetic inhibition. Two control days, followed by a day of optogenetic inhibition, in water-restricted mice for water and neutral trials, or in food-restricted mice for food and neutral trials. Each session lasted 2 hours. Inhibition started after 20 minutes, with 8 cycles of 2-minute light-on / 6-minute light-off. c. Time-lapse microprism-based two-photon imaging of InsCtx. *Left*: schematic coronal brain section illustrating the microprism approach. *Right*: Example two-photon field-of-view in each of 3 days of the experiment (722 neurons tracked across the 3 days). d. Percent consumed rewards during the first 2 hours of a session, across 3 days. Water consumption during water restriction with optogenetic inhibition in eOPN-expressing mice (*left*), food consumption during food restriction with optogenetic inhibition in eOPN-expressing mice (*middle*), and water consumption during water restriction for control mice without eOPN3 expression (*right*). No significant change in consumption in any group. e. Single-neuron examples of the long-term effect of optogenetic inhibition of VPM/Lpc→InsCtx axons on water responses. *Top*: example neuron showing an initially strong response that becomes weaker as the physiological need decreases. *Bottom*: example neuron showing an initially weak response that becomes stronger as the physiological need decreases. Pink: responses after the optogenetic inhibition epoch. Gray: responses during the same time period on control days without illumination. Note the stabilization of responses following optogenetic inhibition of VPM/Lpc→InsCtx axons, i.e., prevention of the changes associated with the decrease in physiological need. f. Change in InsCtx average group responses with a decrease in physiological need in two control days (Control 1, Control 2), pooled across all mice. Only responsive cells showing a stable change across both control days were included, grouped by whether their responses became stronger (*top*) or weaker (*bottom*) over time. Solid lines: excitatory responses, dashed lines: suppressed responses. Blue (water responses in water restricted mice. n = 6 mice, stronger: 108 neurons, weaker: 235 neurons) and brown (food responses in food restricted mice. n = 5 mice, stronger: 41 neurons, weaker: 176 neurons): responses ∼25 min into the session. Gray: responses after 1 hour of consumption. Dotted vertical line: consumption onset. g. Same cells as in ‘f’, but during the day with optogenetic inhibition of VPM/Lpc→InsCtx axons. Blue (water) and brown (food): responses ∼25 min into the session, before optogenetic inhibition. Red: responses after optogenetic inhibition. Solid lines: excitatory responses, dashed lines: suppressed responses. Dotted vertical line: consumption onset. Note the stabilization of responses following optogenetic inhibition of VPM/Lpc→InsCtx axons, i.e., prevention of the changes associated with the decrease in physiological need. h. Quantification of the change in individual neuron response magnitude for each cell type, pooled across all mice (water: n = 6 mice, no change-78 neurons, stronger-108 neurons, weaker-235 neurons, opposite sign-108 neurons, food: n = 5 mice, no change-27 neurons, stronger-41 neurons, weaker-176 neurons, opposite sign-66 neurons; mean±SEM). For each cell, response was measured as the mean of the 2 seconds following reward consumption onset, and the delta between start and end of the session was computed for each of the three days (Control 1 - C1, Control 2 - C2, Opto). Cells were grouped into four categories based on their natural change with decreasing physiological need in control days: No change, Stronger, Weaker, and Opposite sign, shown separately for excited (Exc.) and suppressed (supp.) responses. Note that in all groups with a natural change in control days, optogenetic inhibition attenuates or completely eliminates this change. *Top*: water responses. **No change**-no significant changes. **Stronger**: Exc.-C1 vs. opto: ***p<0.001, C2 vs. opto: ***p<0.001, Supp.- C1 vs. opto: ***p<0.001, C2 vs. opto: ***p<0.001. **Weaker**: Exc.- C1 vs. opto: ***p=<0.001, C2 vs. opto: ***p<0.001, Supp.- C1 vs. opto: ***p<0.001, C2 vs. opto: ***p<0.001. **Opposite sign**: Exc.- no significant changes, Supp.- C1 vs. opto: ***p<0.001, C2 vs. opto: ***p<0.001). *Bottom*: food responses. **No change**-no significant changes. **Stronger**: Exc. C1 vs. opto: ***p<0.001, C2 vs. opto: *p=0.012, Supp.- C1 vs. opto: *p=0.015, C2 vs. opto: *p=0.015. **Weaker**: Exc.- C1 vs. opto: ***p=<0.001, C2 vs. opto: ***p<0.001, Supp.- C1 vs. opto: ***p<0.001, C2 vs. opto: ***p<0.001. **Opposite sign**: Ecx.- C1 vs. opto: ***p<0.001, C2 vs. opto: ***p<0.001, Supp.- C1 vs. opto: ***p<0.001, C2 vs. opto: **p=0.004). No significant difference was found between Control 1 and Control 2 in any group. i. Example mouse showing that optogenetic inhibition attenuates the gradual change in InsCtx population responses to water and food, associated with the decrease in physiological need. Heatmaps show the mean reward response across neurons for each consumption trial on a control or optogenetic inhibition day. Line plots below each heatmap show the population response similarity (cosine similarity, normalized to baseline) for the data in that heatmap. Gray/pink rectangles: optogenetic inhibition epoch in control (gray) and optogenetic inhibition (pink) days. Green rectangle: time window used for calculating change in similarity in ‘j’. j. Summary of the change in population response similarity before and after the illumination period, normalized by Control Day 2 to control for day-to-day variability. Note significantly lower changes during the optogenetic day for water (*p=0.021, n = 6 mice) and food (*p=0.012, n = 5 mice), indicating an overall stabilization of responses, attenuating the natural effects of the decrease in physiological need. 406±192 and 325±131 neurons per mouse, for water and food, respectively; mean±SEM.

To test whether these effects were specific to the head-fixed cued behavioral task, we also examined home-cage consumption in the absence of any structured task. We observed similar effects in water-restricted mice during home-cage free consumption of water, and in food-restricted mice during home-cage free consumption of food (also with access to water to avoid dry food inducing thirst, i.e., dehydration induced anorexia^48^; **Fig. 1h-i, Extended Data Fig. 1e**). Interestingly, when we optogenetically inhibited VPM/Lpc→InsCtx inputs in ad libitum fed mice, we did not observe an increase in consumption (**Extended Data Fig. 1f**). Together, these results suggest that inhibition of VPM/Lpc→InsCtx inputs prevented behavioral changes associated with decreased hunger or thirst, indicating that these inputs may carry signals critical for updating need-driven behavior.

Given that VPM/Lpc→InsCtx optogenetic inhibition affected both feeding in hungry mice and drinking in thirsty mice, we sought to test its functional role in relaying hunger- and thirst-related interoceptive information. To do so, we used fiber photometry recordings of bulk InsCtx calcium activity with/without eOPN3-mediated inhibition (**Fig. 1j, Extended Data Fig. 1g**). In food-restricted mice we found that VPM/Lpc→InsCtx optogenetic inhibition attenuated the responses of InsCtx to injection of the GLP1 agonist, Exendin-4^49^, and of cholecystokinin (CCK-8; **Fig. 1k top, l top**). In water-restricted mice we found that VPM/Lpc→InsCtx optogenetic inhibition attenuated the responses of InsCtx to a rehydrating injection of isotonic saline, and in quenched mice the responses to injection of hypertonic NaCl that induces osmotic thirst (**Fig. 1k bottom, l bottom**). We note that we performed these experiments in anesthetized mice to isolate feedforward sensory signals from effects of expectation on InsCtx activity. Importantly, optogenetic inhibition did not affect activity in sham injections (**Extended Data Fig. 1h).** Together, these results are consistent with VPM/Lpc→InsCtx inhibition preventing the updating of need-driven behavior by interoceptive inputs. We next examined how changes in InsCtx activity underlie these behavioral effects.

## Interoceptive thalamic inputs are essential for updating representations of food and water in InsCtx

To examine the effects of VPM/Lpc→InsCtx inhibition on InsCtx activity independently of effects on behavior, we unilaterally expressed eOPN3 in VPM/Lpc neurons (**Fig. 2a; Extended Data Fig. 2a**). We imaged layer 2/3 GCaMP7f-expressing neurons in mid-posterior InsCtx via a microprism^14^ while simultaneously optogenetically inhibiting VPM/Lpc→InsCtx axons (**Fig. 2a; Extended Data Fig. 2b**). We note that this approach targets layer 2/3 anatomically, but does not have genetic specificity to this specific cortical layer and may thus include neurons from other cortical layers.

Following our fiber photometry experiments above (**Fig. 1j-l**), we also verified the efficacy of eOPN3-mediated inhibition of VPM/Lpc→InsCtx axons through the microprism by examining the effects of axon terminal illumination on ongoing InsCtx activity in the absence of any overt behavior (i.e., “quiet waking”; **Extended Data Fig. 2c**). In agreement with previous pharmacological inhibition of VPM/Lpc somata^41^ and with optogenetic activation of VPM/Lpc axons in InsCtx slices ex vivo^50,51^, we found that optogenetic inhibition decreased ongoing activity in a subset of neurons, while increasing activity in others. Overall, the ongoing activity of 15±2% of neurons was significantly affected during the first 2 min of optogenetic illumination in mice not engaged in an overt behavioral task. Notably, this is likely a considerable underestimation of the efficacy of inhibition as most InsCtx ongoing activity in the absence of any visceral or physiological change likely does not rely on inputs outside InsCtx^52,53^. Importantly, we did not observe such effects over the same period of time in the absence of illumination (**Extended Data Fig. 2c**). We note that while optogenetic illumination via an optic fiber (**Fig. 1**) will produce multi-layer suppression of axons in InsCtx, red light delivered through the prism (which penetrates brain tissue relatively well) will be somewhat attenuated as it reaches deeper layers ( attenuation to ∼60%, ∼5.4 mW, at a depth of 500 µm, estimated using Monte Carlo simulations^54^), but this should still be sufficient to efficiently activate eOPN3 in deep cortical layers^47,55^. As such, our measurements of InsCtx layer 2/3 dynamics do not only reflect direct inhibition of thalamic inputs to layer 2/3, but also network effects that span multiple layers.

We next examined how VPM/Lpc→InsCtx inhibition would affect activity in InsCtx, comparing two control days (no-light) with the optogenetic inhibition day (**Fig. 2b-c**). In our protocol, each session started with a 20 min baseline period, followed by 8 cycles of 2 minutes with the light on, and 6 minutes with the light off. After this ∼ 1-hr min illumination period ended, we continued imaging until mice voluntarily stopped consumption. Notably, unilateral optogenetic inhibition did not cause any overt changes in behavior, and did not affect the rate of food or water consumption (**Fig. 2d**).

Seeing as bilateral VPM/Lpc→InsCtx inhibition prevented the decrease in feeding or drinking associated with fulfilling the need for food or water (**Fig. 1**), we first examined how unilateral inhibition affected InsCtx responses to food and water consumption. Addressing this question required across-day comparisons. We therefore first examined the stability of InsCtx responses to the same need-fulfilling reward within the same physiological need across days, and found that they were quite stable (**Extended Data Fig. 2d-f**).

Next, we examined how InsCtx responses to need-relevant rewards change following a decrease in physiological need. To assess the effect of declining thirst on water responses, we compared consumption responses in water-restricted mice between the start of the session (high thirst) and the end of the session, after consumption of dozens of water rewards (reaching a quenched state). To assess the effect of declining hunger on food responses, we compared consumption responses in food-restricted mice between the start of the session (high hunger) and a later session on the same day, after ∼4 hrs of ad libitum access to food and water in the home-cage (reaching satiety). We used this approach as food pellet consumption could induce thirst. We note that for VPM/Lpc→InsCtx optogenetic inhibition experiments, we compared food responses within the same session as we sought to determine its effects on continuous changes in InsCtx activity during consumption.

We first examined individual neuronal food/water responses and found that many responses changed, either decreasing or increasing after prolonged consumption (**Fig. 2e, left and middle; Extended Data Fig. 2g**). Across the population, 68.1% and 79.6% of neurons significantly responded to food or water responses across the two control days, respectively. Of these responsive neurons, 56% of neurons consistently changed during the day as mice consumed food/water across two days (for both food and water, **Extended Data Fig. 2h**). Similar to the effects on behavior described above (**Fig. 1**), optogenetic inhibition of VPM/Lpc→InsCtx inputs appeared to block the change in single neuron responses associated with decreased hunger/thirst (**Fig. 2e, right**). For quantitative analysis of the effects of VPM/Lpc→InsCtx inhibition we applied a stringent criterion, focusing only on neurons with highly consistent changes days in both direction and magnitude, associated with reduced physiological need (**Fig. 2f; Extended Data Fig. 2i-j**). We found that the responses of neurons whose activity changed after prolonged consumption (i.e., increased, decreased, or changed sign) were stabilized by optogenetic inhibition of VPM/Lpc→InsCtx inputs, as evident by significantly smaller within-session changes (**Fig. 2f-h, Extended Data Fig. 2k-m**).

At the neuronal population level, we compared the changes associated with reduced physiological need within-day (e.g., thirsty-quenched population vector similarity) to the similarity across two days with similar need levels (e.g., thirsty-thirsty population vector similarity). We found the within-day similarity to be substantially lower than between the beginnings of different sessions across days (**Extended Data Fig. 2n-o**). Importantly, these changes were not explained by changes in different motor parameters such as lick latency, lick bout structure, lick rate, anticipatory licking rate, chewing-related orofacial movements, pellet reaching/grasping/handling, as well as by pupil size as a proxy for arousal levels **(Extended Data Fig. 3a-i)**. Here as well, VPM/Lpc→InsCtx inhibition prevented the decrease in population vector similarity across trials, associated with prolonged consumption and reduced physiological need, but only for water (**Extended Data Fig. 3j-k**).

To further characterize the continuous evolution of these changes as physiological need gradually decreases, we calculated the normalized cosine similarity of population food/water responses across trials. For each trial, we computed the mean food/water response across neurons, and compared this trial vector to the mean of the first 15 trials. We observed a clear, progressive decrease in cosine similarity as the session proceeded, for both food and water trials (**Fig. 2i top**), indicating that neuronal population responses became increasingly dissimilar from their initial representation with continued consumption. When examining the effect of VPM/Lpc→InsCtx optogenetic inhibition, we found that it attenuated the decrease in similarity that naturally occurred with reduced physiological need (**Fig. 2i bottom, Fig. 2j**). We note that these gradual changes during the session likely reflect decreased hunger/thirst levels, and not other variables associated with time in session (e.g., elapsed time or fatigue), as we did not observe them in water-restricted mice when both dry food and water were available (**Extended Data Fig. 3l**). In this scenario, mice did not reach water satiety within the same time period, likely because water consumption relieved dehydration-induced anorexia^48^, which allowed feeding on dry pellets, which in turn caused additional thirst.

Thus, inhibition of VPM/Lpc→InsCtx inputs appears to stabilize food/water responses in InsCtx neurons, attenuating the effects of the gradual decrease in hunger or thirst. These effects were not observed in control mice not expressing eOPN3 (**Extended Data Fig. 3m-n**). Importantly, consistent with the lack of effect on consummatory behavior of unilateral optogenetic inhibition, we did not observe differences in different motor variables between optogenetic inhibition and control conditions (lick latency, lick bout structure, lick rate, anticipatory licking rate, chewing-related orofacial movements, pellet reaching/grasping/handling), as well as by pupil size as a proxy for arousal levels (**Extended Data Fig. 3o-u**).

The changes for food-restricted mice with access only to food pellets could be attributed to decreased hunger drive or to a gradual increase in thirst and therefore dehydration-induced anorexia^48^. To test this, we compared the change in InsCtx food responses in food-restricted mice across two hunger days: one with only food available, and one with both food and water available. We first compared the change in normalized cosine similarity of food responses across each session, and observed similar changes on both days (**Extended Data Fig. 3v**). Second, we compared population food-response vectors at three time-points: the end of the day with both food and water, at the end of the food-only day, and after ∼4hrs free access to food and water following the food-only session. We examined the similarity between each pair of these population vectors (Pearson correlation). The population responses vectors were similar across all these time-points (**Extended Data Fig. 3w**). These results suggest that the changes we observed are likely due to decreased hunger.

Previous work has shown that pharmacological inhibition of VPM/Lpc somata attenuated taste responses in some InsCtx neurons, but not in others^41^. As such, we also assessed the immediate short-term effects of optogenetic inhibition on water/food responses, comparing them just before and just after the first optogenetic inhibition epoch (**Extended Data Fig. 4a**). As described above, we focused on neurons with changes across the same time-window that were consistent across days. We found that the responses of neurons whose activity changed within this short time-window were also stabilized by optogenetic inhibition, as evident by significantly smaller within-session changes. We also observed a population of neurons whose responses were initially stable (“no change”) but were changed by optogenetic inhibition, only for water (**Extended Data Fig. 4b-d**). Neurons stabilized or destabilized by optogenetic inhibition overlapped significantly above chance on short and long timescales. However, neurons destabilized on short timescales did not overlap with neurons stabilized on long timescales, indicating that optogenetic inhibition affects distinct neuronal populations differently. (**Extended Data Fig. 4e**). When examining activity at the population level, we found a decrease in the correlation slope between before and after optogenetic inhibition (6/7 mice during thirst, and 5/5 mice during hunger, although not statistically significant for food; **Extended Data Fig. 4f-g**), as well as when examining change in cosine similarity (but again, only for water; **Extended Data Fig. 4h**). This decrease likely reflects the fact that the destabilization effect is more apparent early in the session, when the effects of the reduction in the physiological need has still not fully developed. These effects were not observed in control mice not expressing eOPN3 (**Extended Data Fig. 4i-j**).

## Interoceptive thalamic inputs are essential for updating physiological need representations in InsCtx

We have previously shown that slow changes in ongoing InsCtx population activity patterns represent slow changes in thirst or hunger levels, independent of behavior or of artificial activation of hypothalamic “hunger neurons” and “thirst neurons”^13^. We therefore next examined how optogenetic inhibition of VPM/Lpc→InsCtx axons affected the representation of different physiological needs in InsCtx. We used our previously established method to assess changes in ongoing activity levels, reflecting the current physiological need state^8,13,29^. Briefly, we used inter-trial intervals (ITIs) only following neutral trials, when no overt behavioral change occurred^13^ (**Fig 3a**; see Methods). Consistent with our previous work^13^, we observed slow changes in ongoing activity of individual InsCtx neurons (**Fig 3b**).

**Figure 3:**
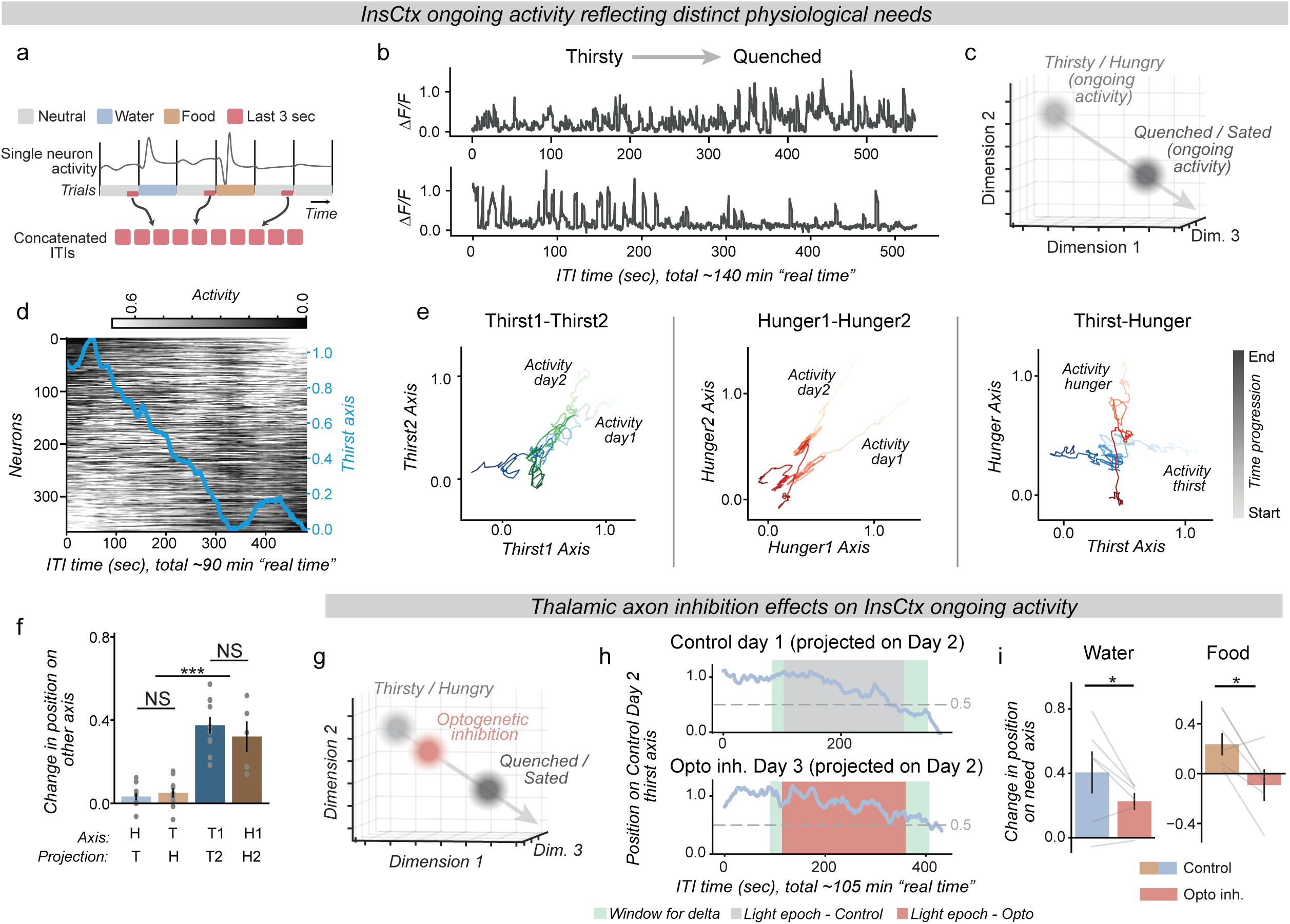
Interoceptive thalamic inputs are essential for updating physiological needs in InsCtx. a. Schematic of the approach for empirical determination of the physiological need state axes in population activity. The last 3 sec of neutral trial inter-trial interval (ITI) activity is extracted, concatenated, and renormalized (referred to as ‘ongoing activity’, following previously established procedures, see Methods). b. Single-neuron examples of changes in ongoing activity in water-restricted mice as the mouse consumes rewards and reaches a quenched state. *Top*: example neuron showing ongoing activity that increases between thirsty and quenched states. *Bottom*: example neuron showing ongoing activity that decreases between thirsty and quenched states. c. Schematic of the approach for empirical determination of the physiological need state axes in population activity space. Population ongoing activity in ITIs during distinct physiological need states is used to define the axis between them (thirsty vs. quenched or hungry vs. sated). d. Example ongoing activity of all neurons in one water-restricted mouse over the course of a session. Each row represents one neuron. Blue line indicates position along the thirst axis (1 = thirsty, 0 = quenched). n = 369 neurons. e. Example of ongoing activity movement along thirst and hunger axes. Comparison of two days of thirst state (*left*), two days of hunger state (*middle*), and one day of thirst vs. one day of hunger (*right*). 2D plots of ongoing activity from each day projected onto the axes of both days. A value of 1 indicates the hunger/thirst state, while a value of 0 indicates the quenched/sated state. Light-to-dark color changes represent time elapsed (light color: start, dark color: end). Note that ongoing activity in the right panel (thirst vs. hunger) shows minimal movement of ongoing activity in each need along the other need’s axis. f. Summary of physiological need state axis specificity. Change in ongoing activity position along the different day axes. Activity projected onto axes from a different day with the same physiological need state (e.g., thirst-thirst (T-T) or hunger-hunger (H-H)) moves significantly more along the other day’s axis than when comparing different physiological need state axes (thirst-hunger, T-H). All comparisons ***p < 0.001, n = 10 mice for T-T (381±174 neurons per mouse), n = 5 mice for H-H (325±131 neurons per mouse; mean±SEM), and n = 10 mice for T-H (349±175 neurons per mouse; mean±SEM). g. Same schematic approach as in ‘c’, now also showing the position along the thirst/hunger axis during unilateral optogenetic inhibition of VPM/Lpc→InsCtx axons (red), relative to natural change (dark gray), quantified in ‘h’ and ‘I’. h. Example of ongoing population activity in thirst state projected onto the Control Day 2 axis, for Control Day 1 (*top*) and the optogenetic inhibition day (Day 3, *bottom*). Rectangles indicate the optogenetic inhibition epoch (pink), or during same time-period in control days without illumination (gray). Note that optogenetic inhibition prevents the full transition to the quenched state. Green rectangles: time window used for calculating the change in position on axis in ‘i’. i. Summary of changes in position along the Control Day 2 axis, between the beginning and the end of the illumination period, comparing Control Day 1 and the optogenetic inhibition Day 3. Note significantly lower change on the optogenetic day for thirst, *p = 0.046, and for hunger, *p = 0.048. Thirst: n = 5 (414±209 neurons per mouse; mean±SEM). Hunger: n = 5 (325±131 neurons per mouse; mean±SEM).

To asses slow changes in activity at the population level, we projected population activity onto an axis defined between ongoing activity associated with physiological need (hungry or thirsty) versus sated or quenched states^13^ (**Fig 3c-d**). Similar to analyses of consummatory responses above (**Fig. 2**), we empirically defined these two need axes using days when mice were water-restricted and had access only to water, or food-restricted with access only to food. Specifically, for the thirst axis we used the beginning and the end of the session, after mice voluntarily stopped consuming water^13^. For the hunger axis we used the beginning of the session and the subsequent session on the same day after ∼4 hrs with ad libitum access to food and water in the home-cage. We note that the change over time in neurons’ ongoing activity was not correlated with their reward responses (**Ext. Data Fig. 5a).**

Seeing as this experiment required comparing physiological need axes across different days, we projected activity from pairs of days with either matching physiological needs (two food-restricted or two water-restricted days) or different needs (**Fig. 3e; Extended Data 5b**). We quantified their similarity by measuring the change in position along each axis across days (from values close to 1, indicating stability, to values closer to 0; **Fig. 3f**). Across matching need-day pairs, activity remained relatively aligned, indicating relatively stable representations of hunger and thirst. Across different-need day comparison, activity moved mostly along one axis but not the other, indicating distinct and somewhat independent representation of hunger and thirst, consistent with a low correlation between the hunger and thirst axes (**Extended Data Fig. 5c**). Similar to our previous work^13^, these changes likely reflected a decreased hunger/thirst levels, rather than other variables associated with time in session (e.g., elapsed time or fatigue), as we did not observe them in water-restricted mice when both dry food and water were available and mice did not reach water satiety within the same time period (**Extended Data Fig. 5d**). Comparing food-restricted mice across days with access to food alone or food and water revealed no significant differences, suggesting that water consumption does not affect representation of hunger (**Extended Data Fig. 5e**).

Using this approach, we next asked whether these physiological need representations reflect continuous incoming interoceptive sensory inputs, or perhaps an estimation that is occasionally updated. We predicted that if VPM/Lpc→InsCtx inputs provide continuous sensory input to form a sensory representation, then their inhibition should deteriorate or even eliminate this cortical representation. If this were the case, we would expect the projected values to potentially become erratic, fluctuate along the axis beyond the typical 0-1 range, and lack any clear gradual change. In contrast, if VPM/Lpc→InsCtx inputs serve to update an estimation, then their inhibition should prevent updating and maintain the representation close to the previous state regardless of the true current physiological need state.

We note that for optogenetic inhibition days, the need axis cannot be accurately determined as activity at the end of the day might be affected by the optogenetic inhibition. Therefore, to circumvent this issue we performed an across-day comparison during the 3-day experiment (days 1 and 2 were control days, and day 3 was the optogenetic inhibition day). We used the axis from day 2 and projected onto it the activity from day 1 (control day) and day 3 (optogenetic inhibition day). This allowed us to test how the need axis is affected by optogenetic inhibition, while controlling for across-day variability in InsCtx activity.

Remarkably, in agreement with updating, we found that inhibition of VPM/Lpc→InsCtx inputs prevented the gradual change of ongoing activity from activity patterns associated with a decrease in hunger or thirst (**Fig. 3g-i**). This is consistent with our behavioral findings (**Fig. 1**), where the same manipulation done bilaterally prevented the updating of need-driven behavior. We also verified that the effects of optogenetic inhibition did not stem from effects on ongoing arousal levels, comparing pupil size during ITIs (**Extended Data Fig. 5f**).

Our results suggests that VPM/Lpc→InsCtx optogenetic inhibition prevented network updating by thalamic inputs while potentially still maintaining the activity within a typical network state. However, it is still possible that optogenetic inhibition drove the network to an abnormal state (“off manifold manipulation”^56^). To test this, we used our previously established approach for characterizing the InsCtx population activity manifold (i.e., the repertoire of population activity patterns) and dynamics within it^57^. Briefly, we chose to use Laplacian Eigenmaps (LEM) for non-linear dimensionality reduction due to its capability to effectively examine the local geometry of high-dimensional data^58–60^. This provides a finer-grain approximation, which is particularly beneficial when studying dynamics. Indeed, in our previous work we showed that this approach is extremely useful for studying insular cortex dynamics^57^.

We first estimated the intrinsic dimensionality in our data and found 5.2±0.8 dimensions, similar to our previous work examining mice in similar behavioral tasks^57^ (**Extended Data Fig. 6a**). We used non-linear dimensionality reduction, followed by automated clustering, which allowed us to describe activity dynamics as transitions between different subspaces (clusters) of the activity manifold (**Fig. 4a**). We examined the InsCtx activity manifold dynamics before and during optogenetic inhibition (**Fig. 4b).** Qualitatively, the optogenetic period did not occupy a distinct subspace (or cluster) in the InsCtx activity manifold (quantified by the number of shared clusters before and during optogenetic inhibition; **Fig. 4c; Extended Data Fig. 6b**), nor did it affect similarity of cluster transition dynamics (**Fig. 4d**).

**Figure 4:**
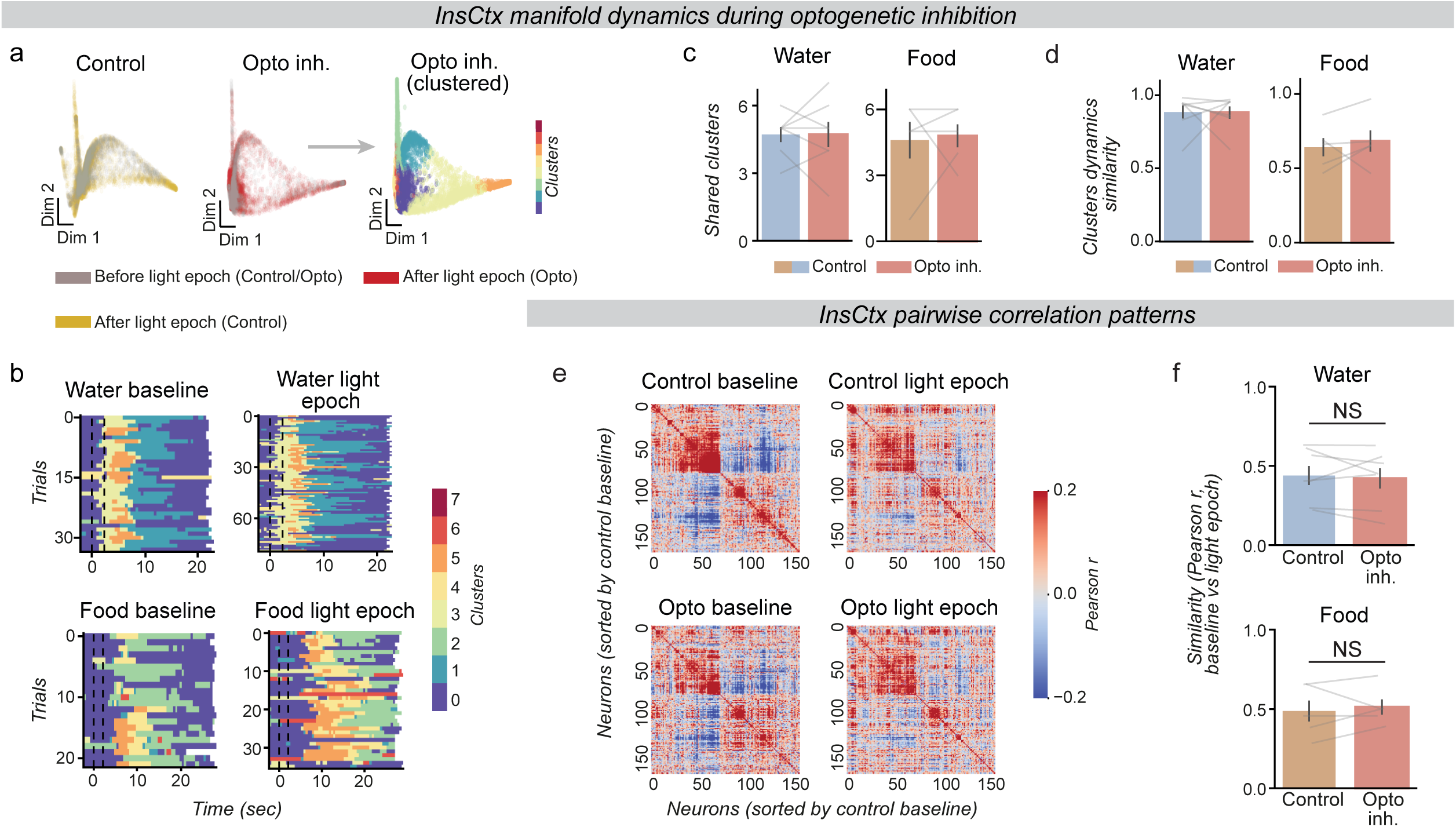
Inhibition of interoceptive thalamic inputs does not cause an abnormal InsCtx network state. a. Example of the lack of effect of optogenetic inhibition on the structure of the InsCtx activity manifold, shown via representative planes of dimensionality-reduced neuronal activity. Colors in the left and middle panels indicate different epochs (*gray*: baseline, *red*: optogenetic inhibition epoch, *yellow*: optogenetic inhibition epoch on the control day with no illumination). Note that due to the passage of time and reduced physiological need, the baseline and illumination epochs occupy partially overlapping yet distinct subspaces of the manifold, but do so similarly on both control (no light, *left*) and optogenetic inhibition days (*middle*). This suggests that optogenetic inhibition does not alter the InsCtx activity manifold. To quantify this similarity, the manifold was clustered into distinct activity clusters (*right*, same data as in the middle panel). b. Example from a single mouse showing manifold cluster sequence dynamics during water (*top*) and food (*bottom*) trials in the baseline versus optogenetic inhibition period. Dashed lines indicate cue period. Note similar dynamics before and during the optogenetic inhibition epochs, suggesting that it does not alter manifold cluster transition dynamics. c. Quantification of the effect of optogenetic inhibition on the activity manifold structure across mice. The number of shared clusters between the baseline and light epoch did not differ between control and optogenetic inhibition days for either water or food responses (water: n = 7 mice, 372±196, food: n = 5 mice, 325±131 neurons per mouse, for water and food, respectively; mean±SEM). d. Quantification of the effect of optogenetic inhibition on manifold cluster sequence dynamics, using the similarity of the distribution of trial clusters across epochs (see Methods). No significant difference in cluster dynamics similarity between control day 2 and the optogenetic inhibition day for both water and food trials (n = 7 mice, 372±196, food: 325±131 neurons per mouse, for water and food, respectively; mean±SEM). e. Example pairwise correlation matrices across all neurons in a water-restricted mouse, comparing baseline versus light-epoch periods on control versus optogenetic inhibition days. Note that neurons on all 4 days were sorted by the correlation structure during control day baseline (i.e., same sorting for all four matrices). *Top*: control day (baseline-left, light epoch-right). *Bottom*: optogenetic inhibition day (baseline-left, test/light epoch-right). Color scale: Pearson correlation coefficient (r). n = 165 neurons. f. Quantification of correlation-structure stability across control and optogenetic inhibition days, for water (*top*) and food (*bottom*) trials. For each mouse, the Pearson correlation coefficient was computed between the upper-triangle values of the baseline and test-period correlation matrices (as in ‘e’), yielding a single “stability” value per day, per mouse. Higher values indicates that the pairwise correlation structure across neurons is preserved between baseline and test periods. Water: n = 7, 372±196 neurons per mouse; mean±SEM. Food: n = 5,325±131 neurons per mouse; mean±SEM.

We further verified that these observations extend beyond the specific non-linear dimensionality reduction method (LEM). To do so, we performed the same clustering method but using a different non-linear dimensionality reduction (isomap^61^ instead of LEM), and observed similar results (**Extended Data Fig. 6c-f**). We also compared the pairwise correlation structure of neural activity^62^, further demonstrating that these conclusions do not rely on non-linear dimensionality reduction (**Fig. 4e-f**). Together, these different analyses suggest that InsCtx network activity remained in normal state during VPM/Lpc→InsCtx axon optogenetic inhibition.

Finally, we also assessed the effects of VPM/Lpc→InsCtx axon optogenetic inhibition on InsCtx activity using electrophysiological recordings of local field potentials (LFP) in anesthetized mice. Optogenetic inhibition caused a slow gradual increase in LFP power across a relatively broad band of frequencies (∼10-50 Hz) 5 minutes after illumination onset that was quite small yet statistically significant, suggesting a potential small general disinhibition (**Extended Data Fig. 6g-i**). This result is weaker yet somewhat consistent with previous work using pharmacological inhibition of VPMpc somata in rats^41^, with some differences potentially being due to the different inhibition method and/or species difference. Nevertheless, the converging analyses of our population calcium imaging data suggest that the small slow broad increase in LFP power we observed did not cause activity patterns to deviate from the normal repertoire off the activity manifold.

## Distinct representations of food and water in InsCtx

Our results regarding the effects of satiety and of VPML/pc→InsCtx inhibition raise a key question: what does InsCtx consummatory activity represent? Many previous studies have interpreted InsCtx activity during consumption as reflecting immediate oral sensory properties such as taste or texture^36,63–65^, while others have interpreted it as encoding affect, valence, or reward value^12,17,66,67^. Additionally, theoretical models of interoception have suggested a more integrative or predictive role for InsCtx responses^3,68^. Thus, to determine whether InsCtx consummatory responses reflect valence, value, sensory properties, and/or predictive computations, we performed a series of additional experiments.

We first tested whether InsCtx responses differ between water and food consumption, and whether they are need-specific. We imaged the same neurons across 4 different consecutive days, either food restricted or water restricted and with either access to food or water or both (**Fig. 5a**). Most neurons exhibited distinct responses to food and water, regardless of the other available reward, either when comparing them within the same day or across days (**Fig. 5b-c; Extended Data Fig. 7a**). Accordingly, population response vectors to the same reward (food or water) were highly similar across days within the same physiological need. In contrast, population vector similarity between food and water rewards was substantially lower, even when comparing them such that each is in its relevant physiological need (**Fig. 5d, Extended Data Fig. 7b-c**). Previous work has investigated the potential topographical organization of sensory information in InsCtx, with some studies observing clear organization and others that did not^69–73^. We thus examined the spatial organization of food and water responses and found no difference (**Extended Data Fig. 7d-f**). We note that because these experiments were done with a fixed order of first water restriction and then food restriction, the low similarity between food and water could potentially be influenced by this fixed order. To address this, we compared the food responses across day 1 (thirst) and day 3 (hunger). We found they were quite similar, as similar as responses on day 3 (hunger) and day 4 (hunger) (**Extended Data Fig. 7g**). This suggests that the distinct responses to food and water are not likely to result from the fixed order.

**Figure 5:**
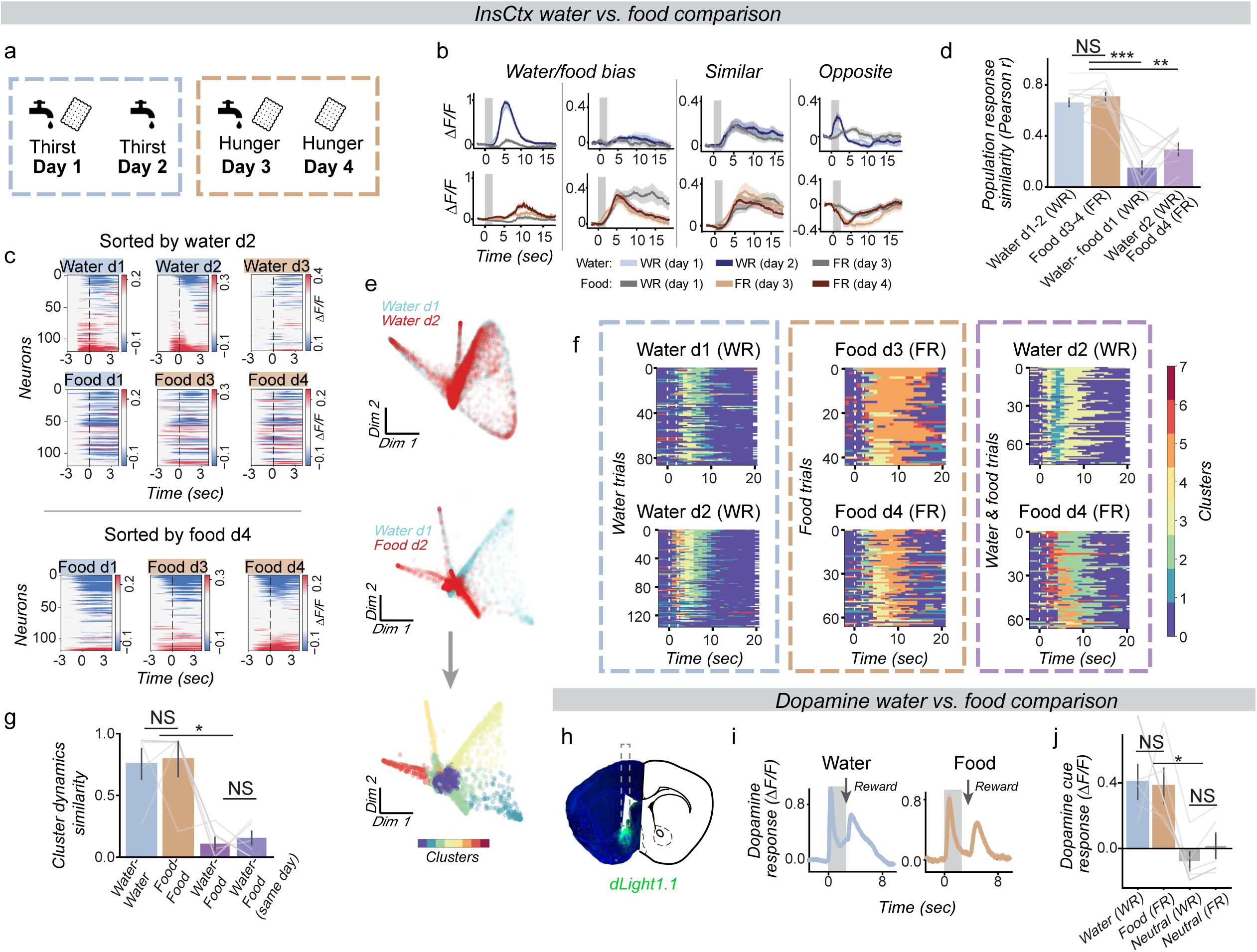
InsCtx food/water responses are distinct and do not reflect pure value. a. Experimental timeline: 4 days under different physiological need states and available rewards. Note that on different days mice were in different physiological need states (dashed rectangles) and had different available rewards - food and/or water - biscuit and faucet icons). b. Example single neuron responses with different response-selectivity to food/water across the different days of the experiment during different physiological needs. Gray rectangles: cue duration. Food restriction: FR, water restriction: WR. Values are mean ± SEM. c. Neuronal responses from an example mouse to food and water rewards across different days of the experiment during different physiological needs and available rewards. Same neurons were sorted by responses to water rewards on day 2 (*Top*, water and food) or to food rewards on day 4 (*bottom*, only food). Dashed line: consumption onset. n = 120 neurons. d. Population response similarity during water and food consumption. Note significantly higher similarity within same-reward consumption (water–water, food–food) compared to the other reward’s consumption (water– food). Same reward vs. water-food d1: ***p < 0.001, Same reward vs. water-food d2-d4 **p=0.001, water-food d1 vs. water-food d2-d4 p=0.093. n = 10 mice, 349±175 neurons per mouse, mean±SEM. e. Example representative dimensionality-reduced neuronal activity over two days (i.e., activity manifold). Each point represents the population activity pattern within a 1-sec time bin, colors indicate different days. *Top*: neuronal activity manifold during the same physiological need and rewards - water rewards during water restriction. *Middle*: neuronal activity manifold during different physiological needs - day 1: water rewards during water restriction (blue), Day 2: food rewards during food restriction (red). Note the low overlap in structure between the two days. *Bottom*: Same manifold as in the middle panel, shown with clustering into distinct activity clusters. f. Example manifold cluster dynamics in a single mouse across pairs of days. *Left*: two days of water restriction with water rewards. *Middle*: two days of food restriction with food rewards. *Right*: day 1 - water rewards during water restriction, day 2 - food rewards during food restriction. Note that for each comparison, cluster indices (colors) are randomly assigned during the clustering procedure. Dashed lines: cue. g. Quantification of manifold cluster dynamics response similarity: distribution similarity of trials-clusters across two days (see Methods). Manifold cluster dynamics similarity during the consumption of a single reward in the same physiological need was significantly higher than the comparison between two different needs and rewards. (water-water vs. water-food: *p = 0.01, food-food vs. water-food: *p=0.03, water-water vs. within day water-food: *p=0.03, food-food vs. within day water-food: *p=0.019), n = 5 mice, 332±134 neurons per mouse, mean±SEM. h. Example of a coronal slice showing dopamine sensor (dLight1.1) expression in the nucleus accumbens (green) and optic fiber track (dashed rectangle). i. Example trial-averaged fiber photometry recordings of dopamine dynamics in the nucleus accumbens in response to food/water cues during food/water restriction (respectively). Arrow indicates timing of reward delivery. j. Nucleus accumbens dopamine responses during food/water cue. Note no significant difference between responses to the two cues, suggesting similar expected values for water and food during the relevant physiological need in our behavioral task. Food/water cue responses were significantly different from neutral cue responses. Water cue: *p=0.041, food cue: **p=0.041, n = 5 mice.

To further examine population activity patterns within individual trials, we used our previously described approach for characterizing the InsCtx population activity manifold^57^, described above (**Fig. 5e**). This approach revealed highly consistent population activity patterns for food or water rewards separately within day and across days (**Fig. 5e top; Fig. 5f left and middle**). The *dynamics* of activity within and across trials were highly stereotyped between days for the same reward. In contrast, activity during food and water reward trials occupied distinct subspaces within the activity manifold and exhibited distinct cluster dynamics across days (**Fig. 5e, middle; Fig. 5f right**) or even within the same day (**Extended Data Fig. 7h**). We quantitatively validated these effects using the similarity in cluster distributions (**Fig. 5g**), and further confirmed them by quantifying cluster transition probabilities (**Extended Data Fig. 7i-j**). Thus, InsCtx population activity patterns during food and water consumption are distinct also at the level of individual trials.

The distinct responses between food and water suggest they do not predominantly represent pure valence. However, they could be explained by different “reward size” or “value” of food and water rewards in our task^74–76^ (e.g., one pellet would have the value of 2 water drops). To test this option, we recorded nucleus accumbens (NAc) dopamine signals during task performance using fiber photometry of a genetically-encoded dopamine sensor (diLight1.1; **Fig. 5h, Extended Data Fig. 8a**). Although NAc dopamine has multiple roles that are still debated^77–79^, NAc dopamine responses to reward-predicting cues are commonly accepted in simple behavioral tasks such as ours to reflect their expected reward value. Indeed, we observed robust NAc dopamine cue responses during both food cues and water cues (**Fig. 5i**), but not during the neutral cue, which induced a small decrease in the dopamine signal. The magnitudes of food cue and water cue dopamine responses were similar during their relevant need (**Fig. 5j, Extended Data Fig. 8b**), suggesting that the predicted reward value of food and water is roughly similar when examined in the relevant need. Thus, the different responses in InsCtx in our behavioral task likely do not reflect pure value. Importantly, we verified in the same mice that our recorded NAc dopamine signals were similar to canonical reward prediction error signals by observing negative responses to omission of water/food reward and enhanced responses to surprise water/food reward (**Extended Data Fig. 8c**).

## InsCtx food and water responses predominantly reflect need-specific prediction error signals when expectations are violated

We next tested whether the different InsCtx response to food and water could reflect oral-sensory properties and/or predictive responses. We designed another set of experiments in which mice received within the same day expected food/water rewards and omissions (10% omissions), as well as surprise delivery of food/water rewards after the neutral cue (10% of neutral cues; **Extended Data Fig. 8d**). Comparison of these 3 conditions allowed us to dissociate neuronal activity related to: [1] *consummatory actions* like licking or chewing (i.e., similar responses in all 3 conditions), [2] *consumption* of food/water (i.e., similar responses to expected reward and surprise reward, but different from reward omission, likely reflecting taste-related signals), [3] *predictions* (i.e., similar responses to expected reward and omission, but different from surprise reward), and [4] *prediction errors* (i.e., different between expected reward and omission or surprise reward, see further parcellation of prediction error types below; **Fig. 6a,g**).

**Figure 6:**
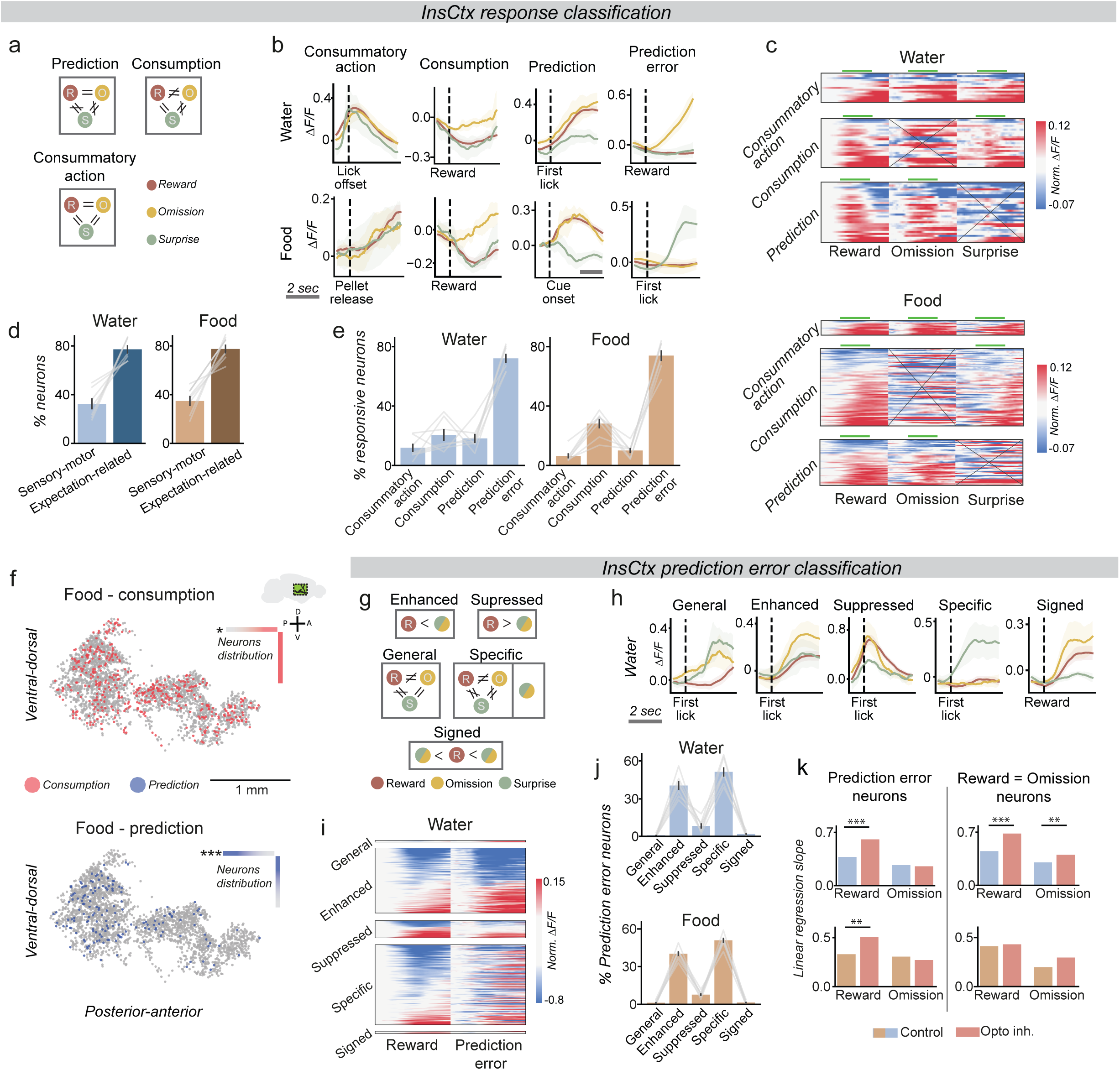
InsCtx food/water responses reflect prediction errors when expectations are violated. a. Classification of neurons to different response types. Neurons were categorized based on response shape and magnitude similarity of their responses to reward delivery, reward omission, and surprise reward trials. b. Examples of different classified response types to water (*top*) or food (*bottom*). Dashed lines indicate response alignment: reward delivery, cue offset, and lick onset/offset. Further details on the various prediction error types can be found in panel ‘h’. Note different time scale bar for the food-“prediction” example neuron, also corresponding to 2 sec. c. Example heatmap of single-mouse responses. Responses of individual neurons from one mouse to reward, omission, and surprise trials, organized by classification group. *Top*: classification of water responses, *bottom*: classification of food responses. Green lines indicate responses that resemble those of another trial type, while X marks responses that are distinct from all other types. d. Summary of response classification. Responses were first broadly grouped into two main categories: ‘sensory-motor’ (‘consummatory action’ and ‘consumption’) and ‘expectation-related’ (‘prediction’ and ‘prediction error’). The majority of responses were classified as expectation-related. n = 7 mice, 269±117, and 233±75 neurons per mouse, for water and food, respectively; mean±SEM. e. Summary of response classification. Percent of neurons exhibiting each response type (of all responsive neurons). n = 7, 269±117 and 233±75 neurons per mouse, for water and food, respectively; mean±SEM. f. Example of the topography of different response types. Locations relative to Bregma of all imaged neurons (dots) across mice, colored by response type. Colored gradients illustrate the spatial bias of each response type: uniform color indicates uniform distribution, while color gradients indicate areas of higher density. Asterisks indicate significant topographical gradient. *Top*: Food-consumption responses (anterior-posterior: *p=0.01, Dorsal-ventral: p=0.8, 432 ‘consumption’ neurons (red) out of total of 2553 neurons (gray)). *Bottom:* Food ‘prediction’ responses (anterior-posterior: ***p<0.001, Dorsal-ventral: p=0.3, 164 ‘predictions’ neurons (blue) out of total of 2553 neurons (gray)). n = 7 mice. See **Ext. Data Fig. 8h** for full data. g. Classification of neurons to different prediction-error response type. Neurons were categorized based on response shape similarity and magnitude across reward delivery, reward omission, and surprise reward trials. Note that the “Specific” category includes two response patterns: (1) neurons that respond exclusively to omission or surprise, and (2) neurons that respond to multiple outcomes, but with distinct response profiles (responses differ in both magnitude and similarity across conditions). h. Examples of classified prediction error response types. Neurons were categorized as “General” (similar omission and surprise responses), “Enhanced” (similar to reward but stronger response), “Suppressed” (similar to reward but weaker response), “Specific” (distinct from reward), or “Signed” (omission and surprise both resemble the reward response, with one stronger and one weaker). Examples are shown for water. Dashed lines indicate response alignment: reward delivery, cue offset, and lick onset/offset. i. Example of single-mouse prediction error responses. Heatmap showing responses of individual neurons from one mouse to reward delivery, reward omission, and surprise reward trials, sorted by prediction-error response classification. Note that ‘general’ and ‘signed’ responses were rare (see also ‘j’ below). j. Prediction error response classification summary. Percent of neurons exhibiting each response type of all prediction error responses. n = 7 mice, 197±96, food: 173±67 neurons per mouse, for water and food, respectively; mean±SEM. k. Quantification of the longer-term effects of optogenetic inhibition of VPM/Lpc→InsCtx axons on different response types. Correlation slope values of population activity before vs. after optogenetic inhibition for delivered and omitted reward responses (neurons pooled from all mice, control values are the mean of two control days for visualization purposes only). The inhibition caused an increase in slope for all groups except omission responses in the prediction error group. *Left-Prediction errors*: water-reward: day1 and day2 vs. opto, both ***p<0.001, water-omission: day1 vs. opto p=0.46, day2 vs. opto p=0.38. Food-reward: day1 vs. opto ***p<0.001, day2 vs. opto **p=0.009, food-omission: day1 and day2 vs. opto, both p=1.0. *Right-Reward=Omission*: water-reward: day1 and day2 vs. opto, both ***p<0.001, water-omission: day1 vs. opto p=0.074, day2 vs. opto **p=0.007, food-reward: day1 and day2 vs. opto, both p=1, food-omission: day1 vs. opto p=0.32, day2 vs. opto p=0.27. n = 5 mice. Water: *reward = omission* 710±6, *prediction error*: 530±104 neurons pooled across 5 mice (mean±SEM across the 3 days of the experiment). Food: *reward = omission* 314±28, *prediction error*: 397±26 neurons pooled across 5 mice (mean±SEM across the 3 days of the experiment).

We found that ∼91% of neurons responded to one or more of these experimental conditions (75.8±10% and 68.9±12% for food and water, respectively) and classified neurons into the 4 groups above (**Fig. 6b-c; Extended Data Fig. 8e**). Based on these 4 groups, we first broadly considered responses as either “sensory-motor” (“consummatory action” and “consumption”) or “expectation-related” (“predictions”, “prediction errors”). We note that each neuron could exhibit more than one type of response (e.g., a “consumption” response to expected reward and surprise reward, but a “prediction error” response to the reward omission, see **Fig. 6b,h**). Further analysis of the 4 groups revealed that 34% of InsCtx neurons had “sensory-motor” responses, whereas 77% of neurons exhibited “expectation-related” responses (**Fig. 6d**). Although individual neurons could appear in both categories, the responses predominantly involved distinct neuronal populations with minimal overlap (**Extended Data Fig. 8f)**. Surprisingly, the most prevalent response type was “prediction errors”, with ∼73% of all neurons exhibiting this response type, followed by “consumption” (∼24%; **Fig. 6e**). We found very little overlap between food and water (**Extended Data Fig. 8g**). Interestingly, we observed a spatial organization of responses, such that “consummatory action” and “prediction” responses were more prevalent posteriorly for both food and water responses. In contrast, “consumption” responses displayed different spatial distributions between water and food conditions (**Fig. 6f, Extended Data Fig. 8h-i**).

We further focused on “prediction error” responses, as they comprised a substantial fraction of ‘expectation-related’ responses. We observed responses akin to well-known types of prediction error responses (**Fig. 6g-h; Extended Data Fig. 9a**). Specifically, “signed prediction errors” (opposite signs to omission and surprise, e.g., “reward prediction errors”^80,81^), “general prediction errors” (also referred to as “unsigned prediction errors”^82^; similar responses to omission and surprise),”suppressed prediction errors” (appear as suppression of the reward responses) and also “specific prediction errors” (different response to omission or to surprise; **Fig. 6i; Extended Data Fig. 9b-c**). Interestingly, consistent with recent work in the visual system^83^, many prediction error signals appeared as an enhancement of the reward response (∼40%; **Fig. 6j**). We observed very little overlap between food and water prediction error signals (**Extended Data Fig. 9d**). Additionally, we did not observe a clear spatial organization of different types of prediction error signals (**Extended Data Fig. 9e**).

Importantly, we took great care to align responses between expected, omitted, and surprise rewards (see e.g., **Ext. Data Fig. 9f-g,** see Methods). We emphasize that our alignment procedure was conservative in that it was biased to find the maximal similarity between all three conditions, i.e., it is biased to identify responses are “consummatory action” (licking/chewing) or as “consummption” (e.g., taste). Therefore, any responses that were still identified as “prediction error” have passed this stringent criterion. Nevertheless, we further verified quantitatively that differential responses across conditions cannot be explained by lick latency, lick bout structure, lick rate, anticipatory licking rate, chewing-related orofacial movements, pellet reaching/grasping/handling, as well as by pupil size as a proxy for arousal levels (**Extended Data Fig. 9h-m**). Thus, InsCtx responses to food/water rewards reflect sensory-motor properties and predictions when expectations are met, but when expectations are violated most responses reflect need-specific prediction error signals.

Finally, we examined how VPM/Lpc→InsCtx inputs contribute to different response types. Because surprise reward trials were not included in these earlier experiments, we focused on comparing responses during delivered (90%) and omitted (10%) reward trials. Specifically, we identified “prediction error” responses as the responses during reward omission trials of neurons with significantly different response between delivered and omitted rewards (**Extended Data Fig. 10a-b**). Similar to the entire neuronal population (**Fig. 2**), optogenetic inhibition stabilized the responses of “prediction error” neurons to (expected) *delivered rewards* (**Fig. 6k**, *left*). In contrast, their (unexpected) *reward omission* responses (i.e., prediction errors) were not affected by optogenetic inhibition (**Fig. 6k**, *left*). In other neurons, in which delivered and omitted rewards evoked similar responses (i.e., “prediction” and “consummatory actions”), optogenetic inhibition stabilized both delivered and omitted reward responses, similarly to the entire population (**Fig. 6k**, *right*). In summary, the only population that was not affected by optogenetic inhibition was prediction error responses. Importantly, under normal conditions, prediction error responses were modulated by decreased physiological need (**Extended Data Fig. 10c**). Interestingly, “consummatory action” responses were consistently less influenced by decreased physiological need compared to both “consumption” and “prediction error” responses, in agreement with their more sensory-motor characteristics (**Extended Data Fig. 10c**). These results suggest differential thalamic contributions to different InsCtx response types (see more in **Discussion**).

In summary, our results suggest that when expectations are met, InsCtx responses reflect sensory-motor and prediction response. However, when expectations are violated, InsCtx responses are dominated by need-specific prediction error signals.

## Discussion

We combined a novel behavioral apparatus, longitudinal microprism-based two-photon imaging of InsCtx activity, and pathway-specific manipulations. We found that inhibition of interoceptive thalamic inputs to InsCtx kept InsCtx representation of physiological needs and related representations of food and water in their initial state, preventing their updating by the current need levels. Consequently, cue-driven feeding and drinking remained closer to the initial thirsty/hungry states, thereby preventing behavioral hallmarks of satiety. When expectations were violated, InsCtx food and water responses appeared to predominantly reflect prediction error signals, which are likely important for updating its predictive representations.

### Updating physiological needs in InsCtx

Previous human neuroimaging studies have suggested that InsCtx ongoing activity reflects hunger and thirst^28–30^. Interestingly, InsCtx appeared to be the only cortical site whose activity tracks bodily physiology changes related to hydration, and not “subjective water satiety”^29^. Indeed, we previously used two-photon population imaging of InsCtx to show that slow changes in ongoing InsCtx activity patterns reflect the current physiological state during water or food restriction^13^. However, such correlational studies could not test how physiological need representations were created in the InsCtx. We now provide experimental evidence for the updating of physiological needs by using precise optogenetic inhibition of thalamic interoceptive inputs to InsCtx. If thalamic interoceptive inputs to InsCtx provide it with continuous sensory input to form a sensory representation, then their inhibition should have deteriorated or eliminated this representation. In contrast, if thalamic interoceptive inputs to InsCtx serve to update an estimation, then their inhibition should have prevented updating and maintained the representation in the previous state regardless of the true current physiological need state.

In agreement with updating, we found that precise optogenetic inhibition of thalamic interoceptive inputs to InsCtx prevented the gradual transition of ongoing activity from activity patterns associated with hunger/thirst to those associated with sated/quenched states. Moreover, performing these experiments unilaterally allowed us to separate direct effects on InsCtx vs. indirect behavioral effect. Using bilateral optogenetic inhibition of thalamic interoceptive inputs to InsCtx, we showed that this information is also used to update behavior, mirroring the effects on InsCtx activity. However, it is important to note that while these results suggest that thalamic input is involved in updating, they do not exclude the possibility of a continuous cortical representation. That is, our results suggest that cortical representations can be maintained in the absence of visceral sensory thalamic inputs, but that does not necessarily mean that they do not naturally continuously change with incoming sensory inputs. We note that jamming of InsCtx activity in a hungry/thirsty state during unilateral inhibition, rather than a deterioration of representations, is unlikely to be explained by inputs from the contralateral (unperturbed) hemisphere as this should have then updated the activity to a sated state.

Our previous work using pharmacological inhibition of InsCtx found no effect on home-cage feeding^14^. The main difference between these two manipulations is whether or not InsCtx activity is preserved during the manipulation. In the case of local pharmacological inhibition using agonists of GABA-A and GABA-B receptors, InsCtx activity is quite inhibited. In contrast, the optogenetic inhibition of thalamic inputs preserved natural InsCtx activity patterns (Fig. 4) but prevented their gradual change to a sated/quenched state. We speculate that InsCtx provides another layer of regulation of feeding and drinking on top of hypothalamic and brainstem circuits. Therefore, during pharmacological inhibition of InsCtx only hypothalamic and brainstem circuits control feeding and drinking without InsCtx regulation. In contrast, during optogenetic inhibition of thalamic inputs, InsCtx activity, and therefore regulation of feeding and drinking, is maintained but is not updated.

Our previous work showed that while hypothalamic “hunger neurons” or “thirst neurons”^84–87^ powerfully regulate InsCtx cue response, they do not affect representations of physiological needs in InsCtx^13^. Together with our current results, this suggests that parallel hypothalamic and thalamic input pathways contribute to distinct aspects of InsCtx activity. Visceral sensory inputs are conveyed to InsCtx at least in part via the direct thalamic pathway from VPM/Lpc that we investigated here. In contrast the multi-synaptic pathway from hypothalamic “hunger neurons” and “thirst neurons” through the midline thalamic paraventricular thalamic nucleus (PVT) and basolateral amygdala appears to convey motivational state information that biases cue responses^13,14^. Importantly, there could be other parallel pathways conveying internal state information and/or visceral signals to InsCtx that have not yet been investigated. Future work could further examine how interoceptive hypothalamic and thalamic pathways interact across the brain to implement the hierarchical antagonistic interactions between different, often conflicting, physiological needs^88–92^.

### Prediction errors in InsCtx

InsCtx activity has been traditionally considered as primarily reflecting the sensorimotor aspects of consumption such as taste, chewing, licking, etc. ^36,63–65^. These oral representations have been shown to be modulated by associative learning^22,23,25,36,52,93–96^. More recent work has highlighted prediction signals in InsCtx, especially to need-fulfilling rewards^12–15,18,22–24^, as well as other valence related signals^12,17^. Interestingly, VPM/Lpc has also been shown to have expectation related signals^43^, which could potentially emerge via cortico-thalamic feedback from InsCtx.

Here, we used a classic comparison of expected, omitted, and surprise rewards, but doing so for two different need-fulfilling rewards under two different physiological needs. Using this straightforward approach, we found that InsCtx represents both sensory-motor (e.g., licking, chewing, taste), as well as prediction and prediction-error signals. Indeed, some of the differences between InsCtx food and water responses can be attributed to sensory-motor signals (i.e., for neurons we classified as “consummatory action” or “consumption”). In contrast, almost all the prediction and prediction-error signals we observed were need-specific and were not explained by arousal and sensory-motor parameters, suggesting that they do not represent general valence per se. These results are consistent with well-controlled human neuroimaging studies focusing on pain-related specific prediction and prediction errors in InsCtx^97,98^.

Similar to other sensory cortices, InsCtx has intermingled representations of sensory, motor, and more ‘cognitive’ signals^8,99–104^. Interestingly, consistent with recent studies of visual prediction errors^83^, many of the InsCtx prediction-error signals were enhancements of expected reward responses. This could explain why previous studies of InsCtx did not consistently observe them. These results suggest that enhancement of the representation of unexpected sensory stimuli could be a general feature of prediction error signals in sensory cortices.

Mid-posterior InsCtx has been suggested to compute prediction error signals by comparing prediction signals form anterior InsCtx with sensory information from interoceptive sensory inputs^3,8,20,21^, including the interoceptive thalamus in VPM/Lpc. While our current study identified prediction error signals in the InsCtx, it did not identify their precise source. Thus, these signals could be computed locally in the layer 2/3 neurons we imaged, or they could be computed in frontal brain regions such as medial prefrontal cortex, orbitofrontal cortex, or anterior cingulate cortex, and then relayed to InsCtx.

Physiological needs in InsCtx were represented both by ongoing activity levels, as well as by modulations of sensory-motor, prediction, and prediction error signals. Inhibition of thalamic inputs to InsCtx blocked the effects of satiety by stabilizing responses to delivered rewards (Fig. 2). As our thalamic axon optogenetic inhibition experiments included reward omission but not surprise rewards, we did not separately examine the role of thalamic inputs in each response type. Nevertheless, we could specifically isolate prediction error responses by focusing on neurons that respond differently to delivered and omitted rewards, and examining their reward omission responses, which therefore reflect prediction error signals. The reward omission responses of neurons with similar response to reward and reward omission (“*predictions*” and “*consummatory actions*”) were stabilized by thalamic inhibition (Fig. 6). In contrast, the prediction error responses of neurons that respond differently to delivered and omitted rewards were not affected by inhibition of thalamic inputs, as their omission responses kept gradually changing during the transition to satiety similar to control conditions. We propose two possible explanations for this observation: (1) The inhibition of thalamic inputs attenuated both prediction error neurons’ slow updating by physiological need state (i.e., their stabilized reward responses; Fig. 2), as well as the fast sensory input used to calculate the error signal (i.e., their unaffected omission responses; Fig. 6). This would result in similar changes to responses in both reward delivery and omission trials, which will falsely appear as no change from the control day. (2) Prediction error computation may rely on inputs from a brain region other than the VPM/Lpc, which is itself modulated by decreased physiological need, thereby leading to state-dependent changes in prediction error responses despite thalamic inhibition.

In summary, our results reveal an important role for interoceptive sensory information from thalamus to InsCtx in updating the representation of physiological needs and need-driven behavior. Aberrant updating of natural and pathological needs is likely an important aspect of different psychiatric conditions, including eating disorders and drug addiction^6,32,105^. Thus, our results further demonstrate how using animal models to complement studies in humans is important for developing a mechanistic understanding of brain dysfunction in such pathological conditions. Such insights include both a deeper conceptual understanding of how needs are formed, but also concrete specific pathways in the brain that underlie representations of natural and pathological needs.

## Acknowledgements

We thank Ofer Mazor, Hakan Kucukdereli, Osama Alturkistani, and Mark Andermann for assistance with early prototyping of the pellet device. We thank Lilia Gofer and Daniel Goldian for assistance with late prototyping and finalizing the pellet device. We thank Ofer Yizhar for advice with eOPN optogenetic protocols. We thank Sasha Devor, Yarden Cohen, and members of the Livneh lab for insightful discussions and comments on the manuscript. This work was supported by research grants from the European Research Council (ERC StG #101039145), Israel Science Foundation (ISF #860/21), the Peter and Patricia Gruber Awards, and the Center for New Scientists of the Weizmann Institute of Science.

## Author contributions

YP and YL designed the experiments and wrote the manuscript. YP performed all imaging, bilateral inhibition behavioral experiments, histology and data analyses. IP performed dopamine fiber photometry experiments, initial head-fixed bilateral inhibition experiments, and assisted with data analysis. YP, YL, and SS prepared prism mice. SS provided conceptual input. YL and KCKM performed electrophysiology experiments. IT developed the pipeline for the manifold analysis.

## Methods

### Experimental and subject details

All animal care and experimental procedures were approved by the Weizmann Institute of Science Institutional Animal Care and Use Committee. Animals were housed with standard mouse chow and water provided ad libitum, unless specified otherwise. We used both male and female mice and did not observe any differences between them.

Sample sizes were chosen to reliably measure experimental parameters, while remaining in compliance with ethical guidelines for minimizing animal use and keeping with standards in the relevant fields. Different animals in each experimental group served as replicates, and multiple repetitions of each cue also served as within-mouse replicates. Experiments did not involve experimenter-blinding, but randomization was used with respect to trial order and data collection. Animal subjects were not randomly allocated to experimental groups as all comparisons were performed within subject.

### Surgical procedures

C57BL6 mice (8-12 weeks old) were anesthetized using isoflurane in air (induction, 3%; maintenance, 1%-2%), placed into a stereotaxic apparatus (RWD) on a heating pad, and ophthalmic ointment was applied to the eyes. For postoperative care, mice were injected intraperitoneally with Carprofen (5 mg/kg), Buprenorphine (2 mg/kg) and saline (1 ml).

#### Stereotaxic injections

After exposing the skull via a small incision, a small hole was drilled for injection. A pulled-glass pipette with 20-40 µm tip diameter was inserted into the brain, virus was injected using an air pressure system (PV850, WPI), and the pipette was withdrawn 5 min after injection. We used the following volumes of virus and injection coordinates: **VPM/Lpc** (AAV5-mCaMKIIa(short)-eOPN3-mScarlet : 300 nL, Bregma: AP: -2.3 mm, DV: 3.7 mm, ML: 0.75 mm), **NAc** (AAV1-CAG-dLight1.1: 300 nL, Bregma: AP: +1.4 mm, DV: 4.6 mm, ML: 0.75 mm). **InsCtx** (AAV8-hSyn-GCaMPf7 diluted 1:3, 300 nL, Bregma: AP: 0 mm, DV: 3.8 mm, ML: edge of the brain).

#### Optic fiber implantation for optogenetic inhibition and fiber photometry

First, mice were stereotaxically injected with AAVs, as described above. An optic fiber (400 µm diameter core NA 0.5; ThorLabs) was then implanted (NAc: AP: +1.4 mm, DV: 4.5 mm, ML: 0.75 mm from Bregma, InCtx: AP: 0 mm, DV: 3.7 mm, ML: edge of the brain, mm from Bregma). The fiber was fixed to the skull using CCB Metabond (Parkell). Mice were allowed at least 4 weeks for recovery before starting behavioral training. Using aseptic technique, a custom-made headpost was secured using cyanoacrylate glue, dental acrylic and CCB Metabond (Parkell).

#### Microprism assembly and surgery

Glass microprism assemblies were fabricated using 2 mm microprisms (#MCPH-2.0; Tower Optical) coated with aluminum along their hypotenuse. Microprisms were attached to a cover glass (#1 thickness), both along the hypotenuse (to prevent scratching of the reflective surface) and at the side of the prism that faces InsCtx, using Norland Optical Adhesive 81 cured using ultraviolet light. The glued prism assembly was then secured to a cannula using Kwik-Cast (WPI), allowing controlled insertion of the prism into the brain.

Mice were anesthetized and prepared to surgery as described above. Using aseptic technique, a custom-made headpost was secured using CCB Metabond (Parkell). The mouse’s head was then rotated ∼70°. A ∼2.2×2.2 mm^2^ craniotomy was made over the left InsCtx, with the lower edge just above the squamosal plate. AAV8-hSyn-GCaMPf7 was diluted in PBS 1:3, mixed with FastGreen for visual confirmation, and injected directly into the exposed brain tissue at a depth of 0.2 mm, (250 nL × 4; PV850, WPI). The brain surface was kept hydrated throughout the procedure using saline-soaked gelfoam. The mouse’s head was then rotated back and the 2×2 mm^2^ microprism attached to the cannula was slowly lowered into the craniotomy until it made contact with the InsCtx. It was then advanced further while being slightly shifted medially (100-200 µm), ensuring that the lower edge of the microprism extended below the squamosal plate and contacted the medial wall of the craniotomy. Once properly positioned, the edges of the window were sealed to the skull with Vetbond (3M), followed by reinforcement with CCB Metabond (Parkell). The cannula was gently removed, and additional Metabond was applied to form a permanent seal. Mice were allowed to recover for at least six weeks prior to the start of behavioral training.

### Bilateral optogenetic inhibition illumination

For all bilateral optogenetic inhibition experiments, including the head-fixed cued Pavlovian task, home-cage free consumption, and fiber photometry recordings in anesthetized mice, each animal underwent two sessions: a control session with no light stimulation and an inhibition session with light delivery. Optogenetic inhibition consisted of alternating 30-second on/off epochs (625 ± 7 nm LED from Prizmatix, 15 Hz, 30% duty cycle, ∼9 mW at the fiber tip).

### Bilateral optogenetic inhibition and GCaMP fiber photometry recordings during manipulation of visceral inputs

To test whether optogenetic inhibition blocks visceral signals from reaching the InsCtx, we performed fiber photometry recordings (see further details in the *Fiber Photometry* section below) during intraperitoneal injection of interoceptive stimuli, on control and bilateral optogenetic inhibition days. Mice were anesthetized during recordings (∼1.5% isoflurane) to isolate responses to the injected substances from associated expectation or stress of the injection itself. We examined four stimuli related to hunger and thirst: in overnight food-restricted mice, Exendin-4 (240 mg/kg) or CCK-8 (5 μg/kg). In water-restricted mice, saline injection (1 mL), and in non-restricted mice, NaCl injection (2M, 200 μL). This yielded two measures, each for hunger and thirst-related signaling. On control days, each session consisted of a 5-minute baseline period, a first (sham) injection, a further 5-minute interval, then a second injection of substance, followed by a final 40-minute recording period. On inhibition days, the baseline period was 3 minutes, after which light delivery began immediately, with the first (sham) injection following 2 minutes later. As an additional control, food-restricted mice also underwent a control session, in which both injections were sham.

### Unilateral optogenetic inhibition during electrophysiological LFP recordings

Four to six weeks post-injection, we anesthetized mice with urethane (800 mg/kg) and chlorprothixene (10 mg/kg). We then exposed the insula and recorded LFPs using borosilicate micropipettes with a resistance of ∼7 MΩ, at a depth of ∼260 µm to target layer 2/3. Signals were acquired using an Axoclamp-700B amplifier (Molecular Devices), band-pass filtered at 0.1-300 Hz, and digitized at 20 kHz. We used a 637 nm LED at a power of 9 mW, pulsed at 15 Hz with a 30% duty cycle, in 2-minute on/6-minute off epochs, similar to our two-photon imaging illumination protocol. Each recording began with an 8-minute baseline period prior to illumination onset.

### Two-photon imaging

Two-photon imaging was performed using a resonant-scanning two-photon microscope (Thorlabs Bergamo) at 30 frames/second, via a 10×0.5 NA air objective (TL10X-2P, Thorlabs) at 1.5 zoom, yielding a 906×906 µm field of view (512×512 pixels), using a 920 nm fixed-wavelength laser for GCaMP imaging (Toptica FemtoFiber ultra 920) and a 1050 nm fixed-wavelength laser for eOPN3-mScarlet structural imaging (Toptica FemtoFiber Uutra 1050). All imaged fields of view (FOV) were at a depth of 90-200 mm below the pial surface, with laser power at 920 nm of 15-80 mW at the front aperture of the objective (power at the sample was likely substantially less due to partial transmission via the microprism). Mice were imaged in consecutive 30-minute runs. Imaging depth was adjusted in between runs (every 30 min) to account for slow drift in the z plane (< 7 µm).

#### Two-photon imaging during optogenetic inhibition

Optogenetic inhibition was performed between imaging frames. Light was switched on and off only between trials, not during trials. We used a 620 nm Thorlabs LED with a power of 9 mW after the objective. To control light on/off timing, we used an Arduino Uno, which switched states in response to commands sent via the MonkeyLogic2 program. The imaging frame rate was reduced to 15 Hz to allow sufficient time for inhibition (33 ms imaging per frame, within a 66.6 ms cycle). The light was turned on 5 ms after frame acquisition, remained on for 20 ms, and then turned off, resulting in a 30% duty cycle. Light delivery followed a schedule of 2 minutes on and 6 minutes off.

### Fiber photometry

To allow for adequate viral expression, dopamine and GCaMP fiber photometry recordings were performed no less than four weeks after the surgical procedure. We used fluorescent miniscube with 405 nm LEDs to excite the isosbestic control channel and 465 nm LEDs to excite the activity-dependent GFP-related channel (dLight1.1, GCaMPf7; Doric Lenses). Recordings were conducted using pyPhotometry hardware and software^106^ at a sampling rate of 45 or 50 Hz, and the data were analyzed using custom Python code.

### Brain tissue preparation and immunohistochemistry

Fiber photometry and bilateral optogenetic inhibition mice were intraperitoneally injected with Pentobarbital (0.1 ml per 10gr body weight). Then, mice were perfused transcardially first with phosphate-buffered saline (PBS) to clear the vasculature, followed by 10% neutral buffered Formalin (Sigma Aldrich) for tissue fixation. The brains were carefully extracted. All microprism mice were intraperitoneally injected with Pentobarbital (0.1 ml per 10gr body weight), and their head were placed in 10% neutral buffered formalin. After several days the brains were carefully extracted and placed in formalin. Then, brains from all mice were cryoprotected in 30% sucrose solution prepared in PBS and then sectioned coronally at 50 μm using a freezing sliding microtome (Leica Biosystems). Sections were collected in three equal series. Before mounting, brain sections from dopamine fiber photometry mice were washed in PBS and incubated overnight at room temperature in blocking solution (3% fetal bovine serum and 0.25% Triton in PBS) containing primary antibody (1:1000, anti-GFP chicken. Invitrogen) on a shaker. The next morning, sections were washed 3 times in PBS with ten minutes on shaker between each wash and then incubated in Alexa fluorophore-conjugated secondary antibody (1:1000, anti-chicken 488 goat, Invitrogen) for 3h at room temperature. After that they were washed 3 times in PBS before the mounting. Sections were mounted onto gelatin-coated slides and fluorescent images were captured with an Olympus BX61VS slide scanner microscope at 10X. All antibodies used were previously verified.

### Postmortem identification of imaging field location

To localize the imaging site within the brain, we first acquired in vivo epifluorescence images of the cortical surface vasculature around the recording location using LED illumination. Post-mortem, prior to brain sectioning, we also captured whole-mount widefield fluorescence images of the exposed cortex. These images provided a view of the blood vessel patterns, GCaMP expression, and the prism indentation (approximately 2.1 mm²), enabling alignment between in vivo imaging and anatomical features. We then sliced the brains to determine the precise location of the implanted prism. By integrating this anatomical information with the in vivo field of view observed through two-photon imaging (approximately 0.9×0.9 mm), we reconstructed the exact imaging location within the prism.

### Behavior protocols for comparisons across need states, thalamic axon optogenetic inhibition, and response types

#### Pavlovian behavioral training

After at least 2 weeks of post-surgical recovery, animals (10-16 weeks old) were water restricted to ∼80-85% of their free-feeding body weight. Animals were head-fixed under the two-photon microscope for habituation prior to any behavioral training (10 min to 1 h). If the mice displayed any signs of stress, they were immediately removed and additional head-fixation sessions were added until there were no visible signs of stress. Once the mice were relaxed, they were given water manually via a syringe to acclimate them to licking. The syringe was then placed in the behavioral device (**Fig. 1a**), and the mice learned to lick water from the syringe positioned on their right. To train the animals to associate water delivery (5 μL) with a specific odor cue (Pavlovian conditioning), the water-predictive odor was presented, followed by unconditional water delivery, for 30 minutes. Training continued until consistent anticipatory licking was observed during the cue period. Next, we added neutral trials, predicted by a different odor, and trained the mice with both trial types. On day 1, inter-trial intervals (ITIs) were short (8 seconds), and on day 2, ITIs were extended to 25 seconds. Training lasted up to two hours per day, continuing daily sessions until mice stopped incorrectly licking during neutral trials. Following this phase, mice were food-restricted overnight and trained the next day to consume food pellets. Initially, food was delivered without any odor cue. The pellet-cup was positioned as close as possible to the mouth to help mice learn to obtain them without using their paws. After 1-2 hours (depending on individual progress), we introduced the neutral cue and a 3^rd^ odor as the food-predictive cue. On the second day of food training, water trials were reintroduced. After successful consumption of both water and food within the same session, mice were returned to water restriction. They continued with at least five additional days of water and food training under water-restricted conditions. All conditions were always presented in equal proportions. The final three training days occurred during two-photon imaging, allowing mice to acclimate to the imaging environment. Licking behavior for both water and food was tracked using a capacitance-sensing lickspout system: a blunt metal syringe needle was used for water, and a thin metal fiber wrapped around the pellet-cup was used for food. Both were connected to an MPR121 capacitance sensor. All behavioral training was controlled using MonkeyLogic2^107^ (Asaad and Eskandar, 2008; https://monkeylogic.nimh.nih.gov/). The behavioral task required water or food-restricted mice to discriminate between three different odors. Citral served as the water cue, Ethyl Butyrate as the food cue, and Isoamyl Acetate as the neutral cue. Odors were diluted to 10% (v/v) in mineral oil, and further air-diluted by to 1% using a custom built olfactometer. The olfactometer was designed such that it maintained a constant airflow of 1 L/min, with a clean air channel (10% of total flow) sporadically transiently replaced with odorized air for 2 sec. We defined mice as well-trained if they exhibited anticipatory licking to the correct spout (water and food) only in response to the appropriate predictive cue (**Fig. 1b-d**).

#### Dopamine recordings

These experiments were carried out over four consecutive days, each consisting of a two-hour session. On the first day, mice were water-restricted and received only water and neutral trials. On the second day, the same water-restricted mice received water, food, and neutral trials. On the third day, mice were food-restricted and again received water, food, and neutral trials. On the fourth day, food-restricted mice received only food and neutral trials. Across all sessions, 10% of the reward-predictive trials were randomly assigned as reward-omission trials, where the odor cue was presented without subsequent reward delivery. Additionally, on days one and four, 10% of the neutral trials were replaced with surprise reward trials, in which a reward was delivered without any preceding odor cue.

#### Bilateral optogenetic inhibition during head-fixed cued Pavlovian task

Sessions were conducted in the same mice given only water and neutral cues (1:1) when water-restricted, or receiving only food and neutral cues (1:1) when food-restricted. Each session continued until the mouse voluntarily terminated reward consumption for at least 15 minutes (total session duration: 90-180 minutes). On inhibition days, sessions began with a light-off baseline period (10 minutes for water, 20 minutes for food), with a baseline duration that was calibrated to precede satiation. Following the baseline period, light was delivered for the remainder of the session. Two mice did not voluntarily stop food consumption during thalamic axon optogenetic inhibition. In these cases, the session was terminated once consumption duration exceeded that of the corresponding control session by more than one hour.

#### Bilateral optogenetic inhibition during home-cage free consumption

We examined the effect of optogenetic inhibition during free consumption in the home cage. Each mouse underwent a control and an inhibition session, while connected to the patch cords in both sessions. On inhibition days, light delivery began immediately upon reward consumption onset and continued until the end of the session. For food consumption: mice were food-restricted overnight prior to control and inhibition sessions, then placed in their home cage with a single large food pellet (∼6 g) and free access to water. Pellet weight was measured every 30 minutes over a 3-hour session. Water consumption: Mice were water-restricted for 3 days prior to control and inhibition sessions, then placed in their home cage with access to a lick spout only (no food), which delivered 5 μl of water per lick, allowing simultaneous measurement of water intake and lick counts. Sessions lasted 25 minutes with typical total consumption of ∼0.8 mL.

#### Imaging across different physiological need states

To assess how changes in thirst and hunger states during water and food consumption affect neural activity, we imaged mice across 4 consecutive days. On the first day the mice were water restricted, and received water, food and neutral trials, for 2-3 hours. Mice did not show signs of satiation on this day as they did not voluntarily stop consuming food or water until the end of the recording. On the second day mice were water restricted and received only water and neutral trials, until they reached satiation (voluntary cessation of consumption). Mice typically consumed approximately 1 mL of water during this day. We continued the imaging for an additional 30 minutes after mice voluntarily stopped performing the task. After the end of this session, mice were switched to food restriction. On the third day, mice were food restricted, and received water, food, and neutral trials. On the fourth day, mice were food restricted and received food and neutral trials. We imaged behaving mice for 1-1.5 hrs, while they consumed ∼1.2 g of food pellets. 10% of all reward trials were “reward-omission”, when no reward was delivered. Since mice on the fourth day received only pellets with no access to water, their consumption behavior could in principle reflect increasing thirst rather than satiation to food alone. To dissociate these factors, we recorded an additional session on this day. After 1-1.5 hours of task performance under the two-photon microscope, mice were returned to their home cage with free access to food and water for approximately 4 hours, allowing them to reach satiety and full rehydration. They were then placed back under the two-photon microscope for an additional ∼1-hour session, receiving only food and neutral trials as before. Mice still consumed a small number of pellets during this session. This design allowed us to examine food-related satiety independent of thirst induced by pellet-only consumption.

#### Identifying prediction and prediction-error signals

This experiment lasted two days. On the first day the mice received water and neutral trials until reaching satiation, similarly to day 2 in the experiment described above. Mice were then switched to food restriction, and on the third day were given only food and neutral trials, similar to day 4. In these two days, 10% of the reward trials were “omission” trials in which the reward was not delivered after the cue. Additionally, 10% of the neutral trials were “surprise” trials, in which the reward was delivered after the neutral cue.

#### VPM/Lpc→InsCtx inhibition during quiet waking

Mice were imaged over two consecutive days without any behavioral task or food/water restriction. The first day served as a no-light control, during which the mice were imaged for 56 minutes at rest. On the second day, imaging began with a 4-minute baseline period, followed by 4 cycles of optogenetic inhibition (15 Hz, 30% duty cycle, 6 minutes off / 2 minutes on, totaling 32 minutes). After the illumination cycles ended, imaging continued for an additional 20 minutes.

#### Unilateral optogenetic inhibition via the microprism during two-photon imaging

We first recorded two control days during which food-restricted or water-restricted animals received food or water and neutral trials, and continued the task until they voluntarily stopped consumption (typically 2-3 hours). In food restriction days mice consumed ∼2.3 g of food pellets. As in the food-restriction day described above (“Imaging across different physiological states”), each food-restriction day was followed by a refeeding and rehydration break, followed by an additional short session to dissociate satiety from thirst (this additional session did not include optogenetic inhibition).

On the optogenetic inhibition day, the recording began with a 20-minute baseline period without optogenetic inhibition (60 trials). This was followed by 8 cycles of optogenetic inhibition, each consisting of 6 minutes off and 2 minutes on (15 Hz, 30% duty cycle, 4 cycles per run), over the course of one hour. After these cycles, the animals continued the task until they voluntarily stopped consumption. For the water-restriction condition, we recorded behavior over three consecutive days. In the food-restriction experiment, due to the refeeding period, we included a one-day break between recording sessions to ensure the mice remained sufficiently food restricted throughout the 5-day experiment. All days included 10% omission trials.

### Behavioral data analysis

#### Data collection - two-photon imaging

Behavioral data included five inputs: (1) odor cue onsets, (2) water spout licks, (3) food cup licks, (4) upper beam break, triggered when a food pellet was dispensed, and (5) lower beam break, triggered when uneaten pellets fell through a gate-controlled opening, 15 seconds after delivery. Lick events and beam breaks were recorded via sensors connected to the behavioral apparatus, while cue onsets were delivered and time-stamped through the MonkeyLogic2 system. All behavioral events were binary and synchronized using the ThorSync system, which acquired data at 6000 Hz and simultaneously received two-photon imaging frame triggers. Behavioral data were subsequently down sampled to match the frame rate of the neuronal recordings (30 or 15 Hz). For each two-photon frame, behavioral variables were summarized. In instances where temporal resolution higher than the frame rate was required (e.g., lick detection at 15 Hz or rare occurrences of consecutive pellet deliveries detected by the upper beam break), relevant behavioral events were registered at the original sampling rate before down-sampling. Additionally, pellet movement and consumption were recorded using a video camera, with each video frame triggered by the two-photon imaging frames. An infrared LED provided illumination for video capture.

#### Data collection - bilateral inhibition with head-fixed Pavlovian task

Behavioral data were collected as described above, using the same five inputs (odor cue onsets, water spout licks, food cup licks, upper beam break, and lower beam break). Signals were acquired and synchronized using a custom Arduino script at a sampling rate of 100 Hz, then down-sampled to 20 Hz. Pellet movement and consumption were recorded using a video camera at 20 Hz, aligned to every fifth acquisition sample. An infrared LED provided illumination for video capture.

#### Data collection - bilateral inhibition during free consumption

For food consumption sessions, food intake was measured by weighing the pellet at 30-minute intervals, as described above. For water consumption sessions, licks and reward deliveries were recorded using the MonkeyLogic2.

#### Consumption onset detection using DeepLabCut

A key behavioral event in our analysis was the onset of reward consumption, defined as the moment of water or food intake. Water consumption was identified as the first water spout lick following reward delivery. Detecting the precise timing of food pellet consumption required further consideration. Although the lower beam break sensor confirmed whether a pellet was taken, it did not provide temporal precision for the consumption event itself. Licking events were not always reliable indicators of pellet consumption, as mice occasionally licked after consumption, took pellets without triggering the lick detector, or dropped the pellet before consumption. To accurately identify actual pellet consumption, we used DeepLabCut^35^ to track pellet position in each video frame. The model was trained on datasets representing diverse experimental conditions, including variations in camera angle, lighting, multiple pellet deliveries, and instances where pellets fell out of the mouse’s mouth after partial chewing. Video brightness was adjusted in some cases to optimize detection accuracy. All videos were processed using the trained DeepLabCut model to extract precise pellet location coordinates, with the *y*-axis position serving as the primary marker. We integrated these positional data with other behavioral variables to classify trial outcomes (consumed, not consumed, or problematic trial) and to determine the exact consumption onset. Consumption onset was defined as the moment the pellet disappeared into the animal’s mouth. In typical trials, the sequence proceeded as follows: activation of the upper beam break, appearance of the pellet in the video with a corresponding decrease in *y*-position as the pellet approached the food cup, followed by potential licking events. If the pellet was consumed, its *y*-position increased (movement towards the mouth) before it disappeared from the frame. If the pellet was not consumed, its *y*-position continued to decrease after the fate gate opened, followed by activation of the lower beam break. This setup also allowed us to identify and exclude rare problematic trials, including instances where the pellet became stuck before reaching the food cup, failed to fall after the gate opened, or when fragments of pellet triggered the lower beam break despite successful consumption. All video processing and classification were performed using Google Colab. Final consumption classifications were manually verified to ensure accuracy.

#### Anticipatory licking

To assess the animals’ discrimination of olfactory cues, we quantified water spout or the pellet cup anticipatory licking during the different cues. These analyses were performed on both thirst-motivated and hunger-motivated days, during sessions where all three trial types were presented in equal proportions. For each animal, we analyzed the first 20 rewarded trials. For water-predictive trials, anticipatory licking was quantified as the lick rate during the 2-second olfactory cue presentation. In food-predictive trials, anticipatory licking was calculated as the lick rate during the cue presentation plus the additional delay period (∼1 second) preceding pellet delivery. To enable comparisons across animals, licking rates were normalized within subjects. Water anticipatory licking was normalized to the average licking rate on thirst-motivated days where only water-predictive and neutral cues were presented. Similarly, food anticipatory licking was normalized to the licking rate on hunger-motivated days with only food-predictive and neutral cues. For sessions acquired at 15 Hz, to ensure accurate lick quantification, individual lick events were extracted directly from the original ThorSync recordings acquired at 6000 Hz.

#### Orofacial movement quantification using Facemap

To quantify orofacial movement, we used Facemap^108^ to extract motion energy from an ROI covering the orofacial region in each session’s video. The ROI was manually defined around the orofacial region for each session. Motion singular value decomposition (motSVD) was applied to this ROI, and the mean absolute value of the first three motion components was used as a summary measure of orofacial movement for each frame. This measure was normalized per session before further analysis.

#### Pupil size

Pupil diameter was extracted from eye videography using a custom MATLAB pipeline. For each session, the eye region was manually cropped, masked to exclude non-pupil regions (e.g., IR reflections), and calibrated for rotation and aspect ratio. Pupil boundaries were then detected in each frame using circular Hough transform-based circle detection, with the best-fit circle selected based on similarity to the previous frame’s radius. Frames failing quality-control criteria (e.g., large frame-to-frame displacement or abrupt size change) were replaced with the last valid detection. Pupil diameter was defined as twice the fitted radius. Frames with insufficient signal quality were flagged as bad frames and excluded from further analysis. Pupil diameter was normalized to the maximum diameter observed within each session and segmented into trials for subsequent analysis.

#### Behavioral quantification-bilateral optogenetic inhibition

To compare behavior between control and inhibition sessions, we defined two equal time windows for each mouse: one from a baseline period, comprising trials preceding the light trigger onset, and one from a test period near the end of the session, whose length was scaled to the duration of the illumination period (i.e., either same duration as the baseline period, or 0.25 of the illumination period, the smaller of the two). Window boundaries were computed from the control session, using the last trial in which the mouse continued drinking, defined as at least one lick occurring more than 4 s into a water trial, and applied identically to the corresponding inhibition session. Within these windows, we computed several licking-related measures per trial, averaged across trials in each window: anticipatory licking (licks during the 2-second interval following cue onset, before reward availability), total licking in the 5 s after cue offset, all licks after cue offset, and latency to the first lick following reward availability. A latency measure was also computed for food consumption, based on pellet consumption onset (see above) relative to cue onset. For each measure, baseline-normalized values were compared between control and inhibition sessions.

#### Behavioral quantification-unilateral optogenetic inhibition during two-photon imaging

We tested trials during the optogenetic inhibition window on the inhibition day, and the corresponding matched time window on control days. For water trials, we quantified lick rate aligned to cue onset and to reward delivery, anticipatory licking (licks in the 0-2 s cue window), and licking bout duration (interval between the first and last lick following cue onset). We additionally quantified drinking bout structure: bout number and mean bout duration, with bouts defined as lick sequences exceeding both a minimum duration and a lick-rate threshold, the latter derived from each day’s own early trials. For food trials, we quantified latency to pellet consumption (from DeepLabCut, see above) and a chewing-related movement measure, using the motion SVD-derived signal described above, taken as the average signal between its rise and subsequent decline following reward consumption. Values from the second control day served as the within-mouse reference for normalization, with the inhibition and first control days each expressed as a ratio to it.

Pupil size was compared during two-photon recordings between the second control day and the inhibition day only from light-off periods. For each trial, we computed the change in pupil diameter from a pre-cue baseline (-2-0 s) to a post-cue response window (2-4 s), restricted to trials occurring within the optogenetic window and excluding trials adjacent to light-offset transitions, and compared mean per-mouse deltas between the two days. We required a minimum number of 10 trials, with 5 frames per trial, that passed pupil analysis quality controls.

#### Consumption quantification during two-photon imaging

For each day, we additionally quantified the percentage of cued trials in which the reward was consumed. Unlike the measures above, this was reported as raw values across all three days rather than normalized to the second control day.

#### Pellet grabbing

Pellet grabbing and hand movement appeared rarely and only in one mouse. Hand location was tracked using DeepLabCut as described above, and trials were classified as containing hand movement if the tracked hand position exceeded a fixed displacement threshold during the pellet delivery period. We compared this measure only between the second control day and the inhibition day. The proportion of trials with hand movement in this one mouse did not differ significantly between these two days (Fisher’s exact test, p = 0.21).

### Two-photon imaging data analysis

#### Image registration and time course extraction

Two-photon imaging data were first processed using Suite2p^109^ for image registration. Within each imaging session, all runs (2-8 runs per session) were aligned to the first run of the day. To improve signal quality, recordings were subsequently denoised using DeepInterpolation^110^. Cellular calcium activity traces and neuropil signals were then extracted using Suite2p. Cell masks were manually inspected to ensure typical soma morphology, appropriate size, and normal activity patterns. To track individual neurons across days, we employed CellReg^111^. Only cells exhibiting clear, consistent signals across days were included in the further analyses. Neuropil-corrected fluorescence time courses were calculated as:

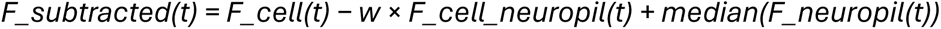

where F_cell(t) is the raw fluorescence signal from the region of interest, F_cell_neuropil(t) is the surrounding neuropil signal, and w is the neuropil contamination coefficient estimated by Suite2p. After motion correction (see below, F_corrected), changes in fluorescence were calculated as:

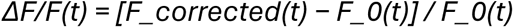

where F_0(t) represents a dynamic baseline fluorescence estimate, defined as the 10th percentile of F_corrected(t) within a 4.5-minute sliding window. For visualization of example single-neuron traces, fluorescence time courses were re-zeroed using the 2-second period preceding odor cue onset.

#### Motion correction

Lateral motion along the x- and y-axes was extracted from Suite2p. Both x and y displacement traces were z-scored. Additionally, we computed a combined movement index as 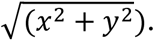 Neuropil-subtracted calcium traces were first high-pass filtered to remove slow fluctuations (real responses), using a cutoff frequency optimized for each animal (typically between 0.1-0.2 Hz). The filtered traces were z-scored and segmented into trials. Trials shorter than a threshold, defined as the number of recorded cells plus 300 frames, were concatenated. Subsequent processing was performed jointly across all cells within each trial. First, principal component analysis (PCA) was applied to the trial data. The minimum number of principal components required to explain at least 50% of the variance was retained, with a minimum of two components. The selected principal components were then subjected to independent component analysis (ICA). Each resulting independent component underwent two additional processing steps: a low-pass filter at 0.4 Hz to isolate slow fluctuations, and separate smoothing using a 0.33-second moving window. We then subtracted the filtered trace from the smoothed trace to isolate transient fluctuations within each component. These component fluctuations were then examined for correlations with movement along the x-axis, y-axis, or combined x-y displacement. Components that had a correlation coefficient >0.3 with at least one movement measure were considered movement-related. Peak detection was performed on these components to identify movement event timings. Peaks across all relevant components were merged to create a comprehensive movement event timeline for the session. For each identified movement event, corresponding segments of the calcium trace for each cell were replaced with interpolated values. To generate the interpolated trace, data were first smoothed with a low-pass filter, followed by cubic interpolation across the affected segment. If cubic interpolation produced unrealistic values, linear interpolation was used instead. To preserve realistic signal variability, Gaussian noise was added to the interpolated segments, matching the spectral properties of the original data. Specifically, we computed the Fourier transform (FFT) of each trial to extract its frequency components and magnitudes. The three largest non-zero frequency components were identified and used to generate noise at corresponding frequencies. Noise amplitude was scaled based on the minimum standard deviation of real (unaffected) data segments within the trial before being added to the interpolated trace.

#### Mean reward response per cell

For quantifying the mean reward response of individual cells, calcium traces for each entire day first underwent min-max normalization (1st to 99th percentile) to reduce the influence of outliers. We then selected the first 30 consumed reward trials. For each trial, the mean activity during the 2-second window following reward consumption was calculated. The mean of these values across trials was defined as the cell’s response to that specific reward condition.

#### Single cell analysis - stable responsive cells and optogenetic inhibition effect

As an initial approach to assess the overall single cell stability of the effects of decreased physiological needs, responsive cells were identified as those showing a significant change from zero in their trial-averaged response (one-sample t-test, α = 0.05) during either the first or last ∼40 trials of the session, on either of the two control days. For each responsive cell, we compared its mean response during the first 10 trials to its mean response during a matched 10-trial window following the light period (immediately after light onset, for short-term effects, or after light offset, for long-term effects), and classified the change as no change, stronger, weaker, or opposite-sign, based on the relative magnitude and sign of the two means. To retain only cells with a consistent classification, this procedure was applied independently to both control days, and a cell was included in the final analysis only if its classification was consistent between the two days (or one day showed no change and the direction of change agreed with the other day), cells with inconsistent classifications across the two control days were excluded from this analysis.

For quantification of the effects of optogenetic thalamic inhibition, we chose to use a stricter stability criterion to assess the effects of optogenetic inhibition only on the cells with the most stable responses, both qualitatively and quantitatively. For each responsive cell, we compared its mean response in a 10-trial window before the first light onset to a matched 10-trial window after the light period (short-term: immediately after light onset, long-term: after light offset), and classified the change as no change, stronger, weaker, or sign-reversed. However, a cell was only classified as “changed” if the end-window mean fell outside ±10% of the start-window mean and the absolute difference between the two exceeded a minimum threshold of 0.025 (min-max normalized ΔF/F). Critically, a cell was retained in the final analysis only if this classification, including both the response-type category and the sign of the effect (excitatory/inhibitory), was consistent across the two control days. Any mismatch, including cases where one day showed “no change” and the other showed a change, resulted in exclusion. Group-wise response magnitudes (start-end deltas) were compared across the three days using one-way ANOVA followed by Tukey’s post hoc test.

#### Single cells - comparing short- and long-term populations

To examine whether the same cells were affected similarly across short- and long-term timescales, we restricted the analysis to responsive cells (as described above) whose pre-light response additionally had a consistent sign across all three days. For each cell, we defined a control baseline as the signed average of its start-end response change across the two control days, and compared this to its start-end change on the inhibition day. Cells were categorized as “stabilized” if the magnitude of the inhibition-day change was smaller than the control baseline magnitude, or “changed” otherwise, irrespective of sign. Within each category, cells were further ranked by the deviation of their inhibition-day change magnitude from their own baseline, and the most extreme 30% of cells were selected for comparison between short- and long-term populations. We then quantified the overlap between short- and long-term cell sets (e.g., stabilized-short vs. stabilized-long, changed-short vs. changed-long, changed-short vs. stabilized-long) using a hypergeometric test, comparing observed overlap to the overlap expected by chance.

#### Single cells - distribution of long-term response changes

To assess the distribution of long-term changes in reward responses across all individual cells, we compared each cell’s mean response during the initial 10 trials to its mean response in a matched 10-trial window after the light period (after light offset), as described above. For each cell, we calculated a signed response change, defined as the difference between the end- and start-window means, multiplied by the sign of the start-window response, such that positive values indicated stronger responses and negative values indicated weaker responses, regardless of whether the response was excitatory or inhibitory. When the start-window response was negligible relative to the end-window response (less than 10% of its magnitude), the end-window response sign was used instead. This measure also accounted for sign reversals, which were assigned values indicating a reduction in response strength beyond complete response loss. All cells were included, and the distributions of response changes were compared across the two control days and the optogenetic inhibition day.

#### Correlation and slope

Neural responses were quantified using the “*Mean reward response per cell”* procedure, applied separately for water and food trials. To compare responses across two experimental conditions, we calculated the Pearson correlation coefficient and the slope of the linear regression between cell response scores for each mouse.

#### Correlation and slope while comparing different physiological states

Using the same correlation/slope approach described above, we additionally compared reward responses across the four-day imaging week (see “Imaging across different physiological states”). Specifically, we compared: water responses between the two water-restricted days (day 1 vs. day 2), food responses between the two food-restricted days (day 3 vs. day 4), water vs. food responses on the same day (day 1, when mice were water-restricted but still received food and neutral trials), and food responses between the two days on which food trials were presented (day 1 vs. day 3).

#### Correlation and slope after decrease in physiological need

To assess whether neural responses changed systematically across a session, we compared mean per-cell responses between trials from the start and end of the relevant day (water-restricted or food-restricted), using up to 30 trials from each period (capped at 30% of total trials per period if fewer were available). For food, “end” trials were taken from the additional session recorded after ∼4 hours of free food and water access (see above), rather than the end of the original session. For each cell, we computed the mean response across the start-window trials and across the end-window trials, and calculated the Pearson correlation coefficient and linear regression slope between these two sets of values across cells.

#### Correlation and slope after decrease in physiological need, while controlling for behavioral differences

To ensure that the results above were not confounded by differences in consumption-related behavior between the start and end of the session, we repeated this analysis using behaviorally matched trial subsets, following the same start/end trial selection described above. For each end-of-day trial, we computed a behavioral measure over a matched window around consumption onset, and identified start-of-day trials with the closest (selected from the first 30 trials of the session). End trials without a suitable match were excluded, and mice were included only if at least 6 end-trials had a match and at least 6 distinct start-trials contributed to those matches. Per-cell mean responses were then recalculated using only these matched trial subsets, and correlation and slope were recomputed as above. As an additional exclusion criterion, mice were retained only if their day1-day2 (start-to-start) correlation, used as a within-mouse stability reference, exceeded 0.3; all remaining measures were then normalized to this reference. This matching procedure was performed separately using five behavioral measures: lick count (±1 lick), anticipatory licking (±1 lick), lick latency (±100 ms), and lick-bout variability (standard deviation of inter-lick intervals; ±10% of the end-of-day value) for water, chewing-related orofacial movement (±10%) for food, and pupil diameter. (±10%; valid trials required at least 6 frames of good-quality pupil tracking within the response window). For chewing, the end-of-day condition consisted of trials from a separate sated session, recorded after 4 hours of free access to food and water, rather than the last trials of the same session as used for water. For pupil measurements in the food condition, start/end comparisons were instead made within the same session, since pupil size was not readily comparable between the sessions before and after the mouse was removed from the microscope for the ∼4 hrs home-cage consumption. As a reference, the same matching procedure was additionally used to compare start-of-day trials between two separate same-restricted days, rather than start vs. end of the same day, to establish a baseline level of correlation/slope expected in the absence of a state change.

#### Correlation and slope during optogenetic inhibition

Changes in Pearson correlation and linear regression slope due to optogenetic inhibition were evaluated during two phases: early and late effects. Trial selection was based on 10% of baseline trials for each reward type (five trials for food, seven for water). For each cell, the baseline period was defined as the mean response across the 5 (food) or 7 (water) trials preceding light stimulation onset. The early effect was calculated as the mean response during the first 5 (food) or 7 (water) trials following light onset. Late effects were assessed by comparing responses before light stimulation with those after light offset. For this analysis, twice the number of trials were selected post-illumination offset (10 for food, 14 for water). For each trial, the mean response during the 2-second window following reward delivery was calculated. Pearson correlation coefficients and regression slopes were computed for all three experimental days. Results were normalized to values obtained on day 2 (baseline). The primary comparison was between the normalized responses on day 1 and the optogenetic inhibition day (day 3).

#### Correlation of food responses during food restriction, with or without water

Using data from the four-day imaging week (see “Imaging across different physiological need states”), we tested whether the presence of water availability during the task affected the stability of food-related responses. We compared per-cell mean responses at the end of consumption across three conditions differing in water access: the end of day 3 (food-restricted, receiving water, food, and neutral trials-water available), the end of day 4 (food-restricted, receiving only food and neutral trials-no water available), and the start of the additional post-meal session on day 4 (referred to as “day 4b”, food and neutral trials only, following ∼4 hours of free water and food access, see above). For each pair, mean per-cell responses were computed over the last (or, for day 4b, first) time window in the relevant session, defined as one-third of the shorter of the two compared sessions (day 3 or day 4). Mice for which this window contained fewer than 15 trials were excluded from this analysis. Pearson correlation coefficients and linear regression slopes were calculated between conditions. We also computed start-vs-end correlations and slopes within day 3 and within day 4 individually. This allowed us to compare how much the population response naturally changes within a session, and to test whether this within-session change was similar regardless of water availability matching the finding that end-of-session responses were similar between day 3 and day 4 above.

#### Selectivity Index

The selectivity index was calculated based on the “*Mean reward response per cell”* for each condition, using the first 30 consumed trials per condition. The index was computed as:

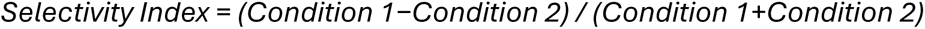

For comparisons involving water-water or food-food trials, day 1 was designated as Condition 1. For water versus food comparisons, water was assigned as Condition 1. Consequently, a positive index indicates a stronger response to Condition 1 / water. If the responses to the two conditions were opposite in sign, and one was at least 1.5 times smaller than the other, the smaller response was set to zero. Cells not meeting this criterion were excluded and classified as “opposite” cells. We observed that water-food comparisons exhibited significantly more opposite cells than same-reward comparisons. We quantified the width of selectivity index distributions by comparing the standard deviations of the distributions. For water-food comparisons, data from experiments containing both water-only and food-only days within the same week were used. For water-water and food-food comparisons we used separate weeks consisting of two days with water-only (thirsty state) and two days with food-only (hungry state).

#### Cosine similarity

To assess long-term changes in decreased physiological need at the population level, we first calculated the mean response of each cell for each consumed reward trial on days when only water or food were available, as described in “*Mean reward response per cell”.* Each cell’s responses were smoothed with a two-trial window. For each cell, we then computed the mean response across the first 15 trials, forming a baseline population vector. Cosine similarity was calculated between the response vector for each reward trial and this baseline.

#### Cosine similarity change during optogenetic inhibition

This analysis followed the same procedure as above, with trial phases and selection criteria as described in the “*Correlation and slope during optogenetic inhibition*” section. In addition to examining long-term changes (before and after the illumination period), we also examined changes during light stimulation by comparing the first 5 (food) or 7 (water) trials after light onset and the 10 (food) or 14 (water) trials before light offset. The resulting change in cosine similarity (delta) was normalized to day 2, with the primary comparison performed between normalized days 1 and 3.

#### Change in cosine similarity of reward responses when both rewards were available

We applied this analysis to food-reward responses during food restriction, comparing day 3, when both food and water were available, with day 4, when only food was available. The change in cosine similarity was quantified by subtracting the mean cosine similarity of the last subset of trials from that of the first subset, using 20 trials or 25% of the trials per session, whichever was smaller. Only the first session of day 4 was included, to match day 3, which had no additional session. We performed the same analysis for water-reward responses during water restriction, comparing day 1, when both food and water were available, with day 2, when only water was available.

#### Dimensionality reduction and clustering

We followed our previously published pipeline^57^. Briefly, data were binned into half-second bins and then subjected to min-max normalization using the 5th and 95th percentiles. Dimensionality reduction was performed in two stages: first reducing to 20 dimensions for denoising, then further reducing to 6 dimensions - the latter determined as the optimal dimensionality for our data (typically 5-6 intrinsic dimensions). Clustering was subsequently applied as previously described^57^, clustering the data into eight clusters. This allowed us to assign each time point to a specific cluster and describe population dynamics as cluster transition dynamics.

#### Concatenating two days for combined manifold analysis

The same approach as above was used for combining data across two days, with modifications to account for the longer recording duration. Specifically, data were binned into 1-second intervals. To improve the alignment of neuronal activity across days, the data underwent min-max normalization followed by z-scoring after the min-max normalization. This enabled direct comparison of neural activity structures between different physiological states or optogenetic conditions and the control day.

#### Similarity of manifold cluster dynamics

To assess the similarity *of* manifold cluster dynamics between two conditions (e.g., water vs. food trials), we used two different methods.

For the first method, we analyzed the temporal distribution of cluster assignments. Trials from each condition were aligned by cue onset, and only clusters present in at least 20% of trials in at least one condition were included, excluding the central point cloud cluster. An equal number of trials were selected per condition. For each included cluster, we computed the time-resolved percentage of trials in which that cluster appeared, resulting in a temporal distribution profile per cluster. Clusters that appeared in fewer than 10% of trials were assigned values of zero to prevent noise-driven similarity. The similarity between conditions was quantified by calculating the cosine similarity for each cluster’s distribution across conditions. The final similarity score was the average of the cosine similarity values across all included clusters.

For the second method, we examined transition probabilities between clusters. We calculated the probability of transitioning from each cluster to every other cluster within trials, considering all data points. Self-transitions (remaining within the same cluster) were excluded, as they were inherently frequent and did not provide meaningful information for this analysis. Transitions with fewer than three occurrences were set to zero. The resulting transition matrix was normalized by dividing all entries by the maximum value within each matrix. To compare two conditions, the transition matrices were flattened into one-dimensional arrays, and their similarity was quantified using the Pearson correlation coefficient. This analysis yielded results that were consistent with the first method.

#### Comparing optogenetic inhibition manifold with control manifold

To test whether optogenetic inhibition altered the manifold structure, we quantified the similarity of manifold cluster dynamics before and during light illumination using the method described above. Specifically, for each mouse, we compared distributions during the baseline period to those during the light period on the optogenetic inhibition day (Day 3). As a control, the same comparison was performed for the equivalent periods on the previous day without optogenetic illumination (Day 2). The structure change due to optogenetic inhibition was then evaluated by comparing Day 3 to Day 2 results.

#### Unique clusters analysis

To further quantify the optogenetic effect, we assessed the appearance of “unique clusters” - clusters that emerge during the light period but are absent in the baseline period. As the prevalence of specific clusters naturally changes over time (e.g., due to decreased physiological need^57^), some cluster differences between early and late periods occur even in control conditions. Therefore, we compared the percentage of unique clusters appearing during the light period on the optogenetic day (Day 3) with the same period on the control day (Day 2). A significantly higher occurrence of unique clusters on Day 3 would indicate that optogenetic inhibition induced new or altered activity patterns, not present under normal conditions.

#### Dimensionality reduction - Isomap

To confirm that our results were not dependent on the specific dimensionality reduction method used, we repeated the manifold comparison between control and optogenetic inhibition days (see “Comparing optogenetic inhibition manifold with control manifold”) using Isomap instead of Laplacian eigenmaps. Data were denoised to 100 dimensions using PCA, then reduced to 6 final dimensions via Isomap (70 neighbors).

#### Pair-wise correlation matrix similarity

To assess whether optogenetic inhibition altered the correlation structure between cells, we computed pairwise cell-cell correlation matrices separately for the period before the optogenetic window (baseline) and during the optogenetic window (test), on the second control day and the inhibition day. For each day, we compared the two matrices by correlating their upper triangles (Pearson correlation) between baseline and test, yielding a single stability score per day, per mouse, where a high correlation indicates that pairwise cell-cell relationships were preserved between the two periods, and a low correlation indicates that the correlation structure reorganized.

#### Estimation of InsCtx ongoing activity

To estimate ongoing activity levels, we used our previously established method^13^. Specifically, we selected time periods that are minimally influenced by task performance or licking behavior: the final 3 seconds of inter-trial intervals (ITIs) from neutral trials, provided they were preceded by at least 6 seconds without licking. Due to task structure, the previous rewarded trial occurred approximately 45 seconds before the selected ITI segment, reducing potential task-related carryover effects. To enable comparisons across days, each neuron’s activity trace was min-max normalized using the 1st and 99th percentiles for that day.

#### Hunger/thirst axes definition

We defined the “need axis” by calculating the difference between the average population activity during the thirsty/hungry state and the quenched/sated state. Specifically, the thirsty state was defined as the inter-trial intervals (ITIs) preceding the first 20% of consumed rewards, while the quenched state was defined as the ITIs following the last 20% of consumed rewards. For food, sated-state ITIs were taken from the additional session recorded after ∼4 hours of free food and water access (see above), rather than from the end of the original session. The ongoing population activity was then projected onto this axis by calculating the dot product between the moment-to-moment activity patterns and the defined “*need axis*”. To compare across mice, projection values were linearly scaled so that activity corresponding to the thirsty state was assigned a value of 1, and activity corresponding to the quenched state was assigned a value of 0. Intermediate values reflected transitional activity states between thirst/hunger and satiety.

#### Relationship between ongoing activity and reward-response

To examine the relationship between ongoing InsCtx activity and reward-related responses, we used single-cell responses calculated as described in the “*Mean reward response per cell”* section. We then computed the correlation between these reward-evoked responses and ΔState, defined as the difference in mean ongoing activity between physiological states (thirsty-quenched or hungry-sated).

#### Estimation of axis stability and similarity across days and states

To assess the stability of defined need state axes (thirst and hunger axes) across days, we calculated each separately for two consecutive days in which mice had access only to water or only to food, and were in a thirsty / hungry state accordingly. As cells were aligned across days, we could project ongoing activity from one day onto the axis defined on the other day. We quantified movement along the axis by calculating the change in projected activity between the start and end of each day. The start and end periods were defined as the 20 ITIs (or 20% of total ITIs) with the lowest need axis values, representing transition points in physiological state. For each mouse, the two projection measurements (Day 1 activity on Day 2 axis and vice versa) were averaged to obtain a measure of axis stability. When assessing hunger axis stability, a 1-day gap was introduced between the two sessions, as longer food restriction was necessary to re-establish the hungry state. To further validate the specificity of these axes, we projected ongoing activity from thirst days onto the hunger axis, and from hunger days onto the thirst axis, using sessions separated by at least one day. Minimal movement along the mismatched axes indicated that the thirst and hunger axes reflect distinct, independent physiological states rather than general across-day variability.

The correlation between state-axes was calculated using the Pearson correlation coefficient, applied to the previously defined axes describing different physiological states.

#### Movement along reward-specific axes when both rewards were available

To test how the availability of both rewards affected changes in internal-state representations, we projected neural activity from day 1, when the mouse was thirsty and both water and food were available, onto the thirst axis defined on day 2, when the mouse was thirsty and only water was available. We quantified the change between the initial and final portions of the session using inter-trial intervals following water consumption, and compared it with the corresponding change measured across two days when only water was available on both the axis-definition day and the activity-measurement day. Projections were normalized to the mean projection during the first 10 ITIs of each day.

Similarly, we projected day 3 activity onto the hunger axis defined on day 4, comparing changes following food consumption when both rewards were available with those measured across two days when only food was available. Projections were normalized in the same way.

#### Estimation of transitions along the need axis during optogenetic inhibition

For optogenetic inhibition days, need axes could be accurately determined using the same day’s activity as activity at the end of the day might be affected by the optogenetic inhibition. Therefore, to circumvent this issue we performed an across-day comparison during the 3-day experiment. We used the axis from day 2 and projected onto it the activity from day 1 (control day) and day 3 (optogenetic inhibition day). This allowed us to test how the need axis is affected by optogenetic inhibition while controlling for across-day variability in activity. Specifically, ongoing activity was projected onto this axis separately for food and water conditions, and the resulting dot product values were normalized by dividing by the mean of the first 10 ITIs. We then quantified changes in axis position by comparison of the 9 ITIs before light onset to the 9 ITIs after light offset. For each phase, the mean projected activity was calculated, and the difference between phases was used to quantify the effect of optogenetic inhibition on need-state axis position.

#### Pupil size changes on inter-trial intervals during optogenetic inhibition

We quantified the change in pupil size during ITIs, used for assessing ongoing activity changes, before and after optogenetic inhibition in order to assess the effects of arousal. We tested the same windows as in the “*Estimation of axes transition during optogenetic inhibition”,* 9 trials before and after optogenetic inhibition. We required a minimum number of 6 trials that passed pupil analysis quality controls (at least 6 good frames per ITI). We therefore did not have a sufficient number of mice to analyze the effect of optogenetic inhibition on pupil size during food restriction.

#### Classification of response-types

To identify distinct response types among InsCtx neurons, we sought to establish a procedure that would be as conservative as possible in terms of classifying responses as different across conditions. That is, a method that would be biased to classify responses as similar whenever possible, thereby reducing false positive identification, e.g., of prediction errors.

#### Amplitude and shape similarity estimation

We performed experiments using three trial types: reward, omission, and surprise. These experiments were conducted on days when mice were either thirsty and received only water or hungry and received only food. To quantify similarity in response shape and amplitude, all neuronal activity traces were min-max normalized (99th to 1st percentile) and smoothed using a 330 msec window to minimize fluctuations that could bias the analysis. We selected trials from comparable time windows, specifically the first 60 consumed reward trials and all relevant omission or surprise trials in this time period. For water trials, this included trials where the animal licked following the cue for both reward and omission trials. For food trials, we used all omission trials, which often lacked anticipatory licking, and all surprise trials in which the animal consumed the reward. For each cell, we subtracted baseline activity from the 2 seconds preceding cue onset, averaged responses across trials, and generated a trial-averaged response for each condition.

To assess the natural trial-to-trial variability of each cell, we performed 500 permutations where, in each permutation, we randomly selected two groups of reward trials (matched in number to the smaller number of omission or surprise trials), calculated the Pearson correlation between the two averaged responses, and recorded the maximum and minimum amplitudes from the first group. This provided a distribution of expected variability of reward responses for both shape and amplitude within each cell. We then calculated the Pearson correlation between the mean response to omission or surprise trials and the mean response to reward trials. Trials were classified as similar in shape if the correlation fell within the mean ± 2*STD of the reward trial variability distribution or if the correlation exceeded 0.8. For direct comparisons between omission and surprise trials, shape similarity was defined by a correlation exceeding 0.8. Amplitude similarity was assessed by first determining whether the reward response peak was positive or negative and requiring the same sign for omission or surprise trials to be considered. We then calculated the mean response within a 250 msec window before and after the peak and classified responses as similar in amplitude if they fell within the mean ± 2*STD of the reward response amplitude and within a 50% to 150% range of the reward response. When comparing omission to surprise responses, amplitude similarity was defined as one peak falling within 50% to 150% of the other.

To account for variability in response timing, the entire procedure was repeated with different alignment points. For water trials, responses were aligned to the first lick, reward consumption, and lick offset. For food trials, responses were aligned to the first lick, cue offset, and reward consumption. Each aligned window was 5 seconds long, starting one second before the alignment event. To minimize confounds from earlier activity, particularly for lick offset alignments, the first second of each response was subtracted in addition to the baseline correction described above. This approach allowed for robust classification of response types across different trial conditions and alignment strategies. We note that in our Pavlovian task with a constant cue duration of 2 sec, water reward is delivered immediately at cue offset. Similarly, food delivery is triggered at cue offset but then takes 1 sec to reach the “pellet cup”. Therefore, the same alignment can be used to capture neuronal responses to any of these events, but with a constant 2-sec delay for cue onset and cue offset with water reward delivery, as well as a constant 3-sec delay for cue onset and cue offset with food reward delivery. We used this alignment only to optimally compare across different experimental conditions without ascribing the neuronal response specifically to any of these events at this point. We emphasize that this alignment procedure was intentionally conservative in that it was biased to find the maximal similarity between all three conditions (expected, omitted, and surprise rewards). Thus, it is biased to identify responses are “consummatory action” (licking/chewing) or as “consummatory” (e.g., taste), and might underestimate the prevalence of other response types

#### Alignment for surprise trials

In surprise trials, the animal naturally did not anticipate the reward, and therefore they lacked anticipatory licking. As such the licking pattern typically differed from omission and reward trials that did usually have anticipatory licking. To ensure that any observed dissimilarity between surprise trials and other trial types was not simply due to a temporal shift in the neural response, we performed a cross-correlation analysis between the mean response of surprise trials and reward trials to identify the optimal lag. Specifically, we allowed for a temporal shift of up to two seconds earlier in the surprise trials to identify the shift that produced the highest correlation between the two conditions. Once the optimal shift was determined, we used this adjusted surprise response to compare with omission trials as well. As noted above, this procedure is biased toward finding greater similarity between reward and surprise responses. However, given that our goal is to rigorously test for genuine prediction and prediction-error signals, this approach applies a conservative, more stringent criterion to the analysis.

#### Responses significance

To determine which cells exhibited significant responses, we first assessed whether their activity reflected genuine signal rather than noise. Importantly, we applied relatively lenient at this step, as this measure was used only for identifying significantly responsive cells in conditions with relatively few omission or surprise trials, where variability is high. This classification was exclusively applied within the cell-type analysis. All neuronal activity was min-max normalized (99%-1%) and smoothed using a 100 msec window. We analyzed the first 50 consumed reward trials, all consumed surprise trials within the same period, and, for the omission trials within the same period, all omission trials with licking for water sessions, or all omission trials for food sessions. Trials were aligned to reward consumption for both reward and surprise trials, and to either the first lick after cue onset (water) or cue offset (food) for omission trials. For each trial, we compared the baseline period (2 seconds before cue onset) to the mean response during the 5 seconds following the alignment point using a paired t-test. If the response was significantly different from baseline, the cell was classified as significantly responsive to that trial type. If no significant response was detected over the full 5-second window, we performed a finer-grained analysis to capture shorter or behaviorally driven responses that may have been missed. Specifically, we divided the post-onset period into 300 msec windows, with windows sampled every 300 msec. We compared each window to the baseline response using a paired t-test, applying Bonferroni correction for multiple comparisons. If at least three consecutive windows (representing one second of activity) had a significant difference from baseline, the cell was classified as significantly responsive to that trial type.

#### Classification of cell-types

As described above, we aligned neuronal responses to different behavioral phases to account for variability in response timing across cells. For each cell, we selected the alignment phase that provided the highest similarity between conditions (reward, omission, or surprise). Two conditions were considered similar only if they exhibited significant similarity in both response shape and amplitude, with the additional requirement that at least one of the conditions showed a significant response (we required only one significantly responsive condition due to the high variability often seen in omission and surprise trials).

Our priority in selecting the optimal phase was to maximize the number of similarities identified between conditions. If all alignment phases produced the same similarity pattern, we selected the phase with the highest shape similarity. In cases where different phases produced different combinations of similarities between conditions, we prioritized the phase with the greatest number of condition pairs classified as similar. If multiple phases showed the same number of similarities but between different condition pairs, the phase with the highest shape similarity was selected. If no phase produced similarity in both shape and amplitude, we relied on shape similarity alone, applying the same prioritization: more similarities across conditions were prioritized, and when multiple options remained, the phase with the highest shape similarity was chosen. Based on these comparisons, cells were categorized as follows: (1) If all three conditions (reward, omission, and surprise) were similar, the cell was classified as responsive to consummatory actions, indicating that its response was unrelated to prediction or consumption-specific processes. (2) If only reward and omission responses were similar, the cell was classified as having a prediction response, meaning its response was independent of reward consumption. (3) If only reward and surprise responses were similar, the cell was classified as consumption-related, suggesting its response was independent of expectation. (4) If only omission and surprise responses were similar, the cell was categorized as encoding a general prediction error. (5) In cases where omission or surprise trials had a similar response shape to reward but exhibited either significantly higher or lower response amplitude, these were classified as enhanced or suppressed prediction errors, respectively. (6) If one condition (omission or surprise) showed an enhanced prediction error and the other a suppressed prediction error, the cell was classified as encoding a signed prediction error. (7) If no condition pairs exhibited similar shapes, but a significant response was detected in either omission or surprise trials, the cell was classified as encoding a specific prediction error. Extremely rare cases with a response only to reward trials, without similarity to omission or surprise conditions, were excluded from the analysis due to their extremely low prevalence. Cells that exhibited multiple response patterns (for example, both consumption-related and enhanced prediction error) were allowed to belong to more than one category. Cells that showed no similarity in shape across conditions and no significant response were classified as non-responsive.

#### Controls for behavioral effects on enhanced and suppressed prediction errors

Enhanced and suppressed prediction errors were defined as responses with similar shape to the reward response but with significantly higher or lower amplitude, respectively. Specific prediction errors were defined as responses present during the prediction-error trial type (omission or surprise) but largely absent from the corresponding reward response, i.e., a response driven by the prediction error itself rather than an amplified or attenuated version of the reward response. However, these differences could potentially be explained by variations in behavior between trial types. To control for this, we performed a behavior-matching analysis for each cell and each relevant comparison (reward vs. omission or reward vs. surprise). This matching was performed separately using several behavioral measures: each with its own matching threshold: lick rate (±1 lick), lick latency (onset of licking; ±100 ms), lick-bout variability (standard deviation of inter-lick intervals; ±10% of the prediction-error-trial value), anticipatory licking (±1 lick), chewing-related orofacial movement (food; ±10% of the prediction-error-trial value), and pupil diameter (water and food; ±10% of the prediction-error-trial value; valid trials required at least 6 frames of good-quality pupil tracking within the response window).

For each measure, the relevant behavioral value was calculated in a 2-second window following the relevant behavioral event, specific to each alignment phase, with the exception of anticipatory licking, which was measured during the cue period. Lick latency and anticipatory licking were used only to control omission comparisons for cells with omission prediction error responses. This is because surprise trials are by design unanticipated, so there is no anticipatory period for the mouse to lick in. Additionally, lick latency is not a meaningful measure in this context since responses are aligned to reward consumption. Chewing-related orofacial movements were used only as controls for surprise comparisons for cells with surprise prediction error responses, since no food is consumed on omission trials and there are no chewing movements to match. For each prediction error trial, we identified reward trials with a matching behavioral value for that measure, requiring at least 3 prediction-error trials with a matching reward trial for that cell to be included in the analysis using that measure. Cells with fewer matched prediction error trials were excluded from this analysis. This provided a set of behavior-matched reward trials for direct comparison. In addition, for each mouse and each prediction error cell group (enhanced, suppressed, or specific), a group-level result was only computed if at least 7 cells passed the per-cell matching criteria for that measure; mice with fewer qualifying cells for a given group were excluded from that group’s summary analysis. For each cell, we calculated the mean response (across trials) for both the prediction error trials and the corresponding matched reward trials, using the previously selected alignment phase.

For all behaviorally-matched cells, we identified the peak of the response for both conditions and computed a modulation index based on these peak values, cells for which the reward and prediction-error peaks had opposite signs and neither was at least 1.5-fold larger in magnitude than the other were excluded from the modulation index calculation, as the direction of modulation could not be quantitatively assigned. This process was also performed on the original (uncontrolled) data to allow comparison. The modulation index was averaged across mice, with a value of +1 indicating a strong prediction error response and a value of -1 indicating a stronger reward response.

#### Effects of decreased physiological need on different response-types

We compared effects of decreased physiological need across different response categories, specifically “consummatory actions”, “consumption”, and “predictions”. For each cell within these groups, we quantified changes associated with decreased physiological need by analyzing changes in reward responses across the session. We selected the first 30 and last 30 reward trials for each cell and calculated the mean activity within a 2-second window around the relevant alignment phase for each trial. Cells from all mice were pooled together for group-level comparisons. To compare decreased physiological need dynamics between groups, we computed Pearson correlations for each group and compared them using Fisher z-transformation followed by a z-test for comparing correlation coefficients. Additionally, we estimated the linear regression slopes for each group using ordinary least squares (OLS) regression with interaction terms to test for differences in trends associated with decreased physiological need. A similar approach was applied to prediction error responses. For omission and surprise trials, we compared the groups “consummatory actions”, and “prediction-errors” (the latter including all cells exhibiting relevant prediction error responses (omission or surprise)). For these analyses, we selected the first and last 4-5 trials per session (maximum of 5 trials, or 4 when fewer were available). The same correlation and linear regression methods were applied to assess group differences in response dynamics. This analysis was conducted only for water trials, as the number of relevant food trials was insufficient to reliably compare early and late-session responses.

#### Classification of response types by reward and omission trials

This classification was performed in the context of the optogenetic inhibition experiment. The approach is similar to the method described above, but only comparing reward and omission trials. The analysis was performed separately for each day, using only trials from the baseline period (prior to the onset of optogenetic inhibition). We used the same procedures described above to assess the similarity of response amplitudes and shapes between reward and omission trials.

#### Responses significance for optogenetic experiment

To determine whether cells responded to reward or omission trials during the optogenetic inhibition experiment, we examined multiple trial selections. Because optogenetic inhibition might alter cell responses or evoke specific activity patterns, we tested response significance in two separate phases: the first 40 trials of each session and the light illumination period. We analyzed two alignments: licking onset and reward consumption, independently for each day. If a cell showed significant response in either phase, it was classified as having a significant response. Significance was assessed by comparing activity during the 2 seconds before cue onset to the activity following cue offset, using the same method described above.

#### Classification to response types

Cells were categorized as having either similar responses to reward and omission (“similar”), different responses (“different”), or no significant response (“no response”). As described above, we prioritized the phase with the highest shape similarity for each cell. At least one of the compared trial types had to show a significant response to be considered as responsive. Cells showing different response shapes, or similar shapes but significantly different amplitudes, were classified as having “different responses.” Cells responding significantly to reward but not omission, without sufficient similarity between the two responses, were excluded because we could not determine whether these represented genuine reward-specific responses or reflected insufficient statistical power to detect an omission response.

#### Long-term optogenetic inhibition effects on different response types

For each group (“similar” or “different”), we calculated the mean response before and after the light period. Specifically, we used the first 5 trials for water and 7 trials for food (approximately 10% of the baseline) as the pre-light period, measuring the mean activity over the 2 seconds following the relevant alignment phase. This was compared to the post-light period, defined as 10 trials for water and 14 trials for food after the light was turned off. Cells from all mice were pooled together for analysis. We then calculated the correlation and regression slope between these groups. For visualization, the mean values from the two control days were displayed.

#### Topography of responses

Spatial organization was analyzed by combining cell coordinates across all mice. For each mouse, the FOV was rotated and cell coordinates were recalculated based on measured shifts relative to bregma, yielding common location coordinates for all cells.

#### Spatial organization of water/food response and of different response types

For each cell, the response was determined as described above. The anatomical coordinates along the posterior-anterior (x-axis) and dorsal-ventral (y-axis) axes relative to bregma were divided into 20 bins. Within each bin, we separately calculated: (1) the mean normalized response of all cells (within-mouse, normalized to the 90^th^ percentile), (2) the percentage of cells belonging to each cell type within each section, relative to the total cells in that section. Correlations (R) between these mean bin scores and the anatomical location of the bins were then calculated.

#### VPM/Lpc→InsCtx inhibition during quite-wakening

We classified neurons as affected by VPM/Lpc→InsCtx inhibition if their activity (z-scored based on the pre-illumination period) following light onset was above the 95^th^ percentile (excitation) or below the 5^th^ percentile (inhibition) of the activity distribution in the control (no light) condition.

### Fiber photometry data analysis

#### Pre-processing

A median filter was applied to reduce noise. Bleaching correction was performed by fitting and subtracting an exponential decay function. Motion artifacts were corrected by linearly fitting the isosbestic control signal to the activity-dependent signal; the resulting estimated motion component was then subtracted from the activity-dependent channel^106^. To quantify changes relative to baseline for each trial, the mean signal amplitude during the 2 seconds preceding cue onset was subtracted from the trial data.

#### NAc dopamine cue responses

The first 40 trials were selected, and the signal was normalized using the 1st and 99th percentiles. The mean signal across the entire cue window (2 seconds) was calculated to determine its polarity (positive or negative), which guided peak direction. The final response value was defined as the mean of a 0.5-second window centered around the peak.

#### NAc dopamine reward, omission, and surprise responses

For the analysis of reward, omission, and surprise responses, only the activity following the cue offset was considered. Each signal was normalized per mouse and per day to 95% of the highest response recorded that day. To correct for baseline activity, the mean of the first five sampling points after cue offset was subtracted from each trial. The resulting traces were then smoothed using a 10-point sliding average to allow for peak detection. In reward trials, the mean response was calculated similarly to the cue response method, but applied across all trials and using a 1-second window. For surprise trials, where responses were consistently positive, the peak was defined as the maximum value, and the final response was computed as the mean signal beginning from the time point at which the signal reached two-thirds of this peak. In omission trials, if the mean response dropped more than 1.65 standard deviations below the reward mean, the peak was defined as the minimum value; otherwise, the maximum was used. The final value was then derived from the mean signal starting at the point where the trace reached two-thirds of the identified peak.

#### InsCtx GCaMP fiber photometry data analysis

On inhibition days, optogenetic illumination was delivered following the baseline period (see above), overlapping with the post-injection recording window. Each session’s z-scored signal was compared between the pre-injection baseline (minutes 6-9) and a 5-minute post-injection window (minutes 10-15), with the response magnitude defined as the difference between the two. We observed mouse-specific variability in the direction of the responses (i.e., increase/decrease relative to baseline) that was consistent per condition, likely reflecting the fact that the fiber photometry method averages signals across a large heterogeneous cortical population. Thus, to allow averaging across conditions and mice regardless of the mouse-specific response direction, values for all days were sign-flipped per mouse based on the control day only such that the control-day response was always positive and other experimental conditions were not constrained. Control and inhibition responses were compared using paired t-tests, both in raw units and as a normalized ratio (inhibition/control).

### Electrophysiology data analysis

We downsampled LFP signals to 250 Hz by averaging consecutive samples, removed 50-Hz line noise with a zero-phase IIR notch filter, and band-pass filtered the signals from 0.1 to 100 Hz using a fourth-order Butterworth filter with zero-phase filtering. We estimated power spectral density using Welch’s method with 1-s windows, 50% overlap, and 1-Hz frequency resolution, and expressed power in decibels. We calculated time-frequency representations using non-overlapping 60-s windows with 1-Hz frequency resolution. We converted spectrogram power to decibels, standardized it within each frequency across time, and then smoothed the standardized spectrograms using a two-dimensional Gaussian filter. We smoothed group spectra over 15 frequency points and displayed them as the mean ± SEM. We used paired two-sided t-tests to compare frequency-band power between baseline and stimulation periods.

### Statistics

All statistical analyses were performed using standard Python libraries (SciPy, statsmodels). For pairwise comparisons between two conditions, we used paired t-tests (one-or two-tailed, as appropriate). When comparing more than two related conditions, we performed pairwise paired t-tests between all conditions of interest, with resulting p-values corrected for multiple comparisons using the Holm-Bonferroni method. For analyses pooling cells across animals, Pearson correlation coefficients were compared using Fisher’s z-transformation followed by z-tests. Linear relationships were assessed using ordinary least squares (OLS) regression models, incorporating interaction terms where applicable

---

**Ext. Data Fig. 1:**
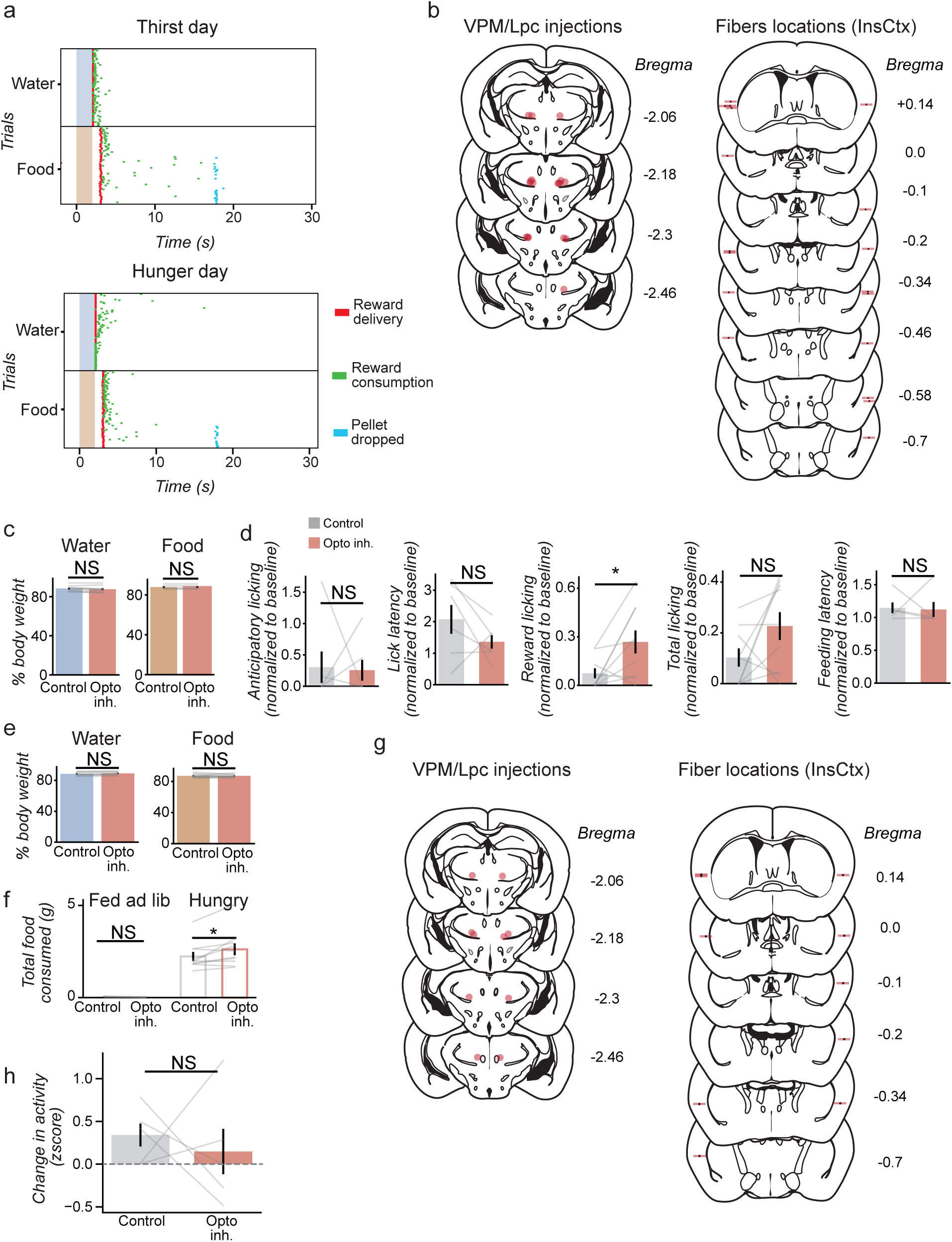
Additional analyses and controls for the effects of VPM/Lpc→InsCtx inhibition on behavior. a. Behavioral events from an example mouse for reward delivery (red ticks), reward consumption (green ticks) and pellets drops (blue ticks, dropping through the “trapdoor” if food rewards are not eaten). Colored rectangles: cues. b. Coronal brain section schematics, showing histological verification for experiments in Fig. 1e-i. *Left*: bilateral injection sites of AAV5-mCaMKIIa(short)-eOPN3-mScarlet in VPM/Lpc (Bregma −2.06 to −2.46 mm). *Right*: bilateral optic fiber placements in the InsCtx (Bregma +0.14 to −0.7 mm). n = 9. c. Percent of ad lib body weight during control versus optogenetic inhibition days, for water-restricted (*left*, mean±SEM, n = 9) and food-restricted (*right*, mean±SEM, n = 5) mice, corresponding to experiments in Fig. 1e-g. No significant difference between control and opto days for either restriction type, indicating that observed effects cannot be attributed to differences in body weight. d. Comparison of behavioral variables between control and optogenetic inhibition days, normalized to each mouse’s own baseline. Variables for water: anticipatory licking, lick latency, licks during reward consumption (5 sec following cue offset), and all trial licks (all licks from cue offset to the end of the trial). Variables for food: latency of food consumption. All comparisons were not significant except licks during reward consumption (*p = 0.016). water n = 9, food n = 5. mean±SEM. e. Percent of ad lib body weight during control versus optogenetic inhibition days, for water-restricted (*left*, mean±SEM, n = 8) and food-restricted (*right*, mean±SEM, n = 10) mice, corresponding to experiments in Fig. 1e-g. No significant difference between control and opto days for either restriction type, indicating that observed effects are not attributable to differences in body weight. f. Effects of bilateral optogenetic inhibition of VPM/Lpc→InsCtx axons on home-cage free consumption of food in ad lib fed mice (*left*) in comparison to food restricted mice (*right*, same as in Fig. 1i). mean±SEM, n = 5. g. Coronal brain section schematics, showing histological verification for experiments in Fig. 1j-l. *Left*: bilateral injection sites of AAV5-mCaMKIIa(short)-eOPN3-mScarlet in VPM/Lpc (Bregma −2.06 to −2.46 mm). *Right*: bilateral optic fiber placements in the InsCtx (Bregma +0.14 to −0.7 mm). n = 5. h. No effect of bilateral optogenetic inhibition of VPM/Lpc→InsCtx axons on InsCtx activity during sham injections.

**Ext. Data Fig. 2:**
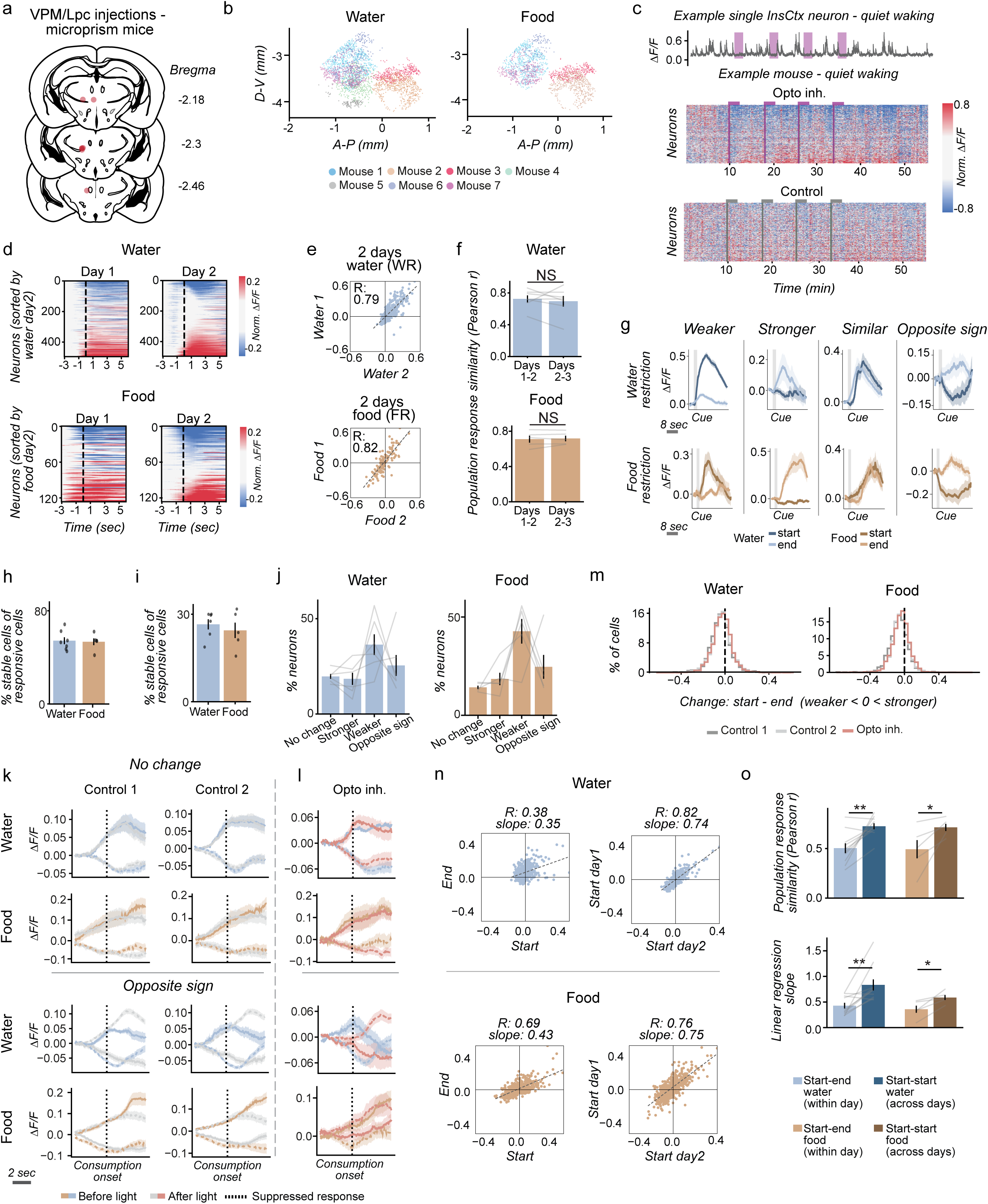
Additional analyses and controls for the long-term effects of VPM/Lpc→InsCtx inhibition on InsCtx food/water responses. a. Coronal brain section schematics of histological verification of optogenetic inhibition experiments in Fig.2–4. Unilateral injection sites of AAV5-mCaMKIIa(short)-eOPN3-mScarlet in VPM/Lpc (Bregma −2.18 to −2.46 mm). n = 7. b. Anatomical location of recorded InsCtx neurons along the dorsoventral (D-V) and anteroposterior (A-P) axes, as imaged through the microprism, for water (*left*) and food (*right*) sessions, corresponding to optogenetic inhibition experiments in Fig. 2–4. Each point represents one neuron. Colors indicate individual mouse. n = 7 mice. c. Example of neuronal activity during optogenetic inhibition of VPM/Lpc→InsCtx axons in the InsCtx during quiet wakefulness (no behavioral task). *Top:* Example activity trace of a single neuron showing reduced activity during optogenetic inhibition (purple bars indicate illumination periods). *Bottom:* Heatmaps showing population activity from one mouse across days, comparing control (no light) and optogenetic inhibition (red light) sessions. Vertical lines mark the onset of each 2-minute inhibition period. n = 334 neurons. d. Neuronal responses from an example mouse to food and water rewards on two days, during food or water restriction. Neurons are sorted by the second day (*right*), separately for food and water. Dashed line: consumption onset. e. Example population response similarity (Pearson correlation) between two days of reward consumption from one mouse, for water (*top*) and food (*bottom*) during the relevant physiological need. f. Quantification of population response similarity (Pearson correlation) across 3 days of for food or water during the relevant physiological need. Thirst: 381±174 neurons per mouse; mean±SEM, n = 10 mice. Hunger: 325±131 neurons per mouse; mean±SEM, n = 5 mice. g. Example single-neuron responses illustrating the types of changes observed as physiological need decreases, comparing responses at the start versus end of the session. From left to right: enhanced response (“stronger”), decreased response (“weaker”), reversed response direction (e.g., from suppression to excitation, “opposite sign”), and no change. Water (*top*) and food (*bottom*). Gray shaded rectangles indicate the cue period. h. Percent of responsive neurons with consistent changes with decreasing physiological need (i.e., changing in the same direction) across the two control days, for water and food during water and food restriction, respectively. Dots represent individual mice. Water n = 6 mice, 171±75 neurons per mouse, food n = 5 mice, 135±68 neurons per mouse; mean ± SEM. i. Percent of responsive neurons with consistent changes with decreasing physiological need under stricter criteria, used to quantify response magnitude changes in the optogenetic inhibition experiments. These neurons that remained in the same change-type group (stronger/weaker/opposite sign/no change) and showed a similar magnitude of change across the two control days, for water (n = 6 mice, 88±33 neurons per mouse) and food (n = 5 mice, 62±35 neurons per mouse); mean ± SEM. j. Percent responsive neurons classified into each change-type group (no change, stronger, weaker, opposite sign) within a single control day, for water (*left*, n = 6 mice, 282±107 neurons per mouse) and food (*right*, n = 5 mice, 204±80 neurons per mouse). Across all mice, mean ± SEM. k. Change in InsCtx average group responses with a decrease in physiological need in two control days (Control 1, Control 2), for “no change” and “opposite sign” groups, pooled across all mice. Only responsive neurons showing a stable change across both control days were included. Solid lines: excitatory responses, dashed lines: suppressed responses. Blue (water responses in water restricted mice. n = 6 mice, no change-78 neurons, opposite sign-108 neurons) and brown (food responses in food restricted mice. n = 5 mice, no change-27 neurons, opposite sign-66 neurons): responses ∼25 min into the session. Gray: responses after ∼1.5 hrs of consumption. Dotted vertical line: consumption onset. l. Same neurons as in ‘k’, but during the day with optogenetic inhibition of VPM/Lpc→InsCtx axons. Blue (water) and brown (food): responses ∼25 min into the session, before optogenetic inhibition. Red: responses after optogenetic inhibition. Solid lines: excitatory responses, dashed lines: suppressed responses. Dotted vertical line: consumption onset. m. Distribution of the change in reward responses across all neurons, before versus after the illumination period, for water (*left*) and food (*right*), on the two control days and the optogenetic inhibition day. Change values were signed relative to the direction of the starting response: 0 indicates no change, positive values a stronger response and negative values a weaker response, for both excited and suppressed neurons. Neurons whose response reversed sign are scored as weaker. Histograms show the percentage of neurons per bin, pooled across mice. Dashed line: no change. Water: n = 7 mice, 2438 neurons, food: n = 5 mice, 1623 neurons. n. Single mouse example of the population response similarity (Pearson correlation) between the start of the day vs. the end of the same day, as compared to the start of two different days with a similar deficiency level. o. Neuronal population activity comparison between the start and end of the same day, and between the start of two different days with similar deficiency levels. Similarity was significantly higher between two deficiency states (start of different days) than between the start and end of the same day (high vs. low deficiency state). *Top*: population response similarity (Pearson correlation; Water: **p = 0.002, n = 10; Food: *p = 0.023, n = 5). *Bottom*: Population correlation slope (Water: **p = 0.003, n = 10 mice; Food: *p = 0.013, n = 5 mice). 381± 174 and 325±131 neurons per mouse, for water and food, respectively; mean±SEM.

**Ext. Data Fig. 3:**
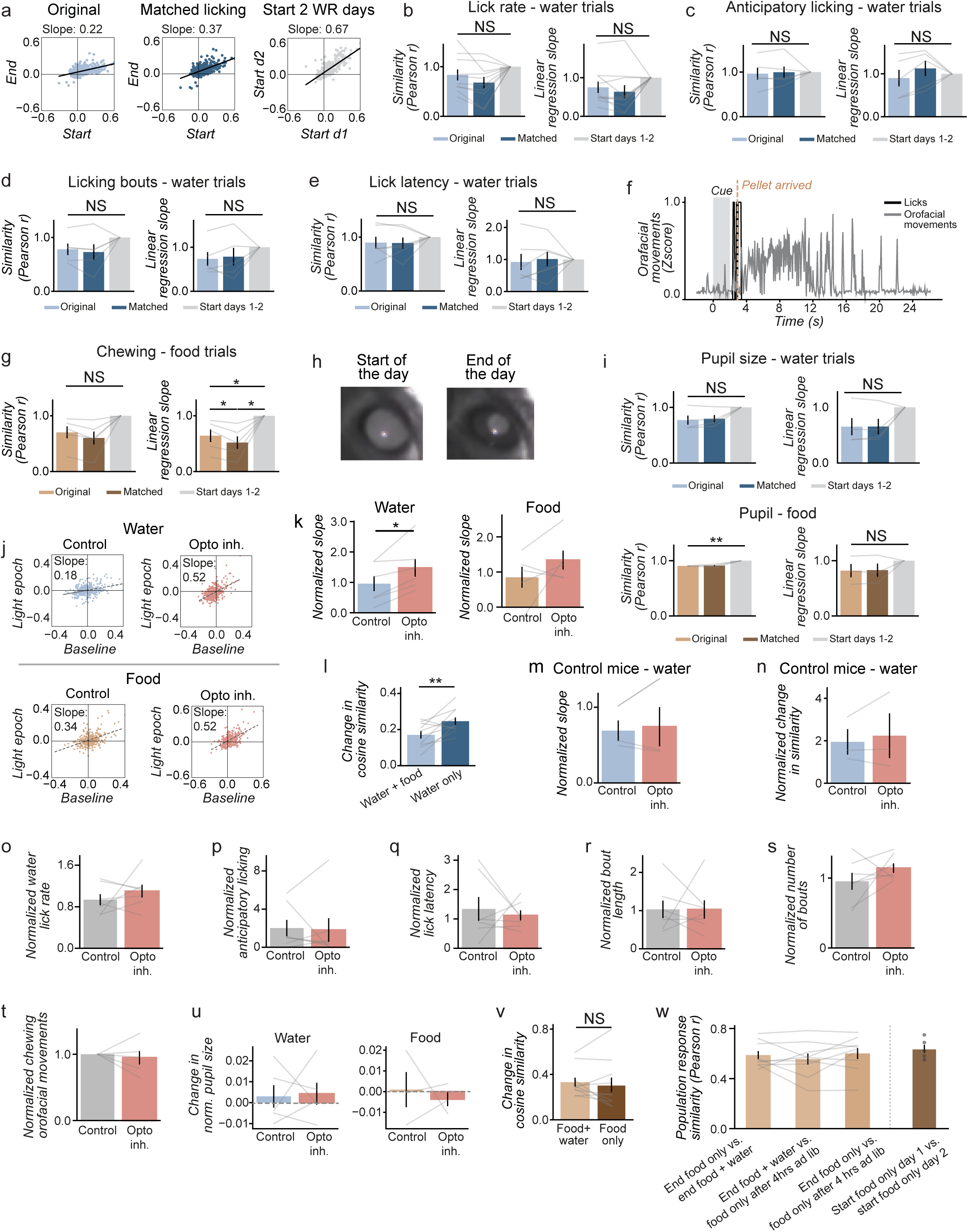
Additional analyses and controls for the effects of VPM/Lpc→InsCtx inhibition on behavior and InsCtx ongoing activity. a. Single mouse example for water responses with decreasing physiological need while controlling for lick rate. *Left*: original population response similarity between the start of the day vs. the end of the same water restricted day. *Middle*: same as left, but only using trials for the start and the end of the day with similar lick rates in both conditions. *Right*: comparing the start of two different days. n = 496 neurons. b–e,g,i. Same analysis as in ‘a’, comparing population response similarity between ‘original’, ‘matched trials’, and ‘Start days 1–2’ conditions, but performed for different behavioral variables used to match trials between the start and end of the session. *Left*: population response similarity (Pearson correlation). *Right*: correlation slope values. Note that matching these behavioral variables did not substantially increase the correlation coefficients and slopes, and they remained lower than across day comparison with a similar need level (start of the day), as well as similar or lower than the original unmatched comparison. b. Matching water responses by lick rate during the reward consumption. All comparisons were not significant. n = 10 mice, 381±174 neurons per mouse; mean±SEM. c. Matching water responses by anticipatory licking during the cue (2 sec). All comparisons were not significant. n = 5 mice, 312±113 neurons per mouse; mean±SEM. d. Matching water responses by number of lick bouts. All comparisons were not significant. n = 5 mice, 346±146neurons per mouse; mean±SEM. e. Matching water responses by first lick latency. All comparisons were not significant. n = 6 mice, 382±188 neurons per mouse; mean±SEM. f. Example quantification of chewing orofacial movements using facemap. g. Matching food responses by orofacial movements. Slope Matched vs. Original *p=0.037, Matched vs. Start-Two-Days *p=0.037, Original vs. Start-Two-Days *p=0.045. n = 5 mice, 325±131 neurons per mouse; mean±SEM. Note that matching trials for chewing significantly decreased the slope rather than increasing it, further supporting the conclusion that the within-day changes are not driven by chewing. h. Example pupil images at the start (*left*) versus the end (*right*) of a session. i. Matching by pupil size. For water, all comparisons were not significant (*top*. n = 4 mice, 407±226 neurons per mouse); and food, correlations Original vs. Start-Two-Days **p=0.002 (*bottom*. n = 3 mice, 318±161 neurons per mouse); mean±SEM. This significant change indicates that the beginnings of two days are more similar to each other than the start and end of each day, regardless of matching pupil size. j. All neurons from one mouse, comparing responses to water (*top*) and food (*bottom*) before and after the optogenetic inhibition epoch. Note an increase in correlation slope, indicating an overall stabilization of responses, blocking the effects of decreasing physiological need. Water: 425 neurons, food: 306 neurons. k. Quantification of the long-term effects of optogenetic inhibition of VPM/Lpc→InsCtx axons. Correlation slope values across all mice of population activity before and after optogenetic inhibition. The inhibition caused a significant increase in the slope for water (6/6 mice, *p = 0.01) and a tendency for an increase in food (3/5 mice, p = 0.13). 406±192 and 325±131 neurons per mouse, for water and food, respectively; mean±SEM. l. Change in cosine similarity of water responses across water restricted mice, between a day with both water and food available, and a day on which only water were available. When food is present, mice remain thirsty, as food intake increases thirst, resulting in a smaller change in response similarity (**p=0.008). This suggests that the change observed on day with only available water is not simply due to the passage of time but reflects a shift in physiological state. n = 10 mice, 349±175 neurons per mouse; mean±SEM. m. Control for long-term stabilization of the correlation-slope effect during optogenetic inhibition. Mice without eOPN3 underwent the same optogenetic inhibition protocol. Summary of the change in population response similarity before versus after the illumination period, normalized to control day 2. No significant effect (p = 0.38). n = 3 mice, 401 ± 101 neurons per mouse; mean ± SEM. n. Control for long-term stabilization of the population cosine similarity effect during optogenetic inhibition. Mice without eOPN3 underwent the same optogenetic inhibition protocol. Summary of the change in population response similarity before versus after the illumination period, normalized to control day 2. No significant effect (p = 0.3). n = 3 mice, 401 ± 101 neurons per mouse; mean ± SEM. o-u. Comparison of behavioral variables between control day 1 and the optogenetic inhibition day during the illumination window, normalized to control day 2, for water and food trials. Testing whether behavior itself differed between conditions. No significant difference was found between control and opto for any variable. Lick rate during consumption of water trials (o, n = 7 mice); anticipatory licking during the 2 sec cue of water trials (p, n = 7 mice); licking duration for water trials (q, n = 7 mice); lick bouts length for water trials (r, n = 7 mice); number of consumption bouts (s, n = 7 mice); orofacial movements during food consumption of food trials (t, n = 5 mice); pupil size during consumption of water and food (u, n = 5 mice and n =3 mice, respectively). All values are mean±SEM. v. Summary of the same mice change in normalized cosine similarity between the start and end of the session, comparing food-restricted day with food only available versus food-restricted day with both food and water available. No significant difference was found between the two conditions, indicating that food responses change similarly with decreasing hunger regardless of water availability. n = 10 mice; 349±175 neurons per mouse; mean ± SEM. w. Population response similarity (Pearson correlation) for consumption responses, comparing three conditions: (1) end of session in food-restricted mice with both food and water available. (2) end of session in food-restricted mice with only food available. (3) after 4 hours of free access to food and water in the home cage. No significant difference was found between conditions. n = 9 mice, 344±184 neurons per mouse; mean ± SEM. “Start 2 hungry”: correlation between the start of two similar food-restricted (hungry) days, shown for reference. n = 5 mice, 325±131 neurons per mouse; mean ± SEM

**Ext. Data Fig. 4:**
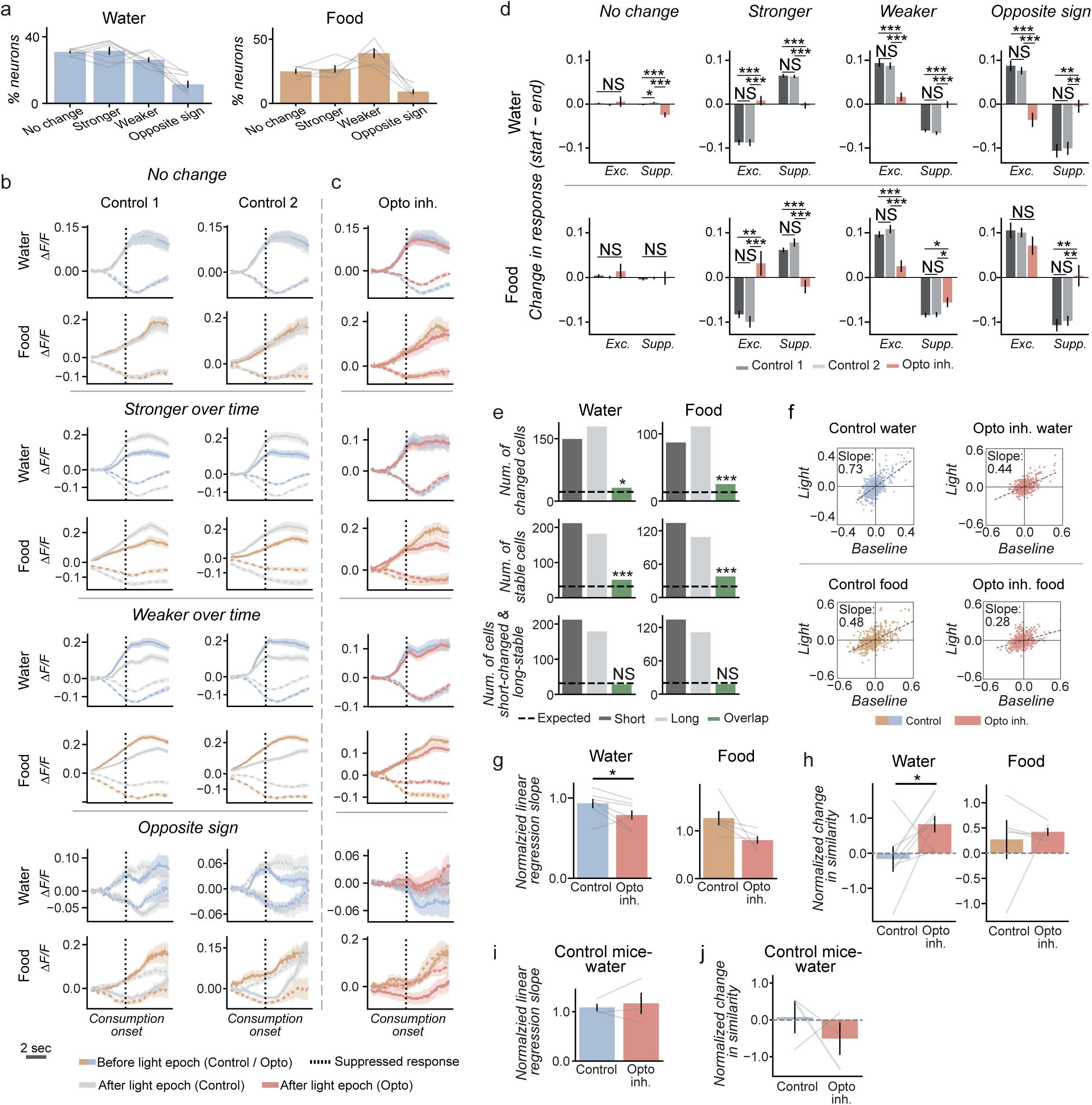
Analyses and controls for the short-term effects of VPM/Lpc→InsCtx inhibition on InsCtx. a. Percent of responsive neurons classified into each change-type group (no change, stronger, weaker, opposite sign), comparing responses just before versus just after the onset of the first optogenetic inhibition epoch, for water (*left*, n = 7 mice, 259±114 neurons per mouse) and food (*right*, n = 5 mice, 204±80 neurons per mouse). mean ± SEM. b. Short-term change in InsCtx average group responses, comparing just before versus just after the onset of the first optogenetic inhibition epoch, across two control days (Control 1, Control 2) for the 4 groups (no change, stronger, weaker, opposite sign), pooled across all mice. Only responsive neurons showing a consistent change across both control days were included. Solid lines: excitatory responses; dashed lines: suppressed responses. Blue (water responses in water-restricted mice. n = 7 mice, no change-174 neurons, stronger-154 neurons, weaker-139 neurons, opposite sign-29 neurons) and brown (food responses in food-restricted mice. n = 5 mice, no change-56 neurons, stronger-60 neurons, weaker-143 neurons, opposite sign-24 neurons): responses ∼25 min into the session. Gray: responses after 10 minutes of consumption. Dotted vertical line: consumption onset. c. Same neurons as in ‘b’, but during the day with optogenetic inhibition of VPM/Lpc→InsCtx axons. Blue (water) and brown (food): responses ∼25 min into the session, before optogenetic inhibition. Red: responses after the start of optogenetic inhibition. Solid lines: excitatory responses, dashed lines: suppressed responses. Dotted vertical line: consumption onset. d. Short-term effects of optogenetic inhibition: quantification of the change in individual neuron response magnitude for each neuron type, pooled across all mice (same as ‘b’; mean ± SEM). For each neuron, the response was measured as the mean over the 2 seconds following reward consumption onset, and the difference between responses just before and just after the onset of the first optogenetic inhibition epoch was calculated, for each of the three days (Control 1 - C1, Control 2 - C2, Opto). Neurons were grouped into four categories based on their natural change with decreasing physiological need across control days: No change, Stronger, Weaker, and opposite sign, shown separately for excitatory (Exc.) and suppressed (Supp.) responses. Note that in all groups showing a natural change across control days, optogenetic inhibition attenuates or completely eliminates this change. *Top*: water responses. **No change**: Exc.- C1 vs. opto: ***p<0.001, C2 vs. opto: ***p<0.001, C1 vs. C2: *p=0.014, Supp.- no significant changes. **Stronger**: Exc.- C1 vs. opto: ***p<0.001, C2 vs. opto: ***p<0.001, Supp.- C1 vs. opto: ***p<0.001, C2 vs. opto: ***p<0.001. **Weaker**: Exc.- C1 vs. opto: ***p=<0.001, C2 vs. opto: ***p<0.001, Supp.- C1 vs. opto: ***p<0.001, C2 vs. opto: ***p<0.001. **Opposite sign**: Exc.- C1 vs. opto: ***p<0.001, C2 vs. opto: ***p<0.001, Supp.- C1 vs. opto: **p=0.001, C2 vs. opto: **p=0.001). *Bottom*: food responses. **No change**- no significant changes. **Stronger**: Exc. C1 vs. opto: **p=0.001, C2 vs. opto: ***p<0.001, Supp.- C1 vs. opto: ***p<0.001, C2 vs. opto: ***p<0.001. **Weaker**: Exc.- C1 vs. opto: ***p=<0.001, C2 vs. opto: ***p<0.001, Supp.- C1 vs. opto: *p=0.021, C2 vs. opto: *p=0.014. **Opposite sign**: Exc.- no significant changes, Supp.- C1 vs. opto: **p=0.002, C2 vs. opto: **p=0.001). When not mentioned, no significant difference was found between Control 1 and Control 2 in any group. e. Overlap between neurons with short-term and long-term effects of optogenetic inhibition, for water (*left*) and food (*right*). In each group, we selected the top 30% of neurons for comparison. Bars show the number of neurons classified in the long-term population (“Long”), short-term population (“Short”), and their overlap (“Overlap”). Dashed line: overlap expected by chance (hypergeometric test). *Top*: “stable” neurons (Water: long-179 neurons, short-149, *p=0.013, Food: long-111, short-87, ***p<0.001). *Middle*: “changed” neurons (Water: long-182 neurons, short-212, ***p<0.001, Food: long-109, short-134, ***p<0.001). *Bottom*: long term stable vs. short term changed Water: long stable-179 neurons, short changed-212, *p=0.8, Food: long stable-111, neurons, short changed-134, *p=0.76). f. All neurons from one mouse, comparing responses to water (*top*, n = 735 neurons) and food (*bottom*, n = 520 neurons) before and during the optogenetic inhibition epoch in the immediate period after inhibition onset. Note a decrease in correlation slope, indicating an overall decrease in response. g. Quantification of the immediate effects of optogenetic inhibition of VPM/Lpc→InsCtx axons. Correlation slope values across all mice of population activity before and after optogenetic inhibition onset. The inhibition caused a significant reduction in the slope for water (7/7 mice, *p = 0.03) and a tendency for reduction in food (4/5 mice, p = 0.055). Water: n = 7 (372±196 neurons per mouse; mean±SEM); Food: n = 5 (325±131 neurons per mouse; mean±SEM). h. Summary of the immediate change in population response similarity due to optogenetic inhibition. Change in similarity during the illumination period, 5-7 trials after light onset and 5-7 trials before light offset (significantly lower delta on the optogenetic day for water responses, *p = 0.047). *Water*: n = 7, 372±196 neurons per mouse; mean±SEM. *Food* n = 5, 325±131 neurons per mouse; mean±SEM. i. Control for immediate effect on correlation slope values using mice without eOPN3, comparing before and after optogenetic inhibition onset (p=0.35). n = 3 mice, 401±101 neurons per mouse; mean±SEM. j. Control for immediate effect of optogenetic inhibition on normalized cosine similarity values using mice without eOPN3 expression, comparing before and after optogenetic inhibition onset (p=0.28). n = 3 mice, 401±101 neurons per mouse; mean±SEM.

**Ext. Data Fig. 5:**
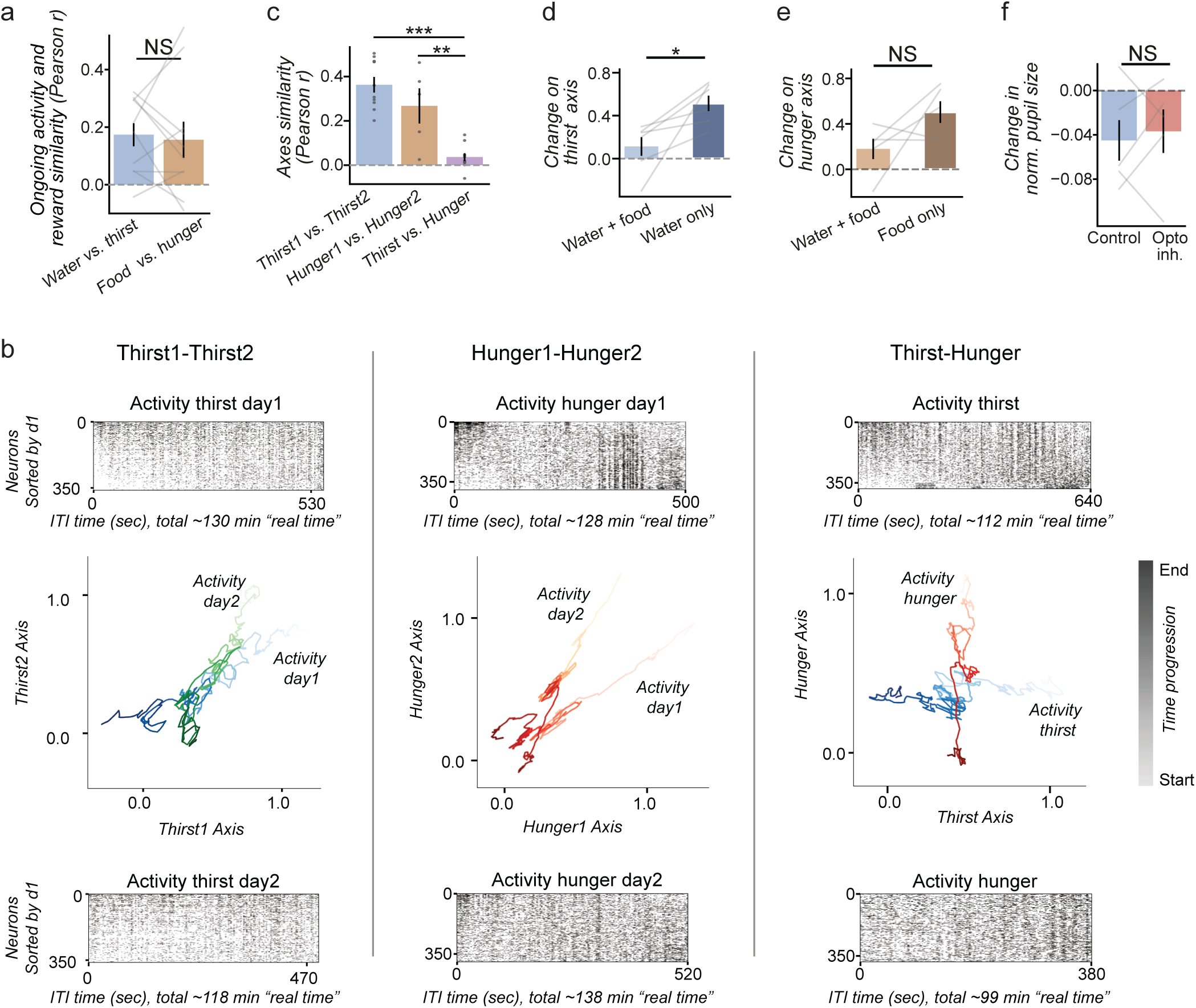
Additional analyses and controls for the long-term effects of VPM/Lpc→InsCtx inhibition on InsCtx ongoing activity. a. No consistent correlation between the change in the amplitude of the reward response and the change in the ongoing activity between the start and the end of the day. n = 10 mice, 349±175 neurons per mouse, mean±SEM. b. Example of ongoing activity changes along different physiological need state axes (same as Fig. 3e but also showing population activity heatmaps). Comparison of two days of thirst state (*left*), two days of hunger state (*middle*), and one day of thirst vs. one day of hunger (*right*). *Top*: Heatmap showing ongoing population activity on the first day (day1), sorted by its change in activity with decreasing physiological need. *Bottom*: Same as the top but for the second day (day2), sorted by Day 1’s activity change. Note the similarity in heatmaps when the physiological need is the same (left and middle panels). *Middle*: 2D plot of ongoing activity from each day projected onto the axes of both days. A value of 1 indicates the restricted state, while a value of 0 indicates the sated state. Color changes represent time (light color: start, dark color: end). c. Similarity (Pearson correlation) between all mice between pairs of physiological need state axes (correlation between population vectors). Comparison of two days of thirst axes, two days of hunger axes, and one day of thirst vs. one day of hunger. Thirst vs. Hunger-Thirst: ***p<0.001, Hunger vs. Hunger-Thirst **p=0.002. n = 10 mice for thirst-thirst (381±174 neurons per mouse, mean±SEM), n = 5 mice for hunger-hunger (325±131 neurons per mouse, mean±SEM), and n = 10 mice for thirst-hunger (349±175 neurons per mouse, mean±SEM). d. Change along the thirst axis (start vs. end of day) in the same mice (n = 6) on two water-restriction days: one with both water and food available (water + food), and one with only water available (water only). In the water + food day, only inter-trial intervals (ITIs) following active water consumption were included, and were retained across subsequent neutral trials until any other trial type occurred. The thirst axis was defined using data from a separate, subsequent water-only day (see Methods). Neurons per mouse: water + food, 295 ± 147; water only, 417 ± 175. *p = 0.027, paired t-test; mean ± SEM. e. Change along the hunger axis (start vs. end of day) in the same mice (n = 5) on two food-restriction days: one with both water and food available (water + food), and one with only food available (food only). In the water + food day, only ITIs following active food consumption (including food taste omission or problem trials) were included, and were retained across subsequent neutral trials until any other trial type occurred. The hunger axis was defined using data from a separate, subsequent food-only day (see Methods). Neurons per mouse: water + food, 332 ± 134; food only, 325 ± 131. p = 0.15, paired t-test; mean ± SEM. f. Change in pupil size during inter-trial intervals (ITIs), used for ongoing activity, across a water-restriction day with water consumption. Bars show the change in pupil size (start vs. end of day). n = 5 mice.

**Ext. Data Fig. 6:**
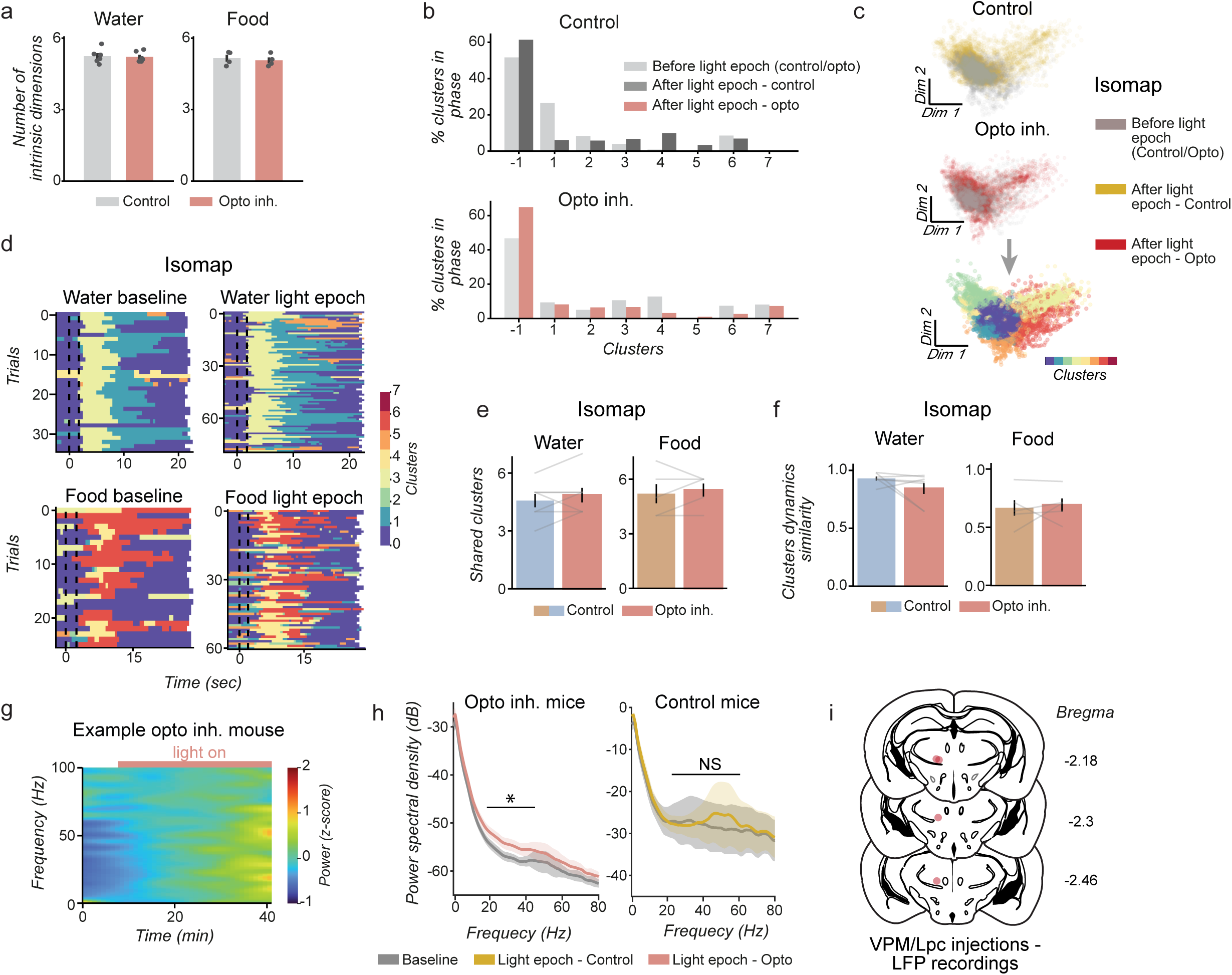
Additional analyses and controls for the long-term effects of VPM/Lpc→InsCtx inhibition on InsCtx population activity manifold. a. Estimated intrinsic dimensionality of neural population activity for water (n = 7 mice) and food (n = 5 mice) trials, control versus optogenetic inhibition. Dimensionality estimates were similar between conditions (approximately 5–6 dimensions in all cases). b. Example from one mouse for manifold cluster distribution, related to the shared clusters analysis in Fig. 4c, between baseline and light epochs for control day and day with optogenetic inhibition. c-f. Structure and dynamics of population activity, visualized using Isomap as an alternative method for dimensionality reduction. c. Similarly to Fig. 4a but now using isomap: example of the lack of effect of optogenetic inhibition on the structure of the InsCtx activity manifold, shown via representative planes of dimensionality-reduced neuronal activity. Colors in the left and middle panels indicate different epochs (*gray*: baseline, *red*: optogenetic inhibition epoch, *yellow*: optogenetic inhibition epoch on the control day with no illumination) for control (*top*) and optogenetic inhibition day (*middle*). To quantify their similarity, the manifold was clustered into distinct activity clusters (see Methods; *bottom*, same data as in the middle panel). d. Example from a single mouse showing manifold cluster sequence dynamics using isomap during water and food trials in the baseline versus optogenetic inhibition period. Dashed lines indicate cue onsets. Note similar dynamics before and during the optogenetic inhibition epochs, suggesting that it does not alter manifold cluster dynamics. e. Similarly to Fig. 4c, but now using isomap. Quantification of the effect of optogenetic inhibition on the activity manifold structure across mice. The number of shared clusters between the baseline and light epoch did not differ between control and optogenetic inhibition days for either water or food responses (n = 7 mice, 372±196, food: 325±131 neurons per mouse, for water and food, respectively; mean±SEM). f. Quantification of the effect of optogenetic inhibition on manifold cluster sequence dynamics using isomap, quantified by the similarity in distribution of trial clusters across epochs (see Methods). No significant difference in cluster dynamics similarity between control day 2 and the optogenetic inhibition day for both water and food trials (n = 7 mice, 372±196, food: 325±131 neurons per mouse, for water and food, respectively; mean±SEM). g. Example spectrogram of InsCtx LFP activity from one mouse before and during VPM/Lpc→InsCtx optogenetic inhibition. h. Change in power spectral density during illumination in optogenetic inhibition mice (left) and in control mice (right), averaged across mice. Gray: baseline, red: optogenetic inhibition. n = 4, mean ± SEM. i. Coronal schematics of histological verification, corresponding to experiments in ‘g’ and,’h’. Unilateral injection sites of AAV5-mCaMKIIa(short)-eOPN3-mScarlet in VPM/Lpc (Bregma −2.18 to −2.46 mm).

**Ext. Data Fig. 7:**
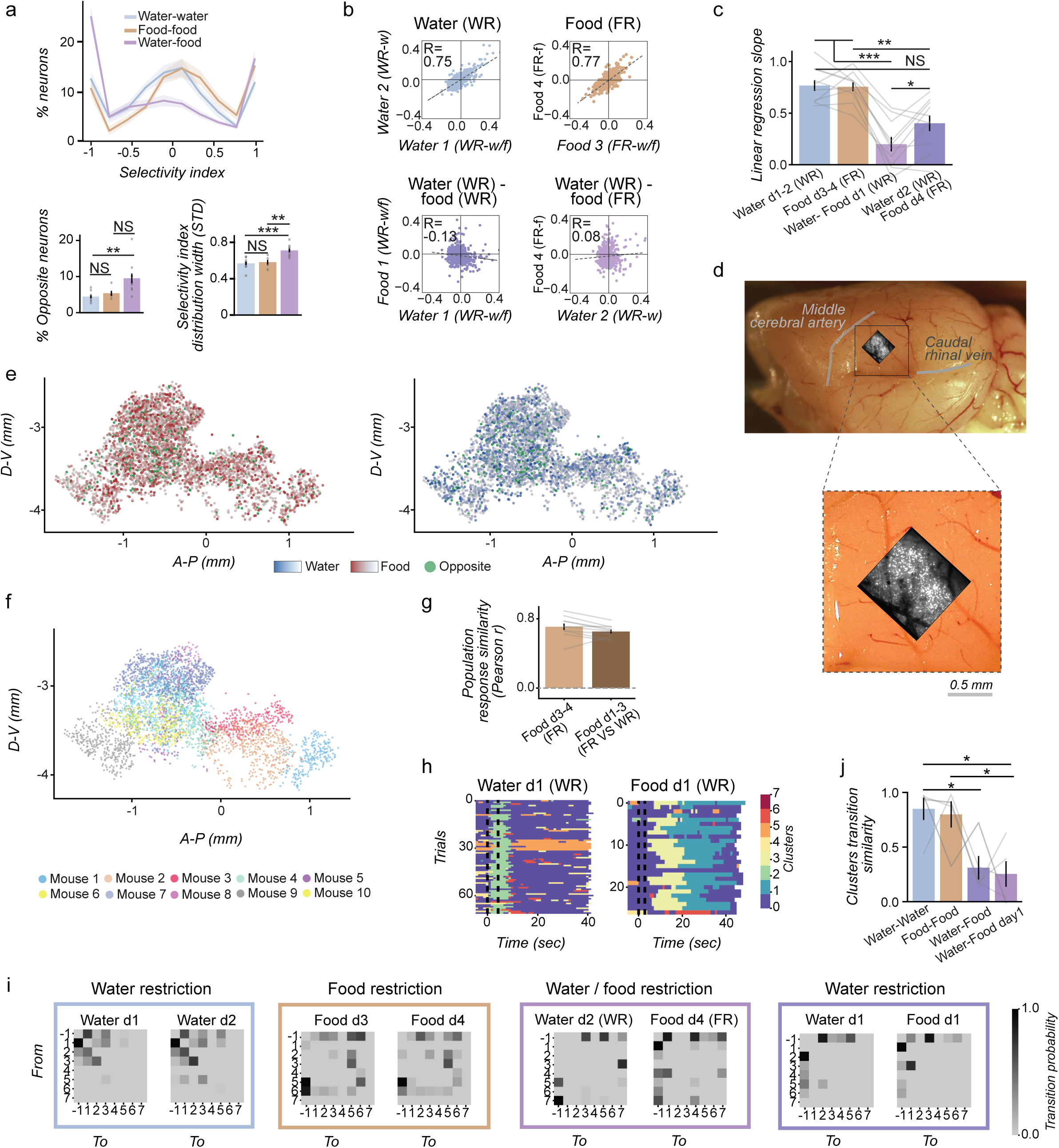
Further analyses and controls for the comparisons between food and water responses in InsCtx. a. *Top*: Selectivity of single neurons for water vs. food rewards. Selectivity index distribution (mean ± SEM across mice) for two days of water consumption during water restriction (water vs. water, blue), two days of food consumption during food restriction (food vs. food, brown) and water while water restricted vs. food while food restricted (purple). Values of +1 indicate preference for the first day, while values of -1 indicate preference for the second day. *Bottom left*: Percent of neurons with opposite responses (excitation/suppression) between days, thereby precluding their inclusion in the selectivity index analysis. Note a significant two-fold increase in sign-switching neurons in the water–food comparison (Water vs. Water-Food: p = 0.002; Food vs. Water-Food: p = 0.026). *Bottom right*: Selectivity index distribution width (assessed by STD) across days. Note higher distribution width in the water–food condition (Water vs. Water-Food: p < 0.001; Food vs. Water-Food: p = 0.002). Water: n = 10 (381±174 neurons per mouse; mean±SEM); Food: n = 5 (325±131 neurons per mouse; mean±SEM). b. Example population response similarity (correlation) during consumption of water (w) and food (f) rewards during the relevant physiological need (WR: water restriction, FR: food restriction). c. Population response similarity correlation slope during water and food consumption (correlation coefficients appear in Fig. 1k). Note significantly higher values within same-reward consumption (water–water, food–food) compared to opposite-reward consumption (water–food). Water-Food d1 vs. Water / Food ***p < 0.001, Water-Food d2-d4 vs. Water p=0.055, Water-Food d2-d4 vs. Food **p=0.003, Water-Food d2-d4 vs. Water-Food d1 *p=0.042, n = 10, 349±175 neurons per mouse; mean±SEM. d. Example determination of two-photon imaging field-of-view location on post-mortem wholemount brains. *Top*: Wholemount brain image overlayed with two-photon imaging field-of-view. *Bottom*: zoomed-in on top panel (scale bar: 500 µm). e. Topography of water-food responses. Locations relative to Bregma of all imaged neurons (dots) across mice, colored by response magnitude to food (red gradient, *top*) or water (blue gradient, *bottom*). Green: Neurons with opposite responses to food/water (excitation/suppression, see Fig 4B). No apparent topographical organization. n = 10 mice, 3492 neurons. f. Anatomical location of recorded neurons along the dorsoventral (D-V) and anteroposterior (A-P) axes, as imaged through the microprism, corresponding to water vs. food experiments in Fig.5. Each point represents one neuron, colors indicate individual mouse. n = 10 mice, 3492 neurons. g. Population response similarity (Pearson correlation) for the same mice between food responses on days 3 and 4 (both food-restricted), compared to similarity between food responses on days 1 (water-restricted) and 3 (food-restricted) with both food and water available. Similar correlation values between the two comparisons indicate that water restriction does not dramatically affect food response representation. n = 10 mice, 349±175 neurons per mouse; mean±SEM. h. Example manifold cluster dynamics in a single mouse during thirst (day 1) when both water and food rewards are available and consumed. *Left*: Consumed water rewards. *Right*: Consumed food rewards. Note that for each comparison, cluster indices (colors) are randomly assigned during the clustering procedure. Dashed lines: cue. Note consistent yet distinct cluster dynamics for food and water within the same day. i. Example from a single mouse of clusters transition probabilities between pairs of days. *T*wo days of water responses during thirst, *t*wo days of food responses during hunger, water rewards during thirst versus food rewards during hunger (days 2 and 4), water versus food trials during thirst (day 1). The probability of each cluster to remain in the same cluster is not shown. Transition probabilities are normalized to the highest probability within day. j. Quantification of similarity in clusters transition probabilities across mice for pairs of days. Similarity was estimated using Pearson correlation (see Methods). Days with the same need and available rewards showed significantly higher similarity than water versus food within the same need or across different needs (Water vs. Water-Food d2-4 *p=0.046, Food vs. Water-Food d2-4 p=0.14, Water vs. Water-Food d1 *p=0.046, Food vs. Water-Food d1 **p=0.024). n = 5 mice, 332±134 neurons per mouse; mean±SEM.

**Ext. Data Fig. 8:**
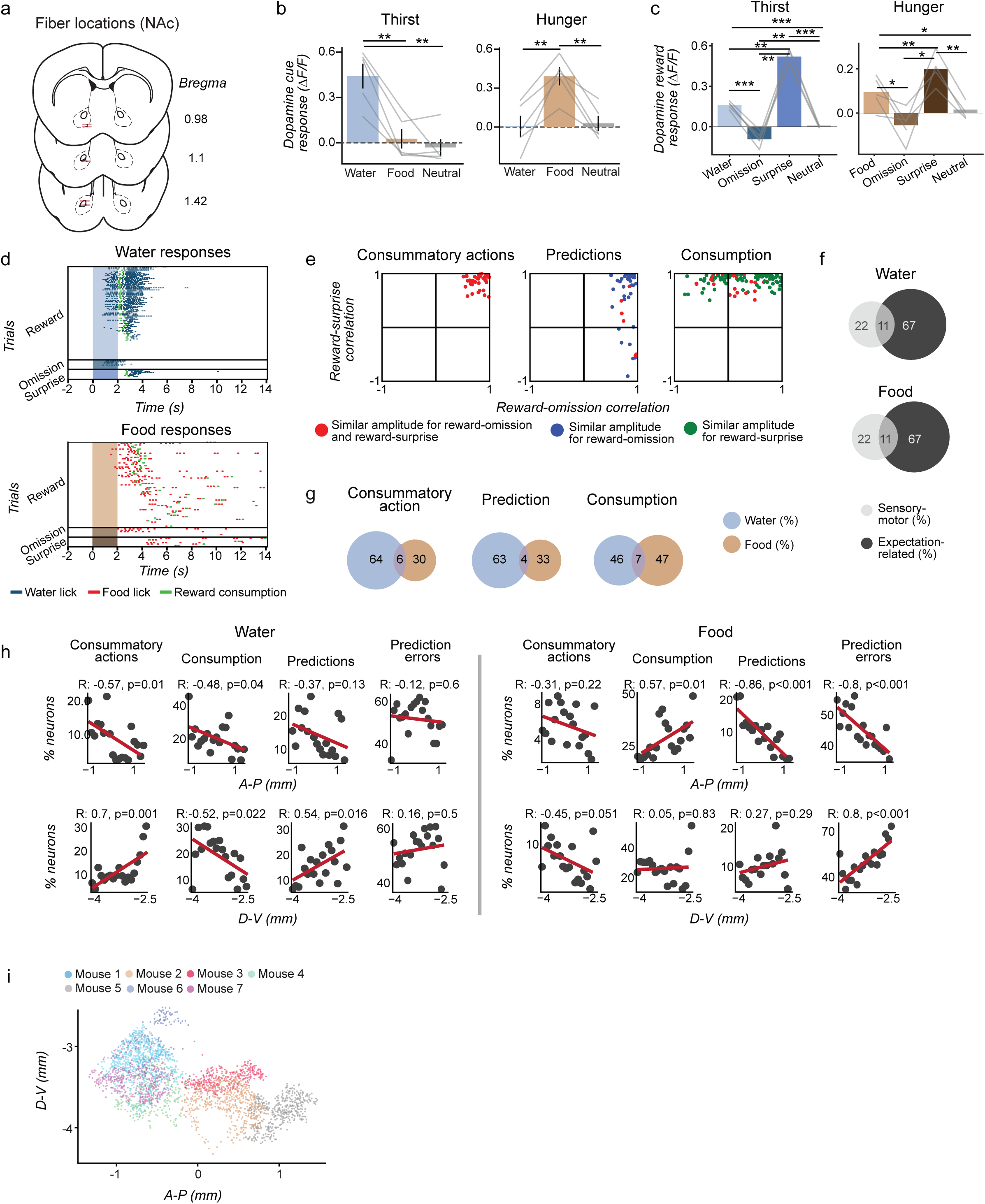
Further analyses and controls for water and food responses during dopamine fiber photometry and response type classifications. a. Coronal schematics of histological verification, corresponding to experiments in Fig. 5h-j. Unilateral fiber location in the nucleus accumbens (Bregma 0.98 to 1.42 mm). n = 5. b. Mean NAc dopamine responses for food, water and neutral cue, during thirst (*left*) and hunger (*right*). Response to the relevant reward is significantly higher in the relevant need (*Thirst*: Water vs. Food **p=0.004, Water vs. Neutral **p=0.004, *Hunger*: Water vs. Food **p=0.01, Food vs. Neutral **p=0.01). n=5 mice. Note that when there was a mismatch between physiological need and reward (i.e., water during food-restriction, and vice versa) dopamine food or water cue responses on consumption trials became on average similar to the neutral cue responses, despite the fact that mice chose to consume food and water on those trials. Thus, although dry food is often considered to be aversive during dehydration, mice in our task chose to consume it during water deprivation and dopamine responses to dry food during water restriction were similar to water during food restriction. n = 5 mice. c. Comparison of NAc dopamine responses to reward delivery, reward omission, surprise reward and neutral trials, for food and water responses. Omissions and surprises exhibit the expected negative and positive prediction error responses (*Thirst*: Water vs. Omission, Neutral ***p<0.001, Water vs. Surprise **p=0.00 3, Omission vs. Surprise **p=0.001, Neutral vs. Surprise ***p<0.001, Neutral vs. Omission **p=0.005. *Hunger:* Food vs. Omission *p=0.038, Food vs. Surprise **p=0.002, Food vs. Neutral *p=0.021, Omission vs. Surprise *p=0.038, Neutral vs. Surprise **p=0.009, Neutral vs. Omission p=0.055). n = 5 mice. d. Example licking behavior from one mouse to reward delivery, reward omission and surprise reward trials during water restriction (*top*) and food restriction (*bottom*). Blue ticks: licking to water spout. Red ticks: licking to food cup. Green ticks: reward consumption. Colored rectangles: cues. Note that in surprise trials the cue is the neutral cue and that there is therefore no anticipatory licking. We therefore optimally aligned responses for maximum response similarity (see Ext. Data Fig 9f,g, and see Methods). e. Demonstration of response type categorization. Example scatter plots from one mouse of the correlations between rewards and omissions vs. rewards and surprises, shown separately for ‘consummatory action’, ‘prediction’, and ‘consumption’ neurons. Each dot represents a neuron and is color-coded based on response amplitude similarity. *Red*: all three responses have similar amplitudes, *blue*: reward and omission responses have similar amplitudes, green: reward and surprise responses have similar amplitudes. f. Overlap between “sensory-motor” and “expectation-related” neurons across mice for water (*top*) and food (*bottom*). Numbers indicate average percentage across mice. n = 7 mice, 269±126, and 233±81 neurons per mouse, for water and food, respectively; mean±SEM. g. Overlap between food and water responses across in response types. Numbers indicate average percentage across mice. n = 7 mice, 131±52, and 99±27 neurons per mouse, for water and food, respectively; mean±SEM. h. Topographical distribution, quantified by the percentage of all imaged neurons per spatial bin relative to bregma for each main response type (consummatory actions, consumption, prediction and prediction errors). *Left*: anterior-posterior, *right*: dorsal-ventral. Neurons pooled across 7 mice: *Water*: consummatory actions - 192, consumption - 422, predictions - 300 and prediction errors - 1381. *Food*: consummatory actions - 100, consumption - 432, predictions - 164 and prediction errors - 1211. i. Anatomical location of recorded neurons along the dorsoventral (D-V) and anteroposterior (A-P) axes, as imaged through the microprism, corresponding to cell types experiments in Fig.6. Each point represents one neuron, colors indicate individual mouse. n = 7 mice, 2553 neurons.

**Ext. Data Fig. 9:**
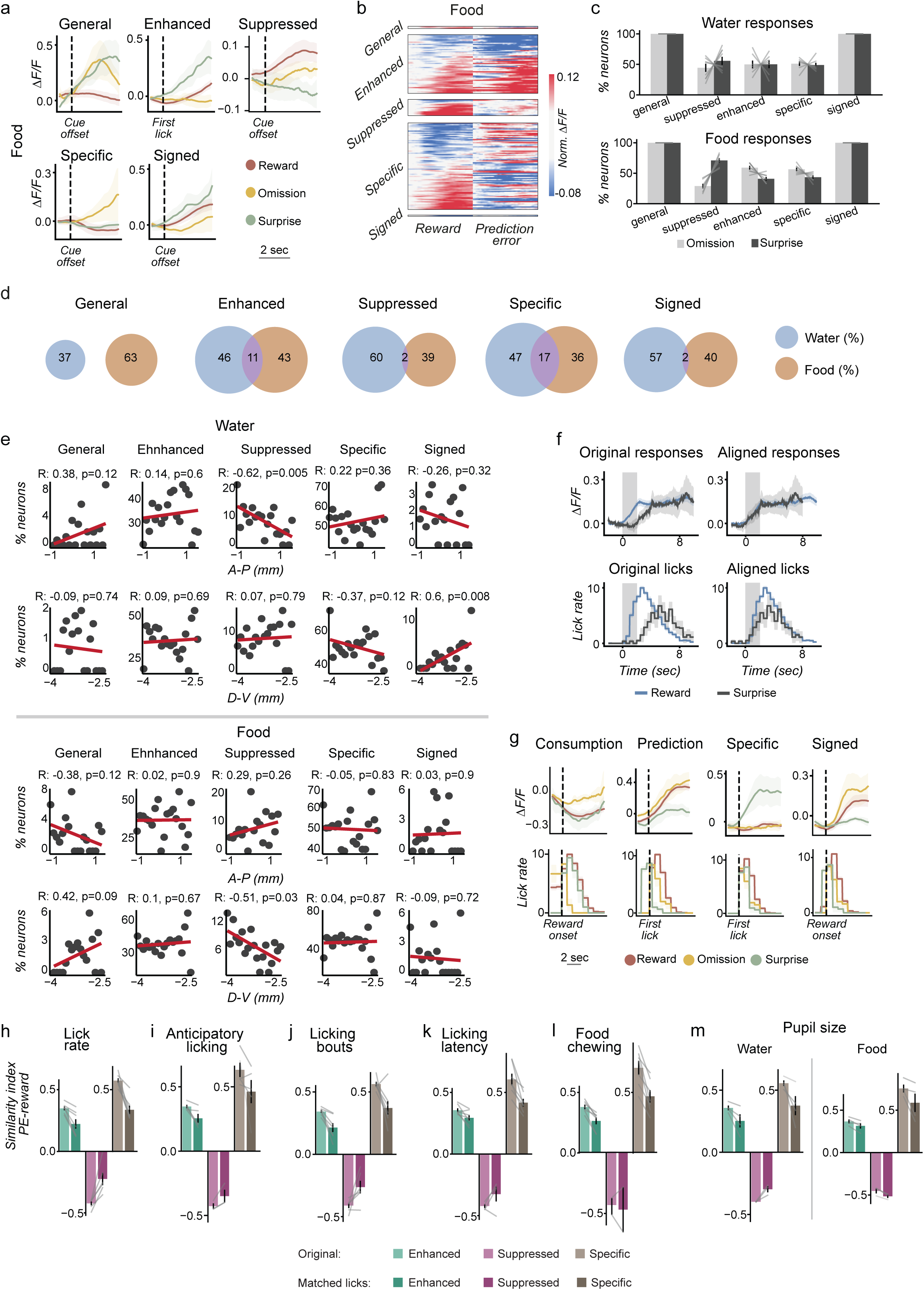
Additional analyses and control for prediction errors classification. a. Example food responses of classified prediction error response types. Neurons were categorized as “general” (similar omission and surprise responses), “enhanced” (similar to reward but stronger response), “suppressed” (similar to reward but weaker response), “specific” (distinct from reward), or “signed” (omission and surprise both resemble the reward response, with one stronger and one weaker). Dashed lines indicate response alignment: reward delivery, or cue offset, or lick onset/offset. b. Example of single-mouse prediction error responses for food. Heatmap showing responses of individual neurons from one mouse to reward delivery, and either reward omission or surprise reward trials, sorted by prediction-error response classification. c. Percent of omission or surprise trials that evoked each type of prediction error response. n = 7 mice, 197±96, and 173±67 neurons per mouse, for water and food, respectively; mean±SEM. d. Overlap between food and water responses across in prediction error types. Numbers indicate average percentage across mice. n = 7 mice, 197±96, and 173±67 neurons per mouse, for water and food, respectively; mean±SEM. e. Topographical distribution quantified as the percent of all imaged neurons relative to the bregma for each prediction error response types. *Top*: anterior-posterior, *Bottom*: dorsal-ventral. Pooled neurons across 7 mice: prediction error response type neuron numbers: *Water*: general - 15, enhanced - 522, suppressed - 138, specific - 760, signed - 28. *Food*: general - 26, enhanced - 489, suppressed - 91, specific - 630, signed - 20. f. Example of single neuron alignment of surprise response to reward response. Original response (left) and after alignment (right). g. Example single neuron responses from different classified response types to water (*top*), shown with lick rate for each trial type (*bottom*), demonstrating that distinct responses to the different conditions cannot be explained by differential licking. Dashed lines indicate response alignment: reward delivery, cue offset, and lick onset/offset. h-m. Verification that enhanced, suppressed, and specific prediction error responses are not due to differences in behavior between trial types (see Methods). Quantification across mice for reward and prediction error (PE) responses, using trials matched for different behavioral variables across conditions. Values represent the mean modulation index across all neurons per group. Values of +1 indicate a stronger response during prediction error (enhanced), whereas values of –1 indicate a stronger response during reward (suppressed). All values are mean±SEM. Importantly, even though the similarity index is affected in some cases, it remains positive for enhanced and specific prediction errors, and negative for suppressed prediction errors, after controlling for behavior. This shows that the classification of these responses as prediction errors does not stem from behavioral variables. h. Lick rate during consumption of water trials (enhanced: n = 7 mice, 73±28 neurons per mouse, suppressed: n = 6 mice, 23±16 neurons per mouse, specific: n = 7 mice, 106±72 neurons per mouse). i. Anticipatory licking during the 2 sec cue of water trials (enhanced: n = 5 mice, 27±14 neurons per mouse, suppressed: n = 3 mice, 16±10 neurons per mouse, specific: n = 5 mice, 41±28 neurons per mouse). j. Lick bouts length for water trials (enhanced: n = 7 mice, 67±34 neurons per mouse, suppressed: n = 6 mice, 22±16 neurons per mouse, specific: n = 7 mice, 97±75 neurons per mouse). k. First lick latency for omission prediction errors (enhanced: n = 7 mice, 33±17 neurons per mouse, suppressed: n = 4 mice, 17±8 neurons per mouse, specific: n = 7 mice, 56±41 neurons per mouse). l. Orofacial movements during food consumption of food trials for surprise prediction errors (enhanced: n = 7 mice, 48±15 neurons per mouse, suppressed: n = 2 mice, 8±1 neurons per mouse, specific: n = 7 mice, 37±18 neurons per mouse). m. Pupil size during consumption of water (left) and food (right; water-enhanced: n = 4 mice, 68±32 neurons per mouse, suppressed: n = 3 mice, 21±21 neurons per mouse, specific: n = 4 mice, 109±98 neurons per mouse; Food-enhanced: n = 4 mice, 73±41 neurons per mouse, suppressed: n = 3 mice, 14±1 neurons per mouse, specific: n = 4 mice, 83±54 neurons per mouse).

**Ext. Data Fig 10:**
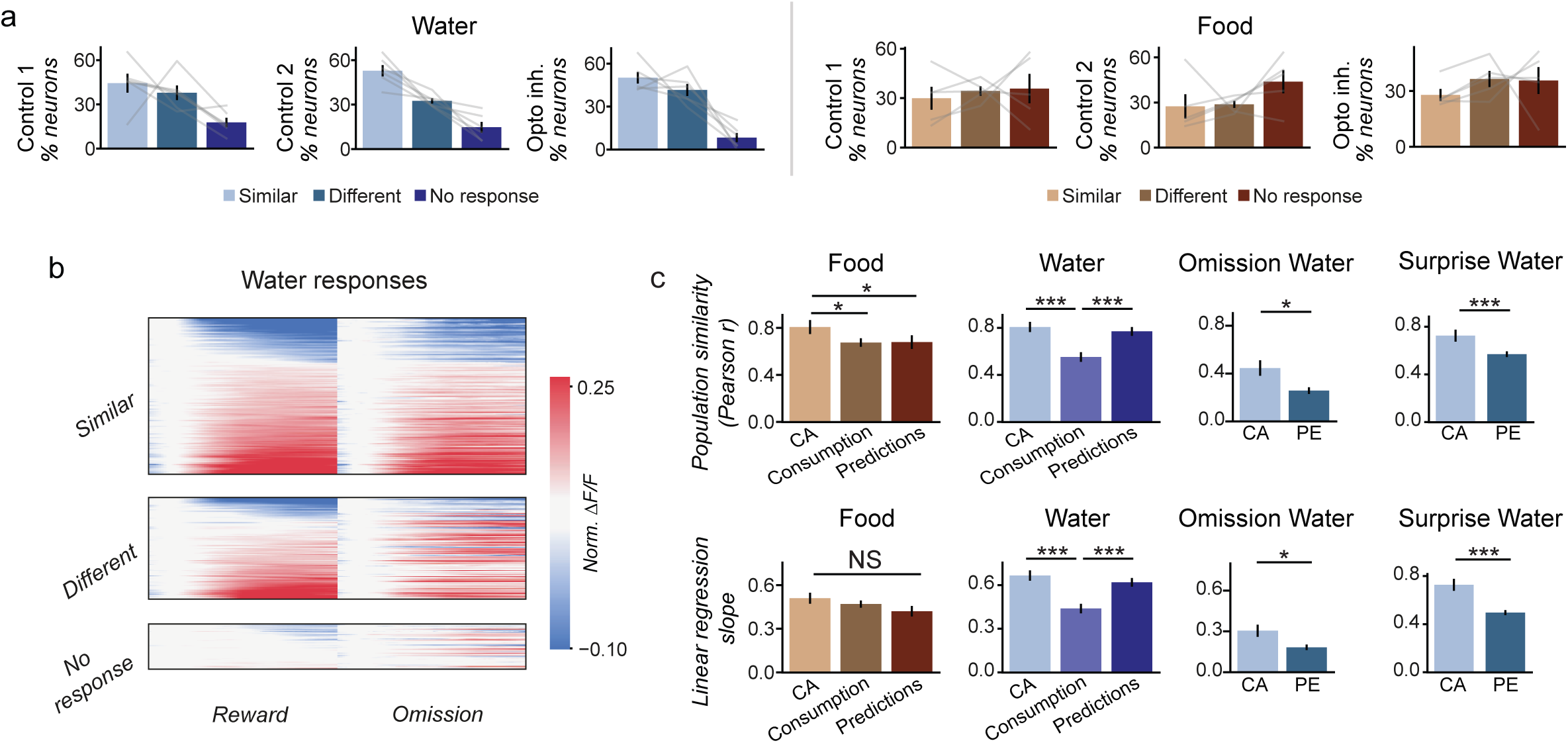
Additional analyses and control for optogenetic inhibition effect on different cells types. a. Summary of reward-omission classification by days for water (*left*) and food (*right*). Water n = 6 mice, 311±139 neurons per mouse; mean±SEM, food n = 5 mice, 325±131 neurons per mouse; mean±SEM. b. Classification of neurons based on responses to reward delivery and reward omission during the optogenetic inhibition experiment. Example heatmap showing the classification of neurons to different response types. Neurons were categorized based on similarity or dissimilarity between their responses to reward delivery and reward omission during the baseline period of each day (before optogenetic inhibition onset). Note that the ‘similar’ group includes ‘consummatory action’ and ‘prediction’, whereas the ‘different’ group includes ‘prediction error’. c. Effects of decreasing physiological need on different response types. *Top:* Population response similarity (Pearson correlation) across all mice comparing neuronal population activity between the start and end of the same day, for different response types. Reward delivery responses for food and water (Food-consummatory actions vs. consumption: *p=0.024, consummatory actions vs. predictions: *p=0.046. Water-consummatory actions vs. consumption: ***p<0.001, consumption vs. prediction errors: ***p<0.001), and reward omission / surprise reward responses for water (Surprise ***p<0.001, Omission *p=0.011). We note that we did not analyze surprise and omitted rewards for food trials because we did not have a sufficient number of such trials to compare the beginning and the end of the session. *Bottom*: correlation slope across all mice comparing neuronal population activity between the start and end of the same day, for different response types. Reward responses for food and water (Food-not significant. Water-consummatory actions vs. consumption: ***p<0.001, predictions vs. consumption: **p=0.001) and omission / surprise responses for water (Surprise ***p<0.001, Omission *p=0.016). n = 7 mice for reward responses (water and food), n = 5 mice for omission / surprise responses. Pooled cells from all mice per group: *Water*: CA: 192, consumption: 422, predictions: 300, omission PE: 933, surprise PE: 1182. *Food*: CA: 100, consumption: 432, predictions: 164. CA: consummatory action, PE: prediction error.

